# Dynamics of CTCF and cohesin mediated chromatin looping revealed by live-cell imaging

**DOI:** 10.1101/2021.12.12.472242

**Authors:** Michele Gabriele, Hugo B. Brandão, Simon Grosse-Holz, Asmita Jha, Gina M. Dailey, Claudia Cattoglio, Tsung-Han S. Hsieh, Leonid Mirny, Christoph Zechner, Anders S. Hansen

## Abstract

Animal genomes are folded into loops and topologically associating domains (TADs) by CTCF and cohesin, but whether these loops are stable or dynamic is unknown. Here, we directly visualize chromatin looping at the *Fbn2* TAD in mouse embryonic stem cells using super-resolution live-cell imaging and quantify looping dynamics by Bayesian inference. Our results are consistent with cohesin-mediated loop extrusion in cells, and with CTCF both stopping and stabilizing cohesin. Surprisingly, the *Fbn2* loop is both rare and dynamic, with a looped fraction of ~3-6.5% and a median loop lifetime of ~10-30 minutes. Instead of a stable loop, our results establish a highly dynamic view of TADs and loops where the *Fbn2* TAD exists predominantly in a partially extruded conformation. This dynamic and quantitative view of TADs may facilitate a mechanistic understanding of their functions.

## INTRODUCTION

Mammalian genomes are folded into loops and domains known as Topologically Associating Domains (TADs) by the proteins CTCF and cohesin (*1*). Mechanistically, cohesin is thought to load on DNA and bidirectionally extrude loops until it is blocked by CTCF such that CTCF establishes TAD boundaries (*2*–*7*). Functionally, CTCF- and cohesin-mediated looping and TADs play critical roles in multiple nuclear processes including regulation of gene expression, somatic recombination, and DNA repair (*8*). For example, TADs are thought to regulate gene expression by increasing the frequency of enhancer-promoter interactions within a TAD, and decreasing enhancer-promoter interactions between TADs (*9*). However, to understand how TADs and loops are formed and maintained, and how they function, it is necessary to understand whether TADs are stable or dynamic structures and to quantify the dynamics and lifetime of CTCF/cohesin-mediated loops.

Though recent advances in single-cell genomics and fixed-cell imaging have made it possible to generate static snapshots of 3D genome structures in single cells (*10*–*15*), live-cell imaging is required to understand the dynamics of chromatin looping (*16*). Furthermore, previous studies have yielded conflicting results as to whether loops are well-defined in single cells (*10*–*15*), perhaps due to the difficulty associated with distinguishing *bona fide* CTCF- and cohesin-mediated loops from mere proximity that emerges stochastically (*16*). Recent pioneering work has visualized enhancer-promoter interactions (*17*, *18*) and long-range V(D)J-chromatin interactions (*19*) in live cells. However, the dynamics of loop extrusion and the lifetime of CTCF/cohesin loops have not yet been quantified in living cells, which we therefore set out to do.

## RESULTS

To visualize the dynamics of CTCF/cohesin looping, we chose as our model system the loop holding together the two CTCF-bound boundaries of the 505 kb *Fbn2* TAD in mouse Embryonic Stem Cells (mESCs). This TAD is verified to be CTCF dependent (*20*) and relatively simple as it contains a single gene, *Fbn2*, which is not expressed in mESCs (**Fig. 1A**). We used genome-editing to homozygously label the left and right CTCF sites of the *Fbn2* TAD with TetO and Anchor3 arrays, which we then visualized by co-expressing the fluorescently tagged binding proteins TetR-3x-mScarlet and EGFP-OR3 (*21*) (clone C36) (**Fig. 1B-D**). We developed a comprehensive image analysis framework, *ConnectTheDots*, to extract trajectories of 3D loop anchor positions from the acquired movies (**Fig. S1**). By optimizing 3D super-resolution live-cell imaging conditions (*16*), we could track *Fbn2* looping dynamics at 20 second resolution for over 2 hours (**Fig. 1D**). After DNA replication in S/G2 phase, it is no longer possible to reliably distinguish intrachromosomal from sister-chromosomal interactions (*16*). We therefore developed and validated a convolutional neural network to filter out replicated and low-quality dots (**Fig. S2**). Thus, we only consider G1 and early S-phase cells.

**Fig. 1.**
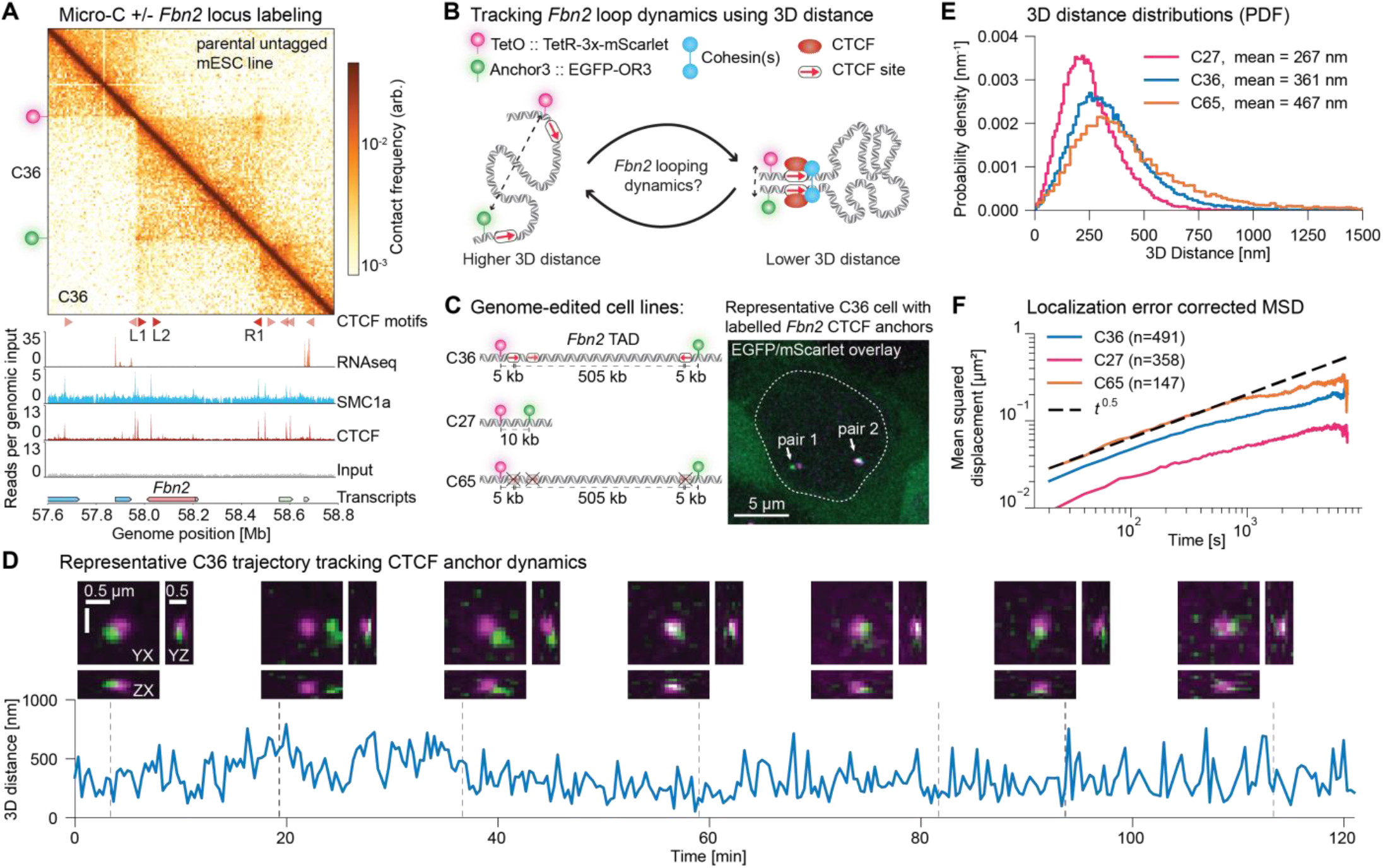
Endogenous labeling and tracking of the *Fbn2* loop with super-resolution live cell imaging. (**A**) Fluorescent labeling of *Fbn2* loop anchors does not perturb the *Fbn2* TAD. mESC Micro-C contact map comparing the parental untagged (C59, top left) and tagged (C36, bottom right) cell lines. Red triangles: CTCF binding with orientation. C36 ChIP-seq shows CTCF (GSM3508478) and cohesin (Smc1a; GSM3508477) binding as compared to Input (GSM3508475). RNA-seq (GSE123636) and transcript annotation tracks (GRCm38) show that *Fbn2* is transcriptionally inactive. Genome coordinates: mm10. (**B**) Overview of tagging and readout using 3D distance. (**C**) Overview of the genome-edited cell lines (left) and a representative maximum intensity projection (MIP) of a cell showing two pairs of “dots” (right). (**D**) Representative 3D trajectory over time of a dot pair. MIPs of the 3D voxels centered on the mScarlet dot (top) and 3D distances between dots (bottom) are shown. (**E**) 3D distance probability density functions of dot pairs (n=32,171; n=46,163; n=13,566 distance measurements for C27, C36, C65 respectively) (**F**) Localization error corrected Mean Squared Displacement (MSD) plots (n=358; n=491; n=147 trajectories in C27; C36; C65 respectively).

To validate our system for tracking *Fbn2* loop dynamics, we carried out a series of control experiments. First, we confirmed using Micro-C (*22*, *23*) that our locus labeling approach did not measurably perturb the *Fbn2* loop (**Fig. 1A**). Second, as a positive looping control we deleted the 505 kb between the CTCF sites, generating clone C27 (**Fig. 1C**). As expected, this significantly reduced the 3D distance (**Fig. 1E**). Third, as a negative control for CTCF-mediated looping, we generated clone C65 (**Fig. 1C**) by homozygously deleting the 3 CTCF motifs in the *Fbn2* TAD (L1, L2, R1; **Fig. 1A**) and validated that this resulted in loss of CTCF binding and cohesin co-localization by ChIP-Seq (**Fig. S3**). As expected, the 3D distance was significantly increased in C65 (**Fig. 1E**). Interestingly, from Mean-Squared Displacement (MSD) analysis, chromatin showed Rouse polymer dynamics with a scaling of MSD~*t*^0.5^ for all three clones (*24*) (**Fig. 1F**). We conclude that our approach faithfully reports on CTCF looping dynamics in live cells without noticeable artifacts.

To elucidate the specific contribution of CTCF and cohesin, we generated cell lines to acutely deplete the cohesin subunit RAD21, CTCF, and the cohesin unloader WAPL. We used genome-editing to endogenously tag these factors with mAID in the C36 line, allowing for degradation with Indole-3-acetic acid (IAA) (*25*) (**Fig. S4**) and validated these cell lines. First, we achieved near-complete depletion of RAD21 and CTCF within 2 hours, while WAPL depletion took 4 hours and was less complete (**Fig. S4**). Second, long-term depletion of RAD21 and CTCF led to cell death as expected for loss of essential proteins (**Fig. S5**), whereas WAPL depletion caused only a minor growth defect and occasionally yielded visible compacted (‘vermicelli’) chromosomes after sustained depletion (*26*). Third, we quantified the protein abundances in the AID cell lines without IAA and note that they are lower likely due to leaky protein depletion (**Fig. S6**). Fourth, we used Micro-C to verify that RAD21 and CTCF depletion led to loss of the *Fbn2* loop or corner peak as expected (*27*–*30*) (**Fig. 2A**) and used ChIP-Seq to verify disrupted CTCF and cohesin chromatin binding (**Fig. S7**). In contrast, WAPL depletion increased corner peak strength (*29*–*31*) (**Fig. 2A**). Thus, our validated AID lines enable efficient and acute protein depletion.

**Fig. 2.**
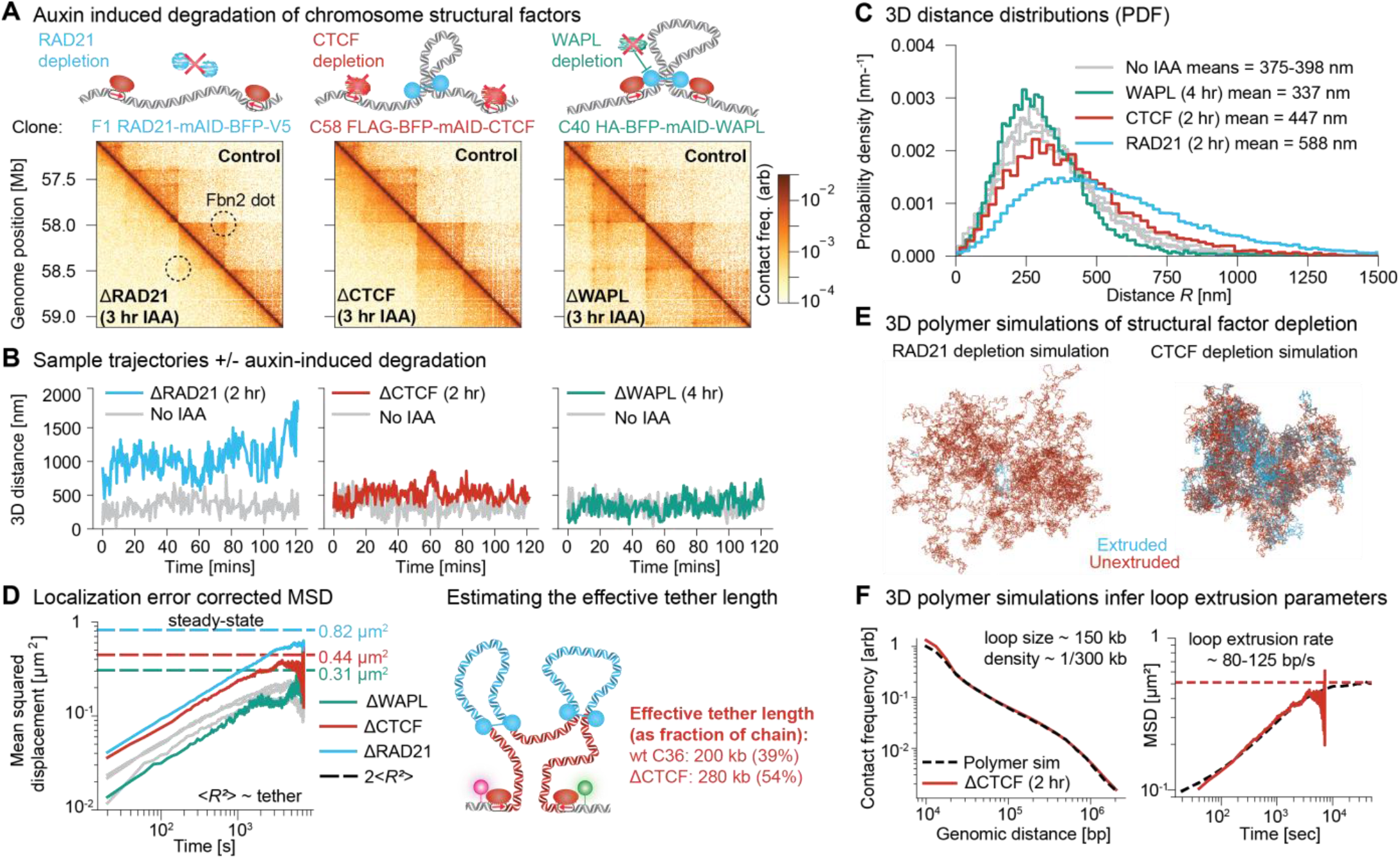
Degradation of CTCF, cohesin, and WAPL reveal their role in loop extrusion and looping-mediated chromosome compaction. (**A**) Micro-C data for the AID-tagged clones for RAD21 (left), CTCF (middle), and WAPL (right), showing control data (no IAA treatment; top half) and protein degradation data (3 hours post IAA; bottom half). Schematics illustrating the expected effect on DNA are shown above each subpanel. (**B**) Representative trajectories with (colored lines) or without IAA treatment (gray lines) for each AID-tagged clone. (**C**) 3D distance probability density functions of dot pairs (n=45,379; n=10,469; n=18,153 distance measurements for ΔRAD21 (2 hr), ΔCTCF (2 hr), ΔWAPL (4 hr) depletion conditions respectively, and n=17,605; n=11,631; n=21,001 for the same clones without treatment). (**D**) Localization error corrected MSD plots for the AID-tagged clones (left) (n=537; n=137; n=215 trajectories in ΔRAD21 (2 hr), ΔCTCF (2 hr), ΔWAPL (4 hr) depletion conditions respectively, and n=183; n=151; n=257 without treatment (gray lines)). The effective tether length is obtained by computing the ratio of the steady-state variance of each clone to the value in the RAD21-depletion condition (note that 2<*R*²> is also the asymptotic value of the MSD; see also Supplementary Material). (**E**) Representative 3D polymer conformation from simulations mimicking the RAD21 (95% cohesin depletion) (left) and CTCF (100% CTCF depletion) (right) depletion conditions. Simulated chromatin segments in loops are colored red and unextruded segments are blue. (**F**) Mean cohesin separations and processivities in simulations are obtained by matching Micro-C contact frequency decay versus genomic distance. Conversion factors for simulated monomer sizes and time steps to nanometers and seconds are obtained by calibrating the ΔCTCF simulations to the 2-hour CTCF depletion MSD plots. The extrusion rate is for two-sided extrusion.

We next studied the specific roles of RAD21, CTCF, and WAPL in loop extrusion *in vivo* by imaging our validated AID lines (**Fig. 2B-C**). Consistent with RAD21 being required for loop extrusion, RAD21 depletion strongly increased the 3D distances (**Fig. 2B-C**). Consistent with CTCF being the boundary factor required for *Fbn2* loop formation (**Fig. 1B**), but not for loop extrusion, CTCF depletion increased 3D distances albeit significantly less than RAD21 depletion (*7*) (**Fig. 2B-C**). Finally, consistent with prior observations that WAPL depletion increases cohesin’s residence time and abundance on chromatin (*26*), potentially allowing it to extrude longer and more stable loops (*29*, *31*), WAPL depletion decreased the 3D distances (**Fig. 2B-C**; ΔWAPL is attenuated due to lower CTCF and cohesin abundances in this clone and less efficient depletion; **Fig. S4, S6**).

To quantify the effect of loop extrusion in our AID lines we turned to polymer physics theory. The Rouse model predicts a linear relationship between chain length and mean squared distance (<*R*²>) between the fluorescent labels (dashed lines in **Fig. 2D**, **Fig. S8**). This relationship allows us to assign an “effective tether length” to each experimental condition by taking ΔRAD21 as a reference value, and assuming that ΔRAD21 represents the fully unextruded state with a genomic separation of 515 kb. We find an effective tether of ~200 kb in wildtype (C36) and ~280 kb in ΔCTCF, corresponding to ~39% and ~54% of the full genomic separation respectively. Thus, conversely, the genomic separation between the two labels shortens by ~46% due to extrusion alone (ΔRAD21 vs. ΔCTCF) and ~61% due to extrusion with boundaries (ΔRAD21 vs. C36). We refer to the latter percentages as the fraction (of the *Fbn2* region) extruded but caution that extrusion of the loop anchors themselves complicates this picture, specifically contributing to the longer effective tether in ΔCTCF. The increased tether length after CTCF depletion is thus consistent with CTCF serving as an extrusion boundary. In summary, we find that on average just over half of the *Fbn2* TAD is extruded into loops.

Having demonstrated that CTCF and cohesin regulate *Fbn2* compaction (**Fig. 2A-D**), we next sought to develop an inference method to quantify loop dynamics. To generate ground truth data to benchmark such a method, we used the Micro-C contact maps (**Fig. 2A**), the absolute 3D distances (**Fig. 2C**), and the MSDs (**Fig. 2D**) in the C36, ΔCTCF, and ΔRAD21 conditions to constrain and parameterize 3D polymer simulations incorporating loop extrusion. Consistent with our ΔRAD21 data, our polymer simulations resulted in chromosome decompaction after near-complete RAD21 depletion (**Fig. 2E**) and accurately matched our experimental data (**Fig. 2F**). We then used these simulations to benchmark a method to infer looped states.

To infer looped state dynamics, we developed Bayesian Inference of Looping Dynamics (BILD). We coarse-grain the possible conformations of the two CTCF sites into two states: 1) a state of sustained contact (the ‘looped state’), where presumably a CTCF-cohesin complex holds together the *Fbn2* loop and 2) all other possible conformations including partially extruded conformations, random contacts, and the fully unlooped conformation (the ‘unlooped state’; **Fig. 3A-B, Fig. S9**). We model the unlooped state (labelled θ=0) as a free Rouse polymer and calibrate it to our ΔCTCF data. For the looped state we introduce a switchable bond between the two CTCF sites (θ=1; **Fig. 3B**), whose strength we set to reproduce the 10 kb distance between the fluorophores, using ΔRAD21 as reference for a free 515 kb chain. Thus, our biological controls allow us to define the looped state as the state that disappears in the ΔCTCF condition. We infer the number of switches *k* between the two states by maximizing the model evidence *E*. We then infer when the switches occur by maximum a posterior estimation (**Fig. 3C**). To reduce false positives, we introduced an evidence margin *ΔE,* which reduces the sensitivity of loop detection but renders the detected looping segments more reliable (**Fig. 3A, Fig. S10**). Our final inference scheme, BILD, accurately inferred both the looped fraction and loop lifetime when applied to our 3D polymer simulation data with experimentally realistic localization uncertainty (**Fig. S10-11**).

**Fig. 3.**
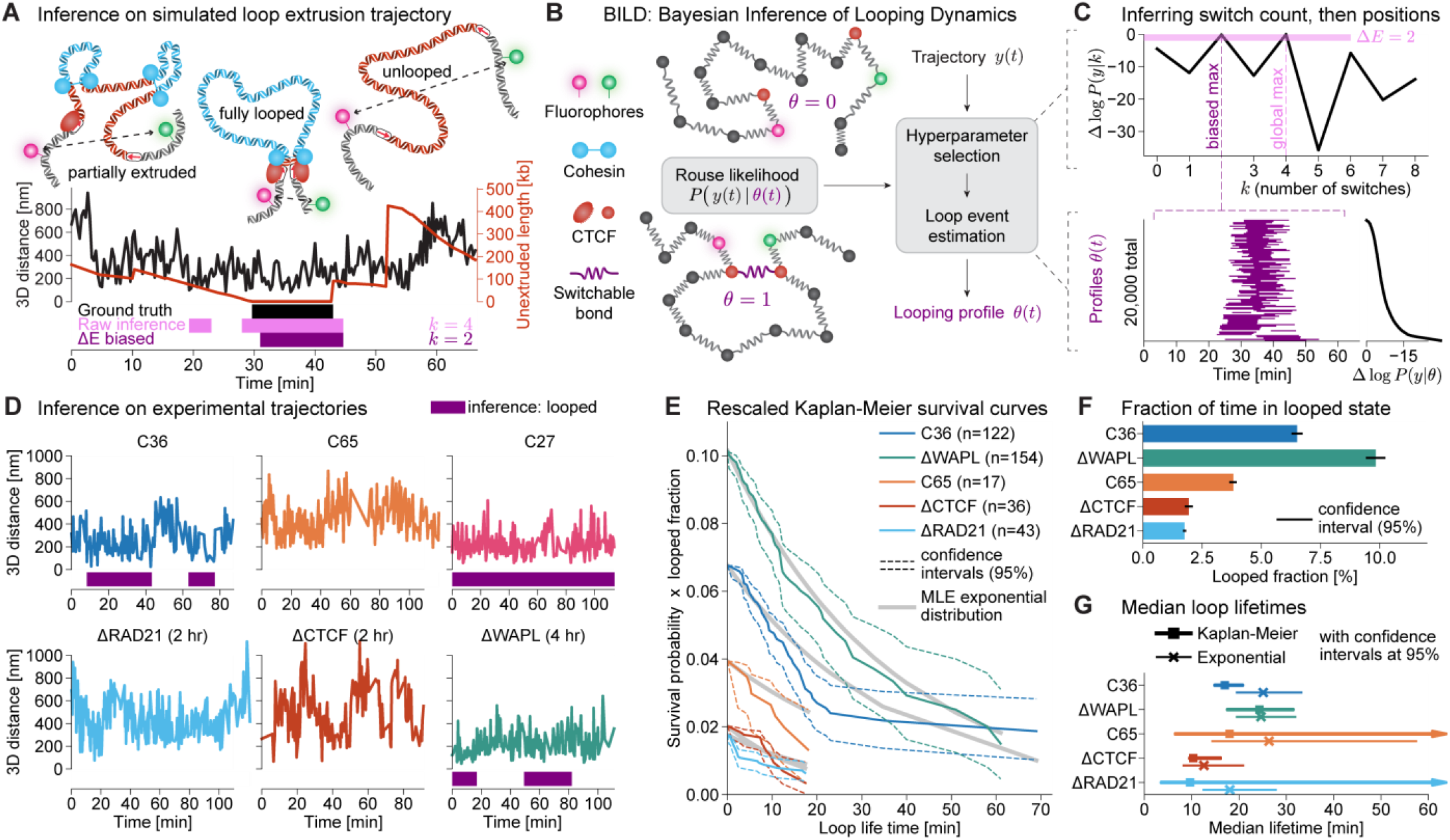
Bayesian Inference of Looping Dynamics (BILD) reveals rare and dynamic CTCF loops. (**A**) Example trajectory from polymer simulations with loop extrusion. Extrusion shortens the effective tether (unextruded length; red) between the CTCF sites; The tether is minimal when cohesin is stalled at CTCF on both sides, which we take as ground truth looping events (black bar). The inference captures these accurately, but raises false positive detections occasionally (pink bars). We limit false positive detections by introducing an evidence bias ΔE (purple bar). (**B**) Schematic overview over BILD. Building on the analytical solution to the Rouse model, we employ an evidence-based optimization scheme to determine the optimal looping profile in two steps. (**C**) BILD procedure. In a first step, we maximize the evidence over the number of switches in the profile. In the second step we then find the best positions for these switches by maximizing the posterior. (**D**) Illustrative examples of inferred profiles on real trajectory data. (**E**) Kaplan-Meier survival curves rescaled by the inferred looped fraction. Gray lines are maximum likelihood fits of a single exponential to the data, accounting for censoring (Supplementary Material). (**F**) Fraction of time the *Fbn2*-locus spends in the fully looped conformation for each of our experimental conditions. Error bars are bootstrapped 95% confidence intervals. (**G**) Median loop lifetimes from the Kaplan-Meier survival curves (squares) or exponential fits (crosses). Confidence intervals are determined from the confidence intervals on the Kaplan-Meier curve and the likelihood function of the exponential fit, respectively. Where the upper confidence limit on the survival curve did not cross below 50% an arrowhead indicates a semi-infinite confidence interval.

We next used BILD to infer looping in our experimental trajectory data (**Fig. 3D-G**). BILD revealed that the *Fbn2* TAD is fully looped ~6.5% (~3%) of the time, but spends ~93.5% (97%) of the time in a fully unlooped or partially extruded conformation (**Fig. 3F**). We use brackets to indicate the looped fraction after false positive correction (**Fig. S11**). In contrast, we observed a minimal looped fraction of ~2% (~0%) in ΔRAD21 and ΔCTCF, and ~4% (~1%) in C65, whereas the looped fraction was significantly increased to ~10% (~6%) in ΔWAPL, consistent with WAPL unloading cohesin from chromatin (*26*). Finally, we estimated the lifetime of the looped state (**Fig. 3E, G**). Lifetime estimation is challenging due to ‘censoring’, which occurs if the trajectory either begins or ends in the looped state. To correct for censoring, we used the Kaplan-Meier estimator of the survival probability (**Fig. 3G**). We also provide an orthogonal estimate of the median lifetime from an exponential fit, which generally agrees with the Kaplan-Meier estimate. Together, these give an estimate of the median loop lifetime of ~10-30 min in the wildtype C36 line (**Fig. 3G, Fig. S11D**). Though our estimates are associated with uncertainty and though we cannot exclude the existence of a very rare but long-lived loop state, these results nevertheless reveal the fully looped CTCF-cohesin complex state to be both rare (~3%) and quite dynamic (median ~10-30 min; mean ~15-45 min). Thus, during an average ~12 hour mESC cell cycle, the looped state will occur ~1-2 times lasting cumulatively ~20-45 min, but the remaining ~11.5 hours will be in the partially extruded or unlooped state.

Wondering how to reconcile a low looped fraction of ~3% with a clear and strong corner peak in the Micro-C map, we set up polymer simulations with loop extrusion. We found that CTCF-mediated stabilization of cohesin was necessary to reproduce both these features in our simulations (**Fig. 4**, **S12**; Supplementary Material), consistent with recent reports (*32*, *33*) (**Fig. 4**, **S12**; Supplementary Material). We confirmed this effect using iFRAP of cohesin, finding that CTCF depletion decreases cohesin’s residence time (**Fig. S13**). Incorporating this effect, we then simulated loop extrusion with a cohesin density of 1/240 kb and processivity of 150 kb (processivity = lifetime * extrusion speed). When cohesin reaches a CTCF site, it has a probability of 12.5% to stall, which, using the estimate of 50% CTCF occupancy (*34*), translates to a ~25% capture efficiency of CTCF. Once stalled by CTCF, cohesin is stabilized 4-fold beyond its base lifetime of ~20 min (*35*) (**Fig. S13**), facilitating the formation of longer loops. These simulations reproduced both our experimental Micro-C maps (**Fig. 4A**) and the median loop lifetime and low looped fraction (**Fig. 4B**).

**Fig. 4.**
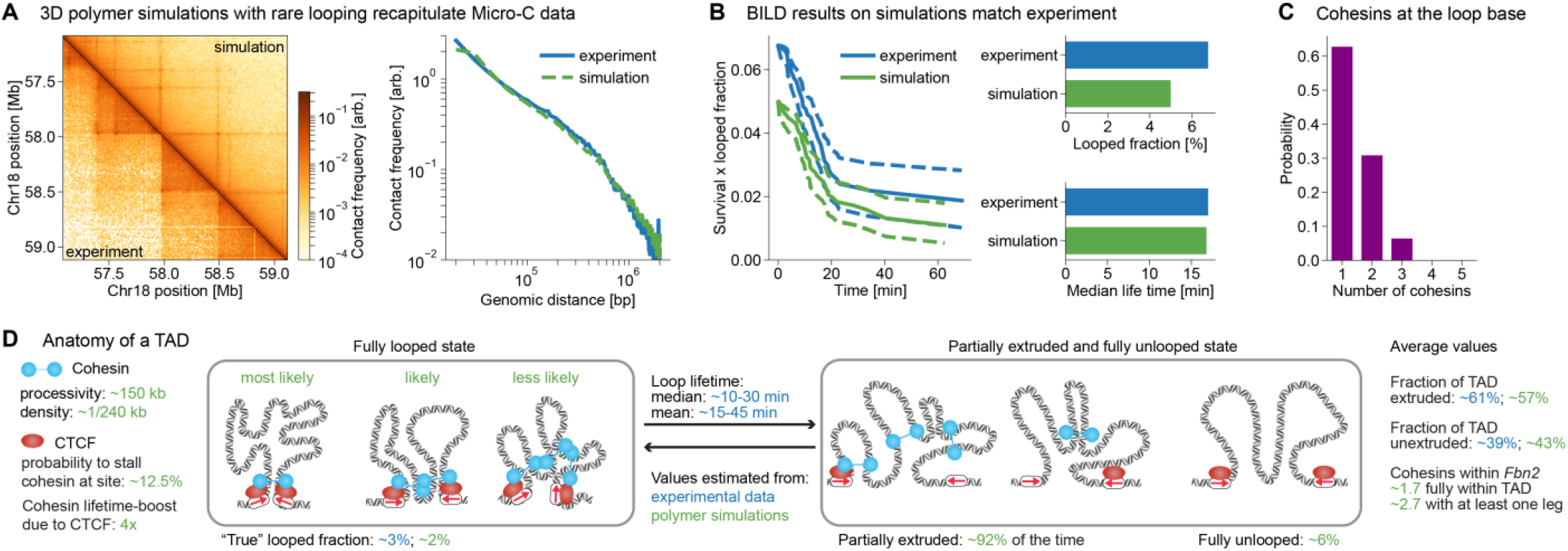
Comprehensive picture of the *Fbn2* TAD. (**A**) Comparison of Micro-C data for the C36 line to *in silico* Micro-C of our best-fit simulation, map (left) and contact probability scaling (right). (**B**) BILD applied to the same simulation (green), comparing to C36 experimental data (blue). (**C**) Number of cohesins forming the looped state in the simulation (n = 18,789). (**D**) “Anatomy” of the *Fbn2* TAD. Quantitative description of the *Fbn2* TAD using both real data (blue) and our best-fit simulation (green). Cohesin processivity and density and CTCF stalling probability and lifetime boost are simulation parameters. Fraction of time in different conformations was extracted from simulation ground truth, using effective tether lengths of 1.1 kb and 505 kb as cutoffs to define “fully looped” and “fully unlooped” respectively. Fraction of TAD unextruded was obtained as mean tether length over the full simulation. Experimental values are from **Figs. 2, 3**.

Together, these results allow us to paint a comprehensive mechanistic picture of the *Fbn2* TAD (**Fig. 4C-D**): most of the time (~92%), the TAD is partially extruded, with ~57-61% of the *Fbn2* region captured in ~1-3 extruding cohesin loops, while ~39-43% remain unextruded. The fully unlooped conformation, as it would be found in the absence of cohesin, occurs only ~6% of the time, while the fully looped state is even more rare at ~3% (~2% in simulations) and has a median lifetime of ~10-30 min. Interestingly, our simulations reveal that the looped state is sometimes held together by multiple cohesins (**Fig. 4C**), which also explains why the loop lifetime can be substantially shorter than the CTCF-stabilized cohesin lifetime (**Fig. S13**). Nevertheless, we stress that both the mechanistic assumptions of our polymer simulations and the experimental data constraining them are associated with uncertainty, resulting in uncertainty of the inferred parameters (**Fig. 4D**). We also note that TADs smaller than the 505 kb *Fbn2* TAD as well as TADs with stronger CTCF boundaries may have a higher looped fraction (Giorgetti and co-workers, personal communication). Furthermore, we propose that our absolute quantification of the *Fbn2* looped fraction may now allow calibrated inference of absolute looped fractions genome-wide, based on Micro-C (*15*).

## DISCUSSION

Our findings reveal the CTCF/cohesin-mediated looped state that holds together CTCF boundaries of TADs to be rare, dynamic, and transient. A key limitation of our study is that it represents just one loop in one cell type. Nevertheless, the *Fbn2* loop is among the strongest quartile of “corner peaks” in Micro-C maps, suggesting that most other similarly sized loops in mESCs are likely weaker than *Fbn2* (**Fig. S14**). Our results thus rule out static models of TADs, where TADs exist in either a fully unlooped state or a fully looped state stably bridged by one cohesin (**Fig. 1B)**. Instead, we show that the *Fbn2* TAD most often exists in a partially extruded state formed by a few cohesins in live cells (~92%; **Fig. 4D**), and that when the rare looped state is formed, it is both transient (~10-30 min median lifetime; **Fig. 4B**) and sometimes bridged by multiple cohesins (**Fig. 4C**). Overall, frequent cohesin-mediated contacts within a TAD rather than rare CTCF-CTCF loops may therefore be more important for regulatory interactions, such as those between enhancers and promoters. Thus, instead of stable loops, we observe a much more dynamic and transitory picture of TADs in live cells (**Fig. 4D**), which may also help explain cell-to-cell variation in 3D genome structure, and consequently stochasticity in downstream processes such as gene expression and cell differentiation.

## Acknowledgements

We thank Drs. Robert Tjian and Xavier Darzacq for hosting early parts of this work, Dr. Jeffrey Alexander for providing the RNA destabilization elements, Lydia Joh for assistance with cloning, and the Hansen, Zechner, and Mirny labs for insightful discussions.

## Funding

This work was supported by NIH grants R00GM130896 (ASH), DP2GM140938 (ASH), R33CA257878 (ASH), UM1HG011536 (ASH and LM), R01GM114190 (LM), NSF grant 2036037 (ASH), the Mathers’ Foundation (ASH), a Pew-Stewart Cancer Research Scholar grant (ASH), Chaires d’excellence Internationale Blaise Pascal (LM), American-Italian Cancer Foundation research fellowship (MG), and core funding from the Max Planck Institute of Molecular Cell Biology and Genetics (CZ).

## Author contributions

ASH conceived and initiated the project. HBB, MG, SGH, LM, CZ, ASH designed the project. ASH performed genome-editing and generated the cell lines. GMD cloned plasmids. MG, AJ, CC, and ASH characterized and validated the cell lines. THSH performed Micro-C. CC performed ChIP-Seq. MG, AJ, and HBB optimized imaging experiments with input from ASH. MG, AJ, and HBB collected the imaging data with acquisition led by MG and AJ. MG and AJ performed control experiments and characterized the AID cell lines. HBB developed image processing pipeline, the CNN, and analyzed the imaging data with input from ASH, SGH, MG, and AJ. HBB performed polymer simulations with input from SGH and LM. MG, AJ, HBB, and ASH annotated trajectory data. SGH and CZ designed BILD with input from HBB, LM, and ASH. SGH developed and benchmarked BILD, applied BILD to trajectory data, and developed MSD analysis with input from HBB and CZ. HBB and SGH analyzed polymer simulations. ASH, LM, and CZ supervised the project. HBB, MG, SGH, AJ, and ASH drafted the manuscript and figures. All authors edited the manuscript and figures.

## Competing interests

Authors declare that they have no competing interests.

## Data and materials availability

All the data and code are available as described in the supplementary materials. The GEO accession number is GSE187487.

## Notes

### Competing Interest Statement

The authors have declared no competing interest.

