## Supplementary Materials for "Dynamics of CTCF and cohesin mediated chromatin looping revealed by live-cell imaging"

**8 Variation in inference results with evidence bias 27**

**9 Software, code, and data availability 27**

**10 Supplementary Figures 28**

**11 Supplementary Tables 41**

**12 Supplementary References 47**

### 1 Cell line generation, culture, and treatment conditions

#### 1.1 Cell culture

Mouse embryonic stem cells (JM8.N4 mESCs) [1] were cultured on plates pre-coated with 0.1% sterile gelatin solution (Sigma-Aldrich, G1890-100G) under feeder free conditions in a base medium consisting of KnockOut DMEM (Thermofisher, 10829-018) with 15% FBS (HyClone, SH30396.03, Lot. No. AE28209315) and 1000U/mL LIF (home-made [2]), 1 mM MEM Non-Essential Amino Acid Solution (Thermofisher, 11140-050), 2 mM GlutaMAX (Thermofisher, 35050-061), 100 µg/ml Penicillin-Streptomycin (Thermofisher, 15140-122), and 0.1 mM 2-mercapoethanol (Sigma, M-3148) supplemented with 2i, 10 µM MEK inhibitor (Tocris, PD0325901) and 3 µM GSK inhibitor (Sigma, SML1046-25MG). mESCs were fed daily by replacing half the medium and passaged every two days with 0.05% Trypsin-EDTA (Thermo Fisher Scientific, 25300062).

#### 1.2 Genome-editing and cell line generation

Genome-editing was performed in JM8.N4 mESCs using CRISPR/Cas9 largely according to published procedures [3], but with modifications. We co-transfected a Cas9 plasmid (encoding Cas9, Venus, and the sgRNA) and a repair plasmid (DNA segment to be inserted typically flanked by ∼500 bp of homology on the left and on the right) using Lipofectamine 3000 (ThermoFisher L3000015) according to manufacturer's protocol, using a ratio of 2 µg repair vector and 1 µg Cas9 vector per well in a 6-well plate (1:2 w/w). We designed sgRNAs using the CRISPOR tool [4]. For each insertion we designed 2-4 individual sgRNAs, cloned them into the Cas9 plasmid and then co-transfected the repair plasmid and a single sgRNA plasmid individually into a well of a 6-well plate. The following day (16-28 hours later) we pooled the several 6-wells containing transfected cells, collected cells by trypsinization and then sorted successfully transfected cells using FACS (gating for single cells based on SSC/FSC; gating for YFP fluorescence by using an untransfected control to set the negative gate). mESCs were then plated at a density of ∼5,000-10,000 cells per P15 plate. The medium in the P15 was supplemented with gentamicin (ThermoFisher #15750072, at 10 mg/mL). Cells were fed every 2 days and single colonies picked inside the biosafety cabinet under a Lynx EVO stereomicroscope (Vision Engineering) using a 20 $\mu$L pipette tip, passaged into 96-well plates, expanded, crude DNA collected using DirectPCR Lysis Reagent (Viagen Biotech #302-C) and genotyped using a 3-primer PCR strategy (two primers external to the homology arms and one primer inside the DNA segment to be inserted). Desired clones were further expanded, high-quality genomic DNA collected (Quick-DNA Kit, Zymo Research), clones were further verified using PCR with multiple primer combinations on high-quality genomic DNA, frozen down and used for downstream experiments after additional clone-specific validation. Pathogen testing of the parent cell line (C59 from [2]) was performed using an IMPACT II assay (performed by IDEXX BioResearch), which tests for pathogens including Ectromelia, EDIM, LCMV, LDEV, MAV1, MAV2, mCMV, MHV, MNV, MPV, MVM, Mycoplasma pulmonis, Mycoplasma sp., Polyoma, PVM, REO3, Sendai, and TMEV. The mESC line was negative for all tested pathogens.

To visualize CTCF looping in living cells we chose the CTCF loop on chromosome 18 surrounding the *Fbn2* gene in mESC (hereafter referred to as the "*Fbn2* loop", since it satisfies the following criteria: 1) simple, contains a single gene (*Fbn2*) not expressed in mESCs; 2) strong and clear loop in Hi-C [5]; 3) large (505 kb); 4) previously validated by de Wit *et al.* to be CTCF-dependent [6]. The *Fbn2* loop is anchored by two CTCF sites on the left (L1, L2, with L2 seemingly only playing a minor role [6]) and one CTCF site on the right (R1). To place the inserted arrays sufficiently far away from the L1 and R1 CTCF sites, but nevertheless close enough to provide an accurate reporter of the CTCF site location, we chose to place the arrays ∼5 kb from the endogenous CTCF sites. Specifically, the L1 (5'-agtttctgctcggcgactgcgactGCGAACAGTAGATGGCAGTAttgctcctgtctttgggaaagagcctctggagc-3') CTCF site is at ∼chr18:58,136,460 (mm9) and we placed the array at ∼chr18:58,131,000 (mm9). The R1 (5'-gggggggggaactgaaaatttgtgaaaTACTGCCCTCTAGTGGAAAAgtaaggttttgctgtatctgtagaaaga-3') CTCF site is at ∼ 58,641,875 (mm9) and we placed the array at ∼chr18:58,646,000 (mm9).

First, we inserted an array containing ∼224x TetO operator sites at ∼chr18:58,131,000 (mm9) using repair plasmid pASH83 and sgRNA plasmid pASH84, 85, 86 (see **Table S1** for plasmids and **Table S2** for sgRNA sequences and full details). The repair plasmid contains ∼224x TetO operators with ∼20-50 bp spacing as well as PGK promoter driven hygromycin resistance. The inserted TetO array is ∼10 kb. After co-transfection of the repair plasmid and one of the three sgRNAs, cells were pooled and selected using 50 mg/mL hygromycin until no more cells survived in an untransfected control plate. Cells were then plated at very low density in a P15 plate, colonies picked and genotyped. Repetitive arrays can be genetically unstable and recombine out. To test for array integrity in homozygous TetO-clones we performed qPCR comparing genomic DNA isolated from a handful of homozygous clones to genomic DNA isolated from wild-type mESCs mixed with the TetO plasmid (containing ∼224x TetO repeats) in a 1:1 molar ratio (5 pg plasmid per 1 $\mu$g genomic DNA). Two pairs of control primers and two pairs of TetO targeting primers were used (see **Table S2** for primer sequences). qPCR was performed using SYBR Green Master Mix and the following clones were found to have maintained array integrity: 1-B12, 2-E1, 2-D8, and 2-F12.

Next, clones 2-E1, 2-D8, and 2-F12 (chr18:58.131::TetO) were used for inserting a ∼1 kb Anchor3 array [7].

We obtained the Anchor3 array from NeoVirTech and cloned it into a pENTR plasmid to generate repair plasmid pASH89, which contains the 982 bp Anchor3 sequence, encodes puromycin resistance and homology flanking chr18:58,646,000. The sgRNA plasmids were pASH101, 102, 103 (see **Table S2** for sgRNA sequences and full details). Genome-editing was performed as above: cells were individually co-transfected with the repair plasmid and one sgRNA plasmid, pooled, selected using puromycin (1 mg/mL), plated at low density, single colonies picked, clones expanded and genotyped (PCR primers can be found in **Table S2**). Clones 26 and 47 were validated and used for downstream work and Sanger sequenced.

Having inserted the arrays, the fluorescently-tagged DNA-Binding proteins were next inserted using PiggyBac transposition. Briefly, TetR was cloned into plasmid pASH135 expressing TetR-NLS-3x mScarlet-NLS followed by 4 RNA destabilization elements (DE 5'-ttatttatt-3'; the DEs were identical to those reported by Alexander *et al.* [8]). TetR-NLS-3x-mScarlet-NLS is expressed from a L30 promoter and the Nuclear Localization Signals (NLSs) are the c-Myc and nucleoplasmin NLSs, respectively. TetR was C-terminally tagged with 3 copies of mScarlet, connected by SGGG-linkers [9]. The Anchor3 array binding protein is OR3 [7]. OR3 was cloned into pASH136 expressing EGFP-OR3. N-terminally EGFP-tagged OR3 is expressed from an L30 promoter and followed by 4x RNA destabilization elements (4xDE). Next, these two plasmids were co-transfected together with a plasmid encoding Super PiggyBac Transposase (500 ng each per 6-well) using Lipofectamine 3000. Two days later, transfected cells were sorted using FACS for EGFP-OR3 (530/40 nm emission, 488 nm excitation) and TetR-3x-mScarlet (585/29 nm emission, 561 nm excitation). Double-positive cells with modest, but not too high, expression of both proteins were sorted. Sorted cells were plated at low density, individual colonies were picked and plated on glass-bottom 96-well plates. 120 clones were screened for optimal expression level on an Opera Phenix High-Content Screening System (PerkinElmer), and clone C36 was selected for having optimal expression of both TetR-3x-mScarlet and EGFP-OR3. Clone C36 (JM8.N4 mESC; C59; chr18:58.131 224x TetO; chr18:58.646 Anchor3; pL30 TetR-NLS-3x-mScarlet-NLS-DEx4; pL30 EGFP-OR3-DEx4) was used for downstream experiments, including both imaging and further genome editing.

To enable acute depletion of CTCF, RAD21, and WAPL using the Auxin-Inducible-Degron (AID) system [10], it is necessary to express the ubiquitin ligase osTir1. We introduced osTir1 expression using a PiggyBac plasmid encoding osTIR1-P2A-BSD which also affords blasticidin resistance [11, 12]. This plasmid, pASH133, expresses osTir1 from an EF1a promoter. OsTir1 contains 3x Myc-epitopes (EQKLISEEDL) at the C-terminus, P2A, and then the blasticidin resistance gene. The C36 clone was co-transfected with pASH133 and Super PiggyBac transposase, selected with blasticidin (5 µg/mL, ThermoFisher #A1113903) to generate C36 osTir1 (JM8.N4 mESC; C59; chr18:58.131 224x TetO; chr18:58.646 Anchor3; pL30 TetR-NLS-3x-mScarlet-NLS-DEx4; pL30 EGFP-OR3-DEx4; pEF1a osTir1-3x-Myc-P2A-BSD).

Next, we endogenously tagged CTCF, RAD21, and WAPL with mAID [13] in the C36 osTir mESC line. Specifically, we cloned plasmids with flanking homology to the genome and encoding FLAG-TagBFP-mAID-CTCF (pASH107), RAD21-tagBFP-mAID-V5 (pASH112), and HA-tagBFP-mAID-WAPL (pASH117). 3 sgRNAs (WAPL) or 4 sgRNAs (CTCF, RAD21) were designed for each insertion (see **Table S2**). As above, the C36 osTir1 line was co-transfected with one sgRNA plasmid and the repair plasmid, sorted based on Venus fluorescence using FACS, plated at low density, and single clones verified using PCR (and additional validation including Western Blot; PCR primers can be found in **Table S2**). Individual clones were further verified by Sanger sequencing and we performed an initial test of IAA-mediated depletion: clones were grown in a 6-well plate, treated with 500 mM IAA for 3 hours and Western blotting was carried out to compare IAA-treated vs. untreated. Several clones were identified for each AID line, and the following clones identified as optimal: clone C58 for FLAG-BFP-mAID-CTCF (JM8.N4 mESC; C59; chr18:58.131 224x TetO; chr18:58.646 Anchor3; pL30 TetR-NLS-3x-mScarlet-NLS-DEx4; pL30 EGFP-OR3-DEx4; C36 osTir1; FLAG-BFP-mAID-CTCF), clone C40 for HA-BFP-mAID-WAPL (JM8.N4 mESC; C59; chr18:58.131 224x TetO; chr18:58.646 Anchor3; pL30 TetR-NLS-3x-mScarlet-NLS-DEx4; pL30 EGFP-OR3-DEx4; C36 osTir1; HA-BFP-mAID-WAPL), and clone F1 for RAD21-mAID-BFP-V5 (JM8.N4 mESC; C59; chr18:58.131 224x TetO; chr18:58.646 Anchor3; pL30 TetR-NLS-3x-mScarlet-NLS-DEx4; pL30 EGFP-OR3-DEx4; C36 osTir1; RAD21-mAID-BFP-V5). These clones were also verified using Sanger sequencing.

Next, we sought to generate a 'positive' and 'negative' control for CTCF-mediated looping by deleting the $\sim$505 kb of DNA that separate the L1 and R1 CTCF sites and by endogenously and homozygously deleting the L1, L2, and R1 CTCF binding sites, respectively. We performed these deletions in the C36 line (JM8.N4 mESC; C59; chr18:58.131 224x TetO; chr18:58.646 Anchor3; pL30 TetR-NLS-3x-mScarlet-NLS-DEx4; pL30 EGFP-OR3-DEx4).

First, we sought to delete the L1, L2, and R1 binding sites. To avoid issues of partial residual binding that can be associated with small deletions/mutations in the long CTCF binding site, we opted to delete $\sim$20-100 bp on both sides of the CTCF binding site, and we thus designed 2-3 sgRNAs for both the left and the right hand side of each of the three CTCF binding sites (see **Table S1** for plasmids and primers). We used PCR primers outside of the area targeted by the sgRNAs to screen for clones with homozygous deletion of L1, L2, and R1. We deleted the three pairs of CTCF sites in two steps. In the first step we deleted L2 and R1. Since cells are diploid, and since the repair after cutting with Cas9 on both sides of the CTCF site can be variable, we PCR amplified the region with the CTCF site deletion, cloned it into a pBlueScript plasmid using blunt-end ligation and then Sanger sequenced at least four clones to capture both alleles. The deletion of the L2 and R1 sites was identical on both alleles. 20 bp upstream and 24 bp downstream of L2 was deleted on both alleles. Thus, L2 is homozygously deleted. The deletion of R1 was also homozygous and identical on both alleles. We deleted from 54 bp upstream of R1 to 58 bp downstream of R1. The ΔL2 and ΔR1 clone, clone C44, was isolated and expanded (JM8.N4 mESC; C59; chr18:58.131 224x TetO; chr18:58.646 Anchor3; pL30 TetR-NLS-3x-mScarlet-NLS-DEx4; pL30 EGFP-OR3-DEx4; homozygous ΔL2; homozygous ΔR1; clone C44). Subsequently, we re-targeted L1 in C44 and generated a ∼300 bp deletion surrounding the L1 site. This resulted in clone C65 with all three CTCF binding sites homozygously deleted "C65 ΔL1, ΔL2, and ΔR1" (JM8.N4 mESC; C59; chr18:58.131 224x TetO; chr18:58.646 Anchor3; pL30 TetR-NLS-3x-mScarlet-NLS-DEx4; pL30 EGFP-OR3-DEx4; homozygous ΔL1; homozygous ΔL2; homozygous ΔR1; clone C65). To further validate that CTCF and cohesin no longer binds the L1, L2, and R1 binding sites in C65, we performed ChIP-Seq using antibodies against CTCF as well as cohesin subunit Smc1a and found complete loss of binding to the L1, L2, and R1 binding sites as measured by ChIP-Seq (**Fig. S3)**. We refer to C65 as a 'negative' control for CTCF-mediated looping, since all endogenous CTCF binding sites have been deleted.

Finally, to generate a 'positive' control for looping, we deleted the ∼505 kb that separate L1 and R1 CTCF binding sites. Here we used sgRNAs targeting L1 and R1 and primers external to L1 and R1 to screen for clones with a homozygous deletion of all 505 kb. We isolated clone C27, PCR amplified the segment from left of L1 to right of R1, cloned the PCR products into a pBlueScript plasmid using blunt-end ligation and performed Sanger sequencing. In C27, the deletions on the two alleles are slightly different. The first allele has 30 bp upstream of L1 deleted going to 61 bp downstream of R1. Thus, the first allele has both the L1 and R1 sites deleted. The second allele is joined 28 bp downstream of L1 to 62 bp downstream of R1. Thus, the second allele retains the L1 CTCF site but is without the R1 CTCF site. Thus, both alleles has ∼505 kb deleted and thus reflect a putative CTCF loop, with the second allele having 77 bp more DNA deleted than the first allele. Since in clone C27 the two arrays are brought into the same proximity as in a putative loop between the two CTCF sites, we refer to is as a positive control for looping (JM8.N4 mESC; C59; chr18:58.131 224x TetO; chr18:58.646 Anchor3; pL30 TetR-NLS-3x-mScarlet-NLS-DEx4; pL30 EGFP-OR3-DEx4; homozygous Δ505kb (L1-R1); clone C27).

##### 1.3 IAA mediated protein depletion and protein analysis

To deplete proteins fused with mAID, we added IAA to a final concentration of 500 µM in culture medium, from a 500x stock solution of 250 mM in DMSO. Target depletion was measured by western blotting. The procedure was performed with adaptation from Cattoglio *et al.*, [14]. Cells were grown in 6 well plates. After IAA treatment, samples were washed with cold PBS containing protease inhibitors: PMSF (Millipore Sigma, Roche, catalog number: 11359061001), aprotinin (Millipore Sigma, catalog number: A6279), benzamidine (Millipore Sigma, catalog number: B6506). Keeping the plate on ice, we lysed cells with 200 µL of high-salt lysis buffer (NaCl 500 mM, Hepes 25 mM, MgCl2 1 mM, EDTA 0.2 mM, NP-40 0.5%), collected residues and transferred to tubes containing 4X Laemmli Sample Buffer (Bio-Rad, catalog number: 161-0747). Samples were boiled for 15 minutes, stored in -20℃ or kept on ice until loaded.

Western blotting was performed using either 4–15% Criterion™ TGX™ Precast Midi Protein Gel (Bio-Rad, catalog number: 5671084) or 4–15% Mini-PROTEAN,TGX™ (Bio-Rad, catalog number: 4561084). We ran samples using premade running buffer (Bio-Rad, catalog number: 1610732) with the Precision Plus Protein™ Dual Color Standards (Bio-Rad, catalog number: 1610374S). We then performed protein transfer for two hours at 120V to a nitrocellulose membrane. We blocked membranes with 5% Nonfat Dry Milk, Blotting Grade (Genesee Scientific catalogue number: 20-241) in TBS complemented with Tween20 (VWR, catalogue number: M147-1L) at a final concentration of 0.05%. we performed the immunostainings with the following antibodies: RAD21 1:1000 (Abcam, catalogue number: ab154769), HA 1:1000 (Abcam, catalogue number: ab9110), CTCF 1:1000 (Millipore, catalogue number: 07-729), ACTB 1:5000 (Sigma, catalogue number: A2228), trueblot anti rabbit HRP 1:10000 (Rockland, catalogue number: 18-8816-31), trueblot anti mouse HRP 1:10000 (Rockland, catalog number: 18-8816-31). We performed chemiluminescence reaction with the Pierce™ ECL Western Blotting Substrate (Thermo Fischer Scientific, catalog number: 32209) and measured with the ChemiDoc MP Imaging System. Protein quantification was performed with Fiji [15].

#### 2 Microscopy experiments and analysis

##### 2.1 Super-Resolution Live-Cell Imaging (SRLCI) to track dynamics of *Fbn2* loop anchors

For Super-Resolution Live-Cell Imaging, mESCs were grown for two days on 35 mm no 1.5H glass-bottom imaging dishes (MatTek, Ashland, MA, P35G-1.5-14C) coated with MatriGel (Corning, 354277), according to manufacturer's instructions. mESCs were grown overnight on the imaging dish in 2i supplemented base medium. One day before imaging, the medium was replaced with the base medium without 2i. On the day of imaging, the medium was changed to imaging medium: DMEM without phenol red (Thermofisher, 31-053-028) and 15% tetracycline-free FBS (Takara, 631107), 1000U/mL LIF (home-made [2]), 1 mM MEM Non-Essential Amino Acid Solution (Thermofisher, 11140-050), 2 mM GlutaMAX (Thermofisher, 35050-061), 100 µg/ml Penicillin-Streptomycin (Thermofisher, 15140-122), and 0.1 mM 2-mercapoethanol (Sigma, M-3148). 5-Phenyl-1H-indole-3-acetic acid (IAA, Bioacademia 30-003) was dissolved to 250 mM in DMSO. The lines C36, C27, and C65, were imaged without administration of IAA. The lines F1M-RAD21 and C58-CTCF were imaged with no IAA and after 2 and 4 hours with 500 µM IAA. The line C40-WAPL was imaged with no IAA and after 4 and 6 hours with 500 µM IAA.

Movie acquisition was performed in a humidified incubation chamber at 37° C supplied with 5.5% $CO_2$. Samples were loaded on the objective (Zeiss Plan-Apochromat 63x/NA1.40) with immersion oil Immersol 518F (Zeiss). Before the acquisition, samples were allowed to equilibrate to 37° C for about 15 minutes. Acquisition was performed on a LSM900 Airyscan 2 (Zeiss) microscope equipped with an Airyscan detector GaAsP-PMT (Zeiss) with the following parameters: Diode laser 488 nm, 10 mW, laser class 3B: 0.5%, Diode laser (SHG) 561 nm, 10 mW, laser class 3B: 0.05%, pixel dwell time: 1.30 µs resulting a frame time of 541.82 ms, and 780 V detector gain. We employed a multiplex acquisition CO-2Y modality with definite focus strategy. For each movie we recorded 365 frames of 49.69 µm x 49.69 µm (584 x 584 pixels, pixel size: 0.085 µm x µm), each separated by an interval of 20 seconds for a total of just over 2 hours. Each frame was composed of 30 z-stacks separated by 0.25 µm, for a total height of 7.25 µm.

To correct for chromatic aberrations, we recorded the chromatic offset daily with 100 nm TetraSpeck microspheres diluted 1:10,000 in MatriGel (Thermo Fisher Scientific, T7279). TetraSpeck beads were added to mESCs in imaging medium to allow for Airyscan detector alignment and realistic estimate of the chromatic offset. AiryScan calibration was performed by calibrating the detector on bright colonies until stabilization, then automatic calibration was switched off. Once a week we recorded chromatic offset in the Z-axis by adding TetraSpeck microspheres on top of mESC colonies. To record the chromatic offset acquisition parameters were set as follows: 488 nm laser: 1.5%, 561 nm laser: 1.5%, pixel time: 1.30 µs with a frame time of 541.82 ms, and 780 V detector gain. Prior to data analysis, raw movies were subjected to 3D Airyscan processing with low strength autofilter by Zen 3.0 software (Zeiss). To image the characteristic “Vermicelli” pattern of RAD21 after prolonged WAPL depletion, we treated the HA-tagBFP-mAID-WAPL cell line with IAA for 6 days and performed confocal imaging every day after staining RAD21-$\mathrm{SNAP}_\mathrm{f}$-V5 with cp-SNAP-JF549 500 nM.

#### 2.2 ConnectTheDots: Image processing to obtain dot pair trajectories

The 3D image times-series were processed into dot pair trajectories using a custom pipeline written in snakemake and our connect_the_dots python package (https://github.com/ahansenlab/connect_the_dots) (**Fig. S1A**). Briefly, .czi formatted 3D Airyscan processed image time-series were loaded into Python using the python-bioformats (https://pythonhosted.org/python-bioformats/) library which was developed for and is used by the image analysis software CellProfiler (cellprofiler.org). We performed a 4D Haar wavelet decomposition (https://pywavelets.readthedocs.io/), filtering out the highest and lowest two zoom level coefficients to reconstruct the entire time-series with minimal levels of cytoplasmic background signal in the eGFP channel and other inhomogeneities.

Dots were identified using an iterative approach (connect_the__dots.tracking.get_localizations_iterative()) (**Fig. S1B**) which uses a pixel intensity percentile threshold of 99.5% on the filtered data followed by a morphological filter to select for dot-sized particles. Only dots with initial sizes between 20 and 10000 pixels (i.e. in volume) were kept for further analysis. Dot localizations were joined into trajectories (separately for each channel) using trackpy (http://soft-matter.github.io/trackpy/v0.5.0/) with a search range of search range of $15 \times 15 \times 5$ pixels$^3$ (X $\times$ Y $\times$ Z dimensions) and a memory allowance of 5 consecutive missing frames. If the trajectory creation via trackpy failed (e.g. due to too many tracked or overlapping objects), the iterative localization algorithm sequentially altered the morphological filtering size thresholds by small increments (e.g. of 5 pixels) and altered the intensity percentile threshold cutoff for the Z-stacks (by 0.1% increments) until the tracking was successful. This procedure ensured that the majority of dots were tracked. Only trajectories with a minimum of 15 frames were kept which helped eliminate the vast majority of tracks arising from background “noise”. Trajectories were then joined across channels in space and time by a combination of metrics including minimizing the distance between trajectories and a positional displacement correlation coefficients cutoff of 0.5.

To maximize the lengths and quality of trajectories, we used the information contained in both imaging channels to fill in any missing data from the original localization. Dot positions for missing frames were then estimated by interpolation or using the displacements from the dot in the other channel plus the original dots’ current position. Finally, we re-analyzed the original movies (i.e. non-wavelet filtered). Using the filled-in trajectories with estimated and known dot positions, we retrieved a voxel of size $12 \times 12 \times 8$ pixels$^3$ (X $\times$ Y $\times$ Z dimensions) around the best-guess dot position. Within the voxel, the maximum intensity pixel location was used to identify the potential dot localization center. Then, using a fixed ellipsoid mask of radii $r_{xy} = 4$ pixels and $r_z$=3 pixels, we computed the intensity-weighted center of mass localization of the chromosome fiber giving us the final dot (X,Y,Z) coordinates (i.e. before registration error correction) (**Fig. S1C**). This voxel was stored alongside the dot coordinates to be later used for quality control of the localization using a convolutional neural network (described in section 2.4 below) (**Fig. S2**). After obtaining the final dot coordinates, XY projection movies with trajectories and particle IDs overlayed were created for human verification and curation of trajectories.

#### 2.3 Registration error and chromatic shift correction

In order to correctly calculate the 3D distance between the GFP and mScarlet color channels, it is necessary to correct for the chromatic shift between the two color channels. To estimate the chromatic shift, we used the same image processing pipeline for tracking dots to localize and identify bead positions. The registration between the color channels varied within the imaging field of view. Therefore, to obtain a function $f(x, y, z)$ that accounted for the registration error per pixel position $(x, y, z)$, we separately projected the registration errors onto each dimension X, Y, Z and used a Savitsky-Golay filter of order 1, with a window size of 91 points (in X, and Y) or 21 points (in Z) to obtain the interpolating polynomial functions $r(x)$, $r(y)$, $r(z)$. Then, $f(x, y, z) = r(x)r(y)r(z)$. Subsequently, all dot coordinates in the GFP channel were corrected based on $f(x, y, z)$. To account for registration drift over time and maximize the accuracy of the bead-based registration error correction, we grouped beads collected from adjacent days (using a 7-day moving average window) to compute a unique $f(x, y, z)$ for each imaging day.

#### 2.4 Creating a convolutional neural network (CNN) to classify dot localizations

We created a dot classifier that could distinguish between good and poor dot localization to help filter for low-confidence localization or tracking errors within a trajectory. A total of 5234 dot pairs were initially identified by a human researcher and classified as one of the following: (A) Good (i.e. good localization and no replication), or (B) Bad (i.e. either poor localization or replication). Dots were evaluated based on their overall shape (e.g. unimodal vs. multimodal), size and intensity of the dot, the signal-to-noise, and the localization accuracy. The 5234 dot pairs were subsequently blindly re-evaluated by 4 humans (including the original identifier) and re-classified. The results for each dot were stored. In the end, a total of 2249 dots were classified as "Good" and 2985 classified as "Bad" (see examples in **Fig. S2A**. For the classification, we found between 79-87% agreement between any two researchers (at the 95% confidence level), 71-83% between any three researchers, and 68% agreement between all four researchers. To train the CNN, we randomly split the data into 85% training, 15% validation. We used Keras (https://keras.io/), a Python deep learning API, to create our CNN model. Our model layers (see **Fig. S2C**) were given varying combinations of "selu" or "sigmoid" activation functions and "lecun_normal" kernel initializers. To train our final model, we used a total of 60 epochs with batch sizes of 32 data points, a learning rate of $3 \times 10^{-4}$, a Nesterov momentum of 0.9, and a binary cross-entropy loss function. Our "Good/Bad" classifier consistently achieved between 80-85% correct recall, which was close to the level of consistency between humans.

Next, we created a second, independent training data set focused exclusively on unimodal vs multimodal localization to help identify trajectories with replicated sister chromosomes (**Fig. S2B**). A total of 4734 dot pairs were identified as (A) Replicated (2569 of 4734) or (B) Unreplicated (2165 of 4734). We found between 85-91% agreement between any two researchers (at the 95% confidence level), 84-90% between any three researchers (at the 95% confidence level), and 78% agreement between all four researchers. Our model layers (see **Fig. S2C**) were given varying combinations of "selu" or "sigmoid" activation functions and "lecun_normal" kernel initializers. To train our final model, we used 90% training to 10% validation split, a total of 250 epochs with batch sizes of 32 data points, a learning rate of $5 \times 10^{-4}$, a Nesterov momentum of 0.9, and a binary cross-entropy loss function. Our "Replicated/Unreplicated" classifier achieved an accuracy between 62-73% on individual dots; this was sufficient to classify trajectories (as a whole) as being likely replicated (see method below) with good agreement to human evaluation.

#### 2.5 Filtering trajectories for replicated dots and poor localization using the CNN

We used the CNN to filter trajectories in two steps. First, we removed entire trajectories that contained substantial signs of chromosome replication. Second, from the remaining unreplicated trajectories, we identified individual dots within a trajectory that were poorly localized due to high background noise, possible tracking errors, or potential chromosome replication that occurred during the course of the two hour imaging session. We discarded trajectories if they met the following criterion: Each dot was given two classifications, one of (Good, Bad) and one of (Unreplicated, Replicated) (see **Fig. S2A,B**). If the number of (Bad, Replicated) localizations in a trajectories exceeded 2 times the number of (Good, Unreplicated) localizations, the trajectory was discarded. All other trajectories were then evaluated by a human researcher for other signs of tracking errors or replication by watching the maximum intensity projection image time-series of the tracked dots.

For the final reporting of unreplicated trajectories, we produced two versions: one with all of the dot localizations, and a filtered set where dots identified as "Bad" by the "Good/Bad" classifier were removed. The filtered data sets were used as inputs to the majority of analyses, including the 3D distance histograms, MSDs, and the looping inference scheme.

#### 2.6 Filtering step-like tracking errors

Tracking errors occasionally occurred, for example, due to homolog switching, or due to occasional high levels of punctate cytoplasmic eGFP signal in some cells. Such tracking errors often appeared as sustained "steps" in the 3D pair distance, whereby the localization algorithm switched from correctly localized to incorrectly localized particle pairs (or vice versa), most often at the end of a trajectory. We found the following procedure could eliminate the majority of such events: First, we computed the median absolute deviation (MAD) of all displacements along the trajectory, and flagged individual points that exceeded distances three MADs above the median. Second, we flagged all points where the displacement from one localization to the next exceeded 500 nm. If any contiguous segment of distances met both criteria (i.e. distances exceeded 3 MADs and were connected to an initial large displacement of 500 nm), then the segment was eliminated from the trajectory. We found this procedure was robust to the choice of displacement (e.g. results did not significantly change if 400 nm or 600 nm were used instead of 500 nm), and eliminated less than 2% of the data points overall. The final trajectory data is summarized in **Table S4**.

#### 2.7 Measuring particle localization uncertainty

Localization errors were computed from maximum likelihood estimation of the MSD as described in section 5, using a Rouse model with localization error. Errors for each dimension were computed separately as shown in **Table S3**.

#### 2.8 Inverse Fluorecence Recovery After Photobleaching (iFRAP) of cohesin

To characterize the effect of CTCF depletion on cohesin dynamics, we performed Inverse Fluorecence Recovery After Photobleaching (iFRAP) in the FLAG-BFP-mAID-CTCF cell line (clone C58). To visualize cohesin, we labeled endogenously tagged RAD21-SNAP$_{\mathrm{f}}$-V5 with 500 nM cp-SNAP-JF549 for 20 minutes followed by PBS washings. Cells were either treated with 500µM IAA for 2 hours, or untreated. After the final wash, the medium was changed to phenol-red free imaging medium (with or without IAA) with SNAP blocking ligand 100 nM (New England Biolabs, S9106S) to prevent re-labeling of RAD21-SNAP$_{\mathrm{f}}$-V5 due to insufficiently removed cp-SNAP-JF549 dye which would otherwise appear as artefactual fluorescence recovery. iFRAP experiments were then performed on an LSM900 confocal microscope in a humidified incubation chamber at 37°C supplied with 5.5% $CO_2$. Bleaching was performed using the 561 nm laser at 100% power, scan speed 4, 1x crop factor, corresponding to a pixel dwell time of 1.02 µs. We imaged 4 frames, then bleached approximately 80% of the nucleus, and then imaged for a total of 250 frames, with 30 seconds of interval between frames. When selecting cells for iFRAP, we sought to enrich for G1/early S-phase cells. Briefly, we imaged a z-stack of 7.25 µm and visually evaluated the size and shape of the TetO and Anchor3 arrays, imaged with the same conditions adopted to perform locus tracking. We then excluded cells with apparent replication to enrich for G1/early S-phase. Furthermore, any cells that underwent mitosis during the movie, that extensively changed morphology, or where the cell movement was too high to allow for reliable quantification where excluded from subsequent analysis. iFRAP movies were analyzed as described below.

#### 2.9 iFRAP analysis

iFRAP was performed using a custom MATLAB (2021b; The MathWorks, Inc., Natick, MA, USA) script. Briefly, we drew a polygonal region of interest (ROI) on each of the bleached and unbleached ROI in cells undergoing FRAP. To account for cell movement, we checked and updated ROI positions every 25 frames (i.e. 12.5 mins); for the intervening frames, we updated the ROI positions using linear interpolation. The mean fluorescence intensity of each of bleached and unbleached region ($I_{\mathrm{bleached}}(t)$ and $I_{\mathrm{unbleached}}(t)$, respectively) were subsequently computed and stored for each frame and for each cell being analyzed.

To correct for photobleaching during image acquisition, we segmented unbleached nuclei in the field of view, and measured their mean intensities over time ($I_{\mathrm{photobleaching}}(t)$). To adjust $I_{\mathrm{bleached}}(t)$ and $I_{\mathrm{unbleached}}(t)$, we normalized these values by $I_{\mathrm{photobleaching}}(t) - I_{\mathrm{baseline}}$, where $I_{\mathrm{baseline}}$ is the background image intensity; we obtained $I_{\mathrm{baseline}}$ by drawing a ROI in a section not containing nuclei, and computing the mean intensity over time. Therefore, the final iFRAP signal (on a per-cell basis) was computed as:

$$\mathrm{iFRAP\ signal} = C \cdot \frac{I_{\mathrm{unbleached}}(t) - I_{\mathrm{bleached}}(t)}{I_{\mathrm{photobleaching}}(t) - I_{\mathrm{baseline}}}\,, \tag{1}$$

where $C$ is a constant measured from the mean pre-bleaching intensities of the unbleached ROI, and was used to normalize the iFRAP curves to their pre-bleaching levels.

Finally, we computed the mean iFRAP signal for the IAA and non-IAA treated cells, and fit the result to the following double-exponential function to obtain the cohesin dissociation rates from the chromosome:

$$f(t) = A_1 \exp(-k_1 t) + A_2 \exp(-k_2 t) + I_{\mathrm{offset}}\,. \tag{2}$$

The values $A_1$, $A_2$, $k_1$ and $k_2$ are fitting parameters, and $I_{\mathrm{offset}}$ is computed directly from the data and accounts for small differences in mean intensity between the ROIs in the bleached and unbleached regions (e.g. due to the presence of nucleoli, which partially exclude cohesins). We calculate $I_{\mathrm{offset}}$ from the difference in normalized mean intensities between the "bleached" and "unbleached" ROIs in the pre-bleaching frame. Fitting was performed in MATLAB using the functions *nlinfit* and *nlparci* to obtain the best fit parameters $A_1$, $A_2$, $k_1$ and $k_2$ as well as their 95% confidence intervals using the Jacobian matrix.

##### 2.10 Plotting of 3D distance histograms and MSD

Histograms of 3D distances between dots were computed using Numpy [16] and plotted with Matplotlib [17]. We pooled together all the registration corrected 3D distances from our dot pair trajectories, using only dot localizations that passed the quality control metrics described in section 2.5 (i.e. dots classified as both "unreplicated" and "good"). Bin sizes were set to "auto".

For the MSD plots, all particle trajectories were used. As with the 3D distance histogram, we only used dot pairs within a trajectory which passed the quality control metrics outlined above. The steady-state values shown in the figures were computed from the variance of the mean frame-to-frame particle displacements. To obtain localization-error corrected MSD curves, the localization error on the 3D distances between particles was first identified by the Bayesian approach described in section 5 and then subtracted from the raw MSDs. Plotting was done using Matplotlib [17].

##### 2.11 Calculating the unextruded fraction

In a partially extruded state, which is the generic conformation in everything but ΔRAD21, the average unextruded length serves as effective tether length. Under the Rouse model (section 6), the mean squared distance between the two tracked loci scales linearly in this effective tether length $L$ (eq. (33)). Assuming the effective tether length in the ΔRAD21 condition to be equal to the true genomic separation of the tracked loci of 515 kb, we can therefore estimate

$$L = \frac{J}{J_{\Delta\mathrm{RAD21}}} 515\,\mathrm{kb}\,, \tag{3}$$

where $J$ and $J_{\Delta\mathrm{RAD21}}$ denote the steady state variances in the condition under study and ΔRAD21, respectively. For the variances we use the values from **Fig. 2D** for ΔRAD21 and ΔCTCF; the corresponding variance for C36 is $0.32\,\mu\mathrm{m}^2$.

#### 3 Genomics experiments and analysis: ChIP-seq and Micro-C

##### 3.1 Library preperation

Chromatin immunoprecipitation coupled with sequencing (ChIP-seq) experiments were performed as described previously [2]. Briefly, ChIP assays in C65 mouse JM8.N4 mES cells were performed essentially as described [18] with minor modifications. Cells were cross-linked for 5 minutes at room temperature with 1% formaldehyde-containing medium; cross-linking was stopped by PBS-glycine (0.125 M final). Cells were washed twice with ice-cold PBS, scraped, centrifuged for 10 min at 4000 rpm, resuspended in cell lysis buffer (5 mM PIPES, pH 8.0, 85 mM KCl, and 0.5% NP-40, 1 ml/15 cm plate) and incubated for 10 min on ice. During the incubation, the lysates were repeatedly pipetted up and down every 5 min. Lysates were then centrifuged for 10 min at 4000 rpm. Nuclear pellets were resuspended in 6 volumes of sonication buffer (50 mM Tris-HCl, pH 8.1, 10 mM EDTA, 0.1% SDS), incubated on ice for 10 min, and sonicated to obtain DNA fragments below 2000 bp in length (Covaris S220 sonicator, 20% Duty factor, 200 cycles/burst, 150 peak incident power, 40 cycles of 20 seconds on and 40 seconds off). Sonicated lysates were cleared by centrifugation and 800 µg of chromatin were diluted in RIPA buffer (10 mM Tris-HCl, pH 8.0, 1 mM EDTA, 0.5 mM EGTA, 1% Triton X-100, 0.1% SDS, 0.1% Na-deoxycholate, 140 mM NaCl) to a final concentration of 0.8 µg/µl,precleared with Protein A sepharose (GE Healthcare) for 2 hours at 4° C and immunoprecipitated overnight with 8 µg of normal rabbit IgGs (ChromPure rabbit normal IgG; Jackson ImmunoResearch), anti-Smc1a (A300-055A; Bethyl Laboratories Inc.), or 1.6 µg of anti-CTCF antibodies (ab128873; Abcam). About 8% of the precleared chromatin was saved as input. Immunoprecipitated DNA was purified with the Qiagen QIAquick PCR Purification Kit, eluted in 60 µl of water and analyzed by qPCR together with 2% of the input chromatin prior to ChIP-seq library preparation (SYBR® Select Master Mix for CFX, ThermoFisher). Primer sequences were as follows:

| | | |
|---|---|---|
| Nanog_pos_for | GCAGAGCCACAGAAGGAATC | positive region |
| Nanog_pos_rev | TTGCCACCTGAAACCACATG | positive region |
| Nanog_neg_for | CTTTGGACAGGCATCGTAGC | negative region |
| Nanog_neg_rev | GCAGTATCTCACCATGCAGC | negative region |

ChIP-seq libraries were prepared using the Solexa rapid library protocol. Briefly, immunoprecipitated DNA or 50 ng input DNA was end-repaired, phosphorylated and adenylated in a single 50 µl reaction containing 31.5 µl DNA, 5 µl spike-in yeast DNA from MNase treated nucleosomes (10 ng/ml) [19] and 13.5 µl end-repair/3' A mix. The reaction was incubated in a thermal cycler for 15 min at 12° C, 15 min at 37° C, 20 min at 72° C, and held at 4° C. The ChIP-seq library preparation reaction mix is shown below:

| End-repair/3' A mix component | Final concentration | Catalog Number |
|---|---|---|
| 10X T4 DNA ligase buffer | 1X | NEB #B0202S |
| 10 mM dNTPs | 0.5 mM each | KAPA #KK1017 |
| 10 mM ATP | 0.25 mM | NEB #P0756S |
| 40% PEG 4000 | 2.5% | |
| 10 U/µl T4 PNK | 0.0025 U/µL | NEB #M0201S |
| 5 U/µl T4 DNA polymerase* | 0.0025 U/µL | Invitrogen #18005025 |
| 5 U/µl Taq DNA polymerase** | 0.0025 U/µL | Thermo #EP0401 |

* diluted 1:20 in 1x T4 DNA ligase buffer
** diluted 1:20 in 1X standard Taq buffer (NEB #B9014S)

To the reactions we added 4 µl water, 1 µl Illumina TruSeq adapters, 55 µl 2x Rapid DNA ligase buffer (Enzymatics #B101L) and 5 µl DNA ligase (Enzymatics #L6030-HC-L), and incubated for 15 min at 20° C. Ligations were cleaned up twice with AMPure XP beads (Agencourt #A63880) diluted 1:2 with 20% PEG, 1.25 M NaCl (first cleanup: 38 µl; beads eluted with 53 µl 10 mM Tris-HCl pH 8.0, 50 µl transferred to a new tube and added of 55 µl beads:PEG solution). Final elution volume was in 22 µl 10 mM Tris-HCl pH 8.0, 20 µl of which were transferred to a new tube and amplified by PCR (45 sec at 98° C; 14 cycles of 15 sec at 98° C and 10 sec at 60° C; 1 min at 72° C; hold at 4° C). The PCR reaction mix is shown below:

| PCR mix component | Final concentration | Catalog Number |
|---|---|---|
| 5X KAPA buffer | 1X | KAPA #KK2502 |
| 10 mM dNTPs | 0.3 mM each | KAPA #KK1017 |
| 5 µM TruSeq PCR primers* | 0.5 µM | |
| KAPA HS HIFI polymerase | 1 U | KAPA #KK2502 |
| Nuclease-free water to 30 µL | | |

* Primer1.0: AATGATACGGCGACCACCGAGATCTACACTCTTTCCCTACACGA;
* Primer2.0: CAAGCAGAAGACGGCATACGAGAT

PCR reactions were cleaned up once with 38 µl AMPure XP beads diluted 1:2 with 20% PEG, 1.25 M NaCl and eluted with 33 µl 10 mM Tris-HCl pH 8.0, 30 µl of which were transferred to a new tube. We assessed library quality and fragment size by qPCR and Fragment analyzer™, and sequenced 8-12 multiplexed libraries per lane on the Illumina HiSeq4000 sequencing platform (single end-reads, 50 bp long).

##### 3.2 Analysis

Input, IgG, CTCF and Smc1a ChIP-seq raw reads from C65 ESCs were quality-checked with FastQC and aligned onto the mouse genome (mm10 assembly) using Bowtie [20], allowing for two mismatches (-n 2) and no multiple alignments (-m 1). We ran bamCoverage [21] (–binSize 50 –normalizeTo1x 2150570000 –extendReads 250 –ignoreDuplicates -of bigwig) and normalized read numbers to 1x sequencing depth, obtaining read coverage per 50-bp bins across the whole genome (bigWig files). To display ChIP tracks we used CoolBox [22], transcripts were displayed with gencode M9 (GRCm38). CTCF motifs were identified with CTCFBSDB 2.0 [23,24] and the orientation determined using the CTCFBS Prediction Tool.

##### 3.3 Micro-C

To test if array insertion would perturb the *Fbn2* loop, we performed Micro-C in the C36 cell line. Micro-C was performed and analyzed as previously described [25]. The detailed procedure can be found here [26]. Micro-C data were plotted with Cooler [27] at a bin resolution of 4 kb. For inferring the sizes of cohesin loops *in vivo* directly from the Micro-C data, we used the method of log derivatives previously described [28].

#### 4 Loop extrusion and 3D polymer simulations

##### 4.1 Time steps and lattice set-up

We use a fixed-time-step Monte Carlo algorithm for 1D simulations as described in previous work [29], where each lattice site corresponded to 1 kb of DNA. The chromosome was defined as a lattice of $G$ = 101,700 sites consisting of 50 repeats of 2034 sites representing the 2034 kb region around the *Fbn2* TAD. Cohesins were comprised of two motor subunits that moved bidirectionally away from each other one lattice site at a time. We assumed that cohesins do not bypass each other upon encounter and that they cannot translocate past the first and the last lattice sites (i.e. cohesins do not "walk off" the chromosome).

To simulate the 2034 kb *Fbn2* locus and its surroundings, we manually annotated locations of CTCF boundaries using a combination of CTCF ChIP-seq and Micro-C data. Each annotated CTCF occupied one simulation lattice site and was given an estimated relative genome occupancy based on the strength of the TAD corner peak, as seen by Micro-C, and the CTCF ChIP-seq peak. Within each 2034 kb locus repeat, the CTCF locations and directions were as follows:

$$\text{Left-pointing CTCF positions} = [204, 241, 301, 469, 874, 884, 1129, \mathbf{1398}, 1508, 1984] \quad (4)$$

$$\text{Right-pointing CTCFs positions} = [50, 311, 469, \mathbf{889}, 949, 1422] \quad (5)$$

The CTCF sites corresponding to the *Fbn2* TAD boundaries are in bold above. The corresponding relative cohesin pausing probabilities for each of the CTCFs listed above were as follows:

$$\text{Relative cohesin stalling probability by left-pointing CTCFs} = [0.4, 0.4, 0.4, 0.4, 0.4, 0.4, 0.02, \mathbf{1.}, 0.34, 0.4] \times s \quad (6)$$

$$\text{Relative cohesin stalling probability by right-pointing CTCFs} = [0.4, 0.4, 0.4, \mathbf{1.}, 0.04, 0.4] \times s \quad (7)$$

The CTCF sites in our simulation were directional and could only stall cohesin movement if the cohesin motor subunit's extrusion direction was convergent with the direction of the CTCF. Upon a CTCF and cohesin motor subunit encounter, the cohesin motor subunit could be stalled by the CTCF with a probability $s$, or would pass the CTCF with a probability $(1-s)$ within the *Fbn2* TAD; for stalling probabilities by other CTCFs, refer to eqs. (6) and (7). Once a motor subunit was stalled by a CTCF site, no further movement of the subunit was allowed until the cohesin dissociated from the locus and re-associated elsewhere.

##### 4.2 Cohesin association and dissociation rates

All 1D loop extrusion simulations were performed with a fixed number of Loop Extrusion Factors (LEFs), determined by the ratio of chromosome length $G$ and cohesin separation $d$ (i.e., the inverse of the density). When a cohesin dissociated from the genome it immediately reloaded at another pair of adjacent lattice positions with uniform probability, provided that the lattice position was not occupied by existing cohesin motor subinits. The cohesin dissociation rate was governed by the LEF processivity $\lambda$ (i.e., the average length of DNA extruded by an unobstructed LEF before it dissociates) and any additional fold-increases in lifetime, $b$ that were gained by stabilization at a CTCF site.

##### 4.3 3D polymer simulations via OpenMM

Next, we coupled the 1D loop extrusion dynamics to a 3D polymer model, and performed molecular dynamics simulations using Polychrom [30], a package that wraps the molecular simulation toolkit OpenMM [31]. In this coupled model, cohesins act as harmonic bonds between two polymer monomers. These bonds are dynamically updated depending on the position of cohesins on the chromosome. Before coupling and 3D polymer steps, the 1D cohesin extrusion simulations were run for a total of 10,000 translocation steps in order to reach a steady-state value.

Chromosomal polymers were constructed of $G$=101,700 consecutive monomers bonded via the pairwise potential:

$$U_{\text{bonds}}(r) = \frac{k}{2}(r - b_o)^2 \,, \quad (8)$$

where $k = 2k_bT/\delta^2$ is the spring constant ($k_b$ being the Boltzmann constant, $T$ the temperature, and $\delta = 0.1$ monomers), $r = |r_i - r_j|$ is the spatial displacement between connected monomers, and $b_o = 1$ is the mean distance between monomers, in simulation units. Monomers bridged by a cohesin were held together by the same potential. To account for excluded volume interactions between monomers, we added a weak polynomial repulsive potential:

$$U_{\text{excl}}(r) = \frac{\epsilon_{\text{exc}}}{\epsilon_{\text{m}}}\left(\frac{r}{\sigma}r_{\text{m}}\right)^{12}\left(\left(\frac{r}{\sigma}r_{\text{m}}\right)^2 - 1\right) + \epsilon_{\text{exc}}\,, \tag{9}$$

defined for $r < \sigma = 1.05$, where $r_{\text{m}} = \sqrt{6/7}$,$\epsilon_{\text{m}} = 46656/823543$ and $\epsilon_{\text{exc}} = 1.5k_bT$.

At the start of each simulation, the polymer was initialized in a compact loop on a cubic lattice (polychrom.starting_conformations.grow_cubic()), with normally distributed velocities. The system thermostat was set with an error tolerance of 0.01 and the collision rate was set to 0.03. Simulations were performed with periodic boundary conditions, defined to maintain a DNA volume fraction of 20% per simulation volume. We calibrated the simulation time and distances to real time and distances using MSDs obtained from the imaging data. Using a maximum likelihood estimation procedure to infer the Rouse model parameters from our 3D polymer simulation MSDs and experimental MSDs (section 5), we inferred that each stored simulation time step corresponded to approximately 15-20 sec intervals and monomers corresponded to $\approx 19$ nm in diameter. Polymer conformations sampled at approximately 15-20 sec intervals corresponded to 5000 Langevin dynamics polymer integration steps using the conditions above.

We used a cohesin extrusion speed of $\approx 100 - 125$ bp/s (or 2 kb per 15-20 sec) on chromatin, unless otherwise specified. This extrusion rate was estimated by using the inferred cohesin processivity of 150 kb (see section 4.4) divided by the average residence time of cohesins measured experimentally of 20-25 min [2]. Simulations were run for a total of 12,096 saved conformations or equivalently, approximately 67 hours of simulated real time with sampling every 20 sec. From the resulting conformations, we generated 3D polymer structures, Micro-C-like contact maps, 3D distance distributions, and time-course data for the *Fbn2* locus dynamics.

##### 4.4 Identifying loop extrusion parameters for the C36 simulations

We needed to identify a total of 5 parameters. ① The cohesin lifetime on DNA, ② cohesin density and processivity, ③, cohesin extrusion rate, ④ probability that the R1 and L1 CTCF sites stall cohesin, ⑤ fold-increase of cohesin lifetime due to CTCF.

We used a combination of the following data as constraints to our model. ① We used iFRAP data for cohesin residence time **Fig. S13** which agreed with previously published results [2]. This identified that the residence lifetime of cohesin on DNA is approximately 20-30 mins. We used the timescale from cells after CTCF depletion, in order to obtain cohesin residence time in the absence of possible CTCF/cohesin stabilizing interactions. We note, however, that in S/G2 phase, cohesin has a longer residence time on chromatin than in G1 [32]; while we tried to identify G1/early S phase cells based on the replication state of the Fbn2 loop anchors, we cannot exclude the possibility of contamination by cells in late S/G2; thus, our iFRAP-measured residence time is likely an overestimate of the true cohesin residence time in G1. ② To obtain the cohesin processivity, we used the log-derivative based analysis of the Micro-C maps [28] in the 3-hours of CTCF depletion condition. We found that cohesins have a mean loop size of approximately ≈125 kb, which empirically translates to a processivity of ≈150 kb [28] due to interactions between LEFs, which shorten the average loop size below the processivity value [33] and a small bias in the estimation of the loop size via the log-derivative method (Polovnikov et al, Personal Communication). The cohesin density was obtained by performing 3D polymer simulations with varying cohesin separations (sep=[100, 150, 200, 250, 300, 350, 400] kb), while keeping the processivity fixed at 150 kb. The simulations were carried out with no CTCF sites to mimic the Micro-C for CTCF depletion after 3 hours. Computing the $P_c(s)$ curves for the simulations and experimental data, we minimized the sum of residuals between the $\log(P_c(s))$ curves, and found that the best-fit simulation had a mean cohesin density of 1/(300 kb). Then, to obtain the best estimate for the density of cohesins in C36 cells, we relied on the Western Blot results (**Fig. S6**) to correct for the fact that in the CTCF-AID tagged lines, the cohesin expression levels were ≈80% of the expression levels in C36. Thus, the average estimated density of cohesins in the C36 cell line was taken to be 1/(240 kb). ③ To obtain the cohesin extrusion rate, we used the relationship $v_{\text{extrusion}} = (\text{processivity})/(\text{residence time})$. Thus, using the processivity 150 kb with the the average cohesin lifetime on DNA of 25 mins (described above), we estimated the average extrusion rate of $v_{\text{extrusion}} = 125$ bp/s. ④-⑤ The probability that the R1 and L1 CTCF sites stall cohesin ($p_{\text{stall}}$) and the fold-increase (boost) of cohesin lifetime due to CTCF were estimated together ($b = \tau_{\text{complex}}/\tau_{\text{cohesin}}$). Via simulations, we found that $p_{\text{stall}} \cdot \tau_{\text{complex}}/\tau_{\text{cohesin}} =$ constant, where the constant factor will depend strongly on the desired mean tether length (i.e. fraction of unextruded DNA within the TAD). From 1D loop extrusion simulations, we found the empirical relationship between the unextruded fraction of DNA within the TAD and the values for $p_{\text{stall}}$ and $b$ (**Fig. S12A**), giving the relation $p_{\text{stall}} \cdot b \approx 0.5$. We thus swept various values of $p_{\text{stall}}$ (and its corresponding $b$). We identified that the best matching simulations according to their agreement to Micro-C contact probability decay versus genomic distance, loop fractions and loop lifetimes correspond to $b = 4$, and $p_{\text{stall}} = 1/8$ (**Fig. S12B**). The boost in cohesin residence times identified via these simulations by matching the looped fractions and loop lifetimes from BILD are also in agreement with the estimates obtained from the two-component fit to the cohesin residence time via iFRAP of between 2 to 5-fold increase in cohesin lifetime due to CTCF (**Fig. S13**).

### 5 Bayesian MSD fitting

For quantitative interpretation of our two-locus tracking data, we fit the obtained trajectories with a stationary, zero-mean Gaussian process. We outline the general approach in section 5.1 and describe details of the implementation in section 5.2.

#### 5.1 General approach

We denote by $\boldsymbol{x}_1(t)$ and $\boldsymbol{x}_2(t)$ the trajectories of the two loci, introduce the relative position $\boldsymbol{y}(t) \equiv \boldsymbol{x}_1(t) - \boldsymbol{x}_2(t)$, and make the following assumptions:

- $\boldsymbol{y}(t)$ is sampled from a Gaussian process: for any finite set $\{t_i\}_{i=1,\dots,T}$ the joint distribution $p\Big(y(t_1),\dots,y(t_T)\Big)$ is a multivariate Gaussian distribution.
- $\langle \boldsymbol{y}(t) \rangle = 0\, \forall t$
- the process is stationary; since we already assumed Gaussianity, a sufficient condition is that $\langle y(t+\Delta t)y(t)\rangle$ is independent of $t$.
- decaying correlations, i.e. $\gamma(\Delta t) \equiv \langle y(t+\Delta t)y(t)\rangle \to 0$ as $\Delta t \to \infty$.

Under these assumptions, we can readily calculate the MSD $\mu(\Delta t) \equiv \left\langle \Big(y(t+\Delta t) - y(t)\Big)^2 \right\rangle$ of the process:

$$\mu(\Delta t) = \left\langle y(t+\Delta t)^2 + y(t)^2 - 2y(t+\Delta t)y(t)\right\rangle = 2\left(\gamma(0) - \gamma(\Delta t)\right) , \tag{10}$$

which we reformulate as

$$\gamma(\Delta t) = \frac{1}{2}\left(\mu(\infty) - \mu(\Delta t)\right) , \tag{11}$$

using the assumption of decaying correlations to identify $\mu(\infty) = 2\gamma(0) \equiv 2\left\langle y^2 \right\rangle$. Since a Gaussian process is completely determined by its first and second order statistics (mean and covariance function; [34]) and we assumed the mean to be zero, this last equation completely defines the process in terms of the MSD.

Given a finite set of sampling times $\{t_i\}_{i=1,\dots,T}$ and a correspondingly sampled trajectory $\boldsymbol{Y} \equiv \Big(y(t_1),\dots,y(t_T)\Big)$, the above construction immediately allows us to write the joint probability for all data points given some MSD $\mu(\Delta t)$ as

$$p(\boldsymbol{Y} \mid \mu) = \left|2\pi\Sigma\right|^{-\frac{1}{2}} \exp\left(-\frac{1}{2}\boldsymbol{Y}\Sigma^{-1}\boldsymbol{Y}\right) , \tag{12}$$

with the covariance matrix $\Sigma_{ij} \equiv \gamma(t_i - t_j) = \frac{1}{2}\left(\mu(\infty) - \mu(t_i - t_j)\right)$. We can now use $p(\boldsymbol{Y} \mid \mu)$ as a likelihood function to perform Bayesian inference of the MSD from our given data. Typically we will choose a prior $p(\mu)$ with support only on some low dimensional subset of possible MSD functions $\mu(\Delta t)$, such that we can reformulate the problem as inference of a finite number of real parameters.

#### 5.2 Technical implementation

We implement the approach outlined in section 5.1 to fit MSDs of the general shape (32) (see section 6.1), plus localization error, to our experimental data (**Fig. S8D**).

We assume the localization error on a single spot to be Gaussian with standard deviation $\sigma$. We thus obtain a squared error of $2\sigma^2$ on the distance $y(t) \equiv x_1(t) - x_2(t)$ and therefore a total additive contribution of $4\sigma^2$ to the MSD $\mu(\Delta t) \equiv \left\langle (y(t+\Delta t) - y(t))^2\right\rangle$. We thus obtain the 3-parameter family of MSD functions

$$\mu(\Delta t; \sigma^2, \Gamma, J) := 4\sigma^2 + 2\Gamma\sqrt{\Delta t}\left(1 - \mathrm{e}^{-\frac{\tau}{\Delta t}}\right) + 2J\,\mathrm{erfc}\sqrt{\frac{\tau}{\Delta t}} , \tag{13}$$

where $\tau \equiv \frac{1}{\pi}\left(\frac{J}{\Gamma}\right)^2$.

Since we expect the localization error to be different and independent along the three spatial dimensions (resolution is worse in the axial direction), we perform the fit separately in each dimension, subject to the constraints $\Gamma_x = \Gamma_y = \Gamma_z \equiv \frac{1}{3}\Gamma$ and $J_x = J_y = J_z \equiv \frac{1}{3}J$, i.e. the polymer dynamics should be the same in all dimensions. This ultimately leaves us with the 5 parameters

$$\sigma_x^2\,,\ \sigma_y^2\,,\ \sigma_z^2\,,\ \Gamma\,,\ J\,. \tag{14}$$

Since these 5 parameters are all positive and have units (i.e. their numerical scale is *a priori* undetermined), we choose a log-flat prior for each of them. Technically we implement this by defining the parameter vector $\boldsymbol{\theta} \equiv \left(\log\sigma_x^2, \log\sigma_y^2, \log\sigma_z^2, \log\Gamma, \log J\right)$ and choosing the prior $p(\boldsymbol{\theta}) \propto 1$ (flat prior in log-space). Via eq. (13) we associate with each $\boldsymbol{\theta}$ the MSDs $\mu_i(\Delta t; \boldsymbol{\theta})$ for all three spatial dimensions $i = x, y, z$, and then use eq. (12) to calculate the likelihood

$$p(\boldsymbol{Y} \mid \boldsymbol{\theta}) = p(\boldsymbol{Y}^{(x)} \mid \mu_x) p(\boldsymbol{Y}^{(y)} \mid \mu_y) p(\boldsymbol{Y}^{(z)} \mid \mu_z) \tag{15}$$

given a single trajectory $\boldsymbol{Y}$. Finally, the likelihood for a given set of parameters $\boldsymbol{\theta}$, given a set of trajectories $\mathcal{Y} \equiv \{\boldsymbol{Y}_n\}_{n=1,\ldots,N}$ is the product over the individual trajectory likelihoods:

$$p(\mathcal{Y} \mid \boldsymbol{\theta}) = \prod_n p(\boldsymbol{Y}_n \mid \boldsymbol{\theta}) \,. \tag{16}$$

We thus obtain, up to a normalization constant, an analytical expression for the posterior $p\left(\boldsymbol{\theta} \mid \mathcal{Y}\right) \propto p\left(\mathcal{Y} \mid \boldsymbol{\theta}\right) p(\boldsymbol{\theta})$. This we maximize numerically, using the Nelder-Mead (simplex) method, followed by gradient ascent.

Errors on this point estimate are given by the marginal posterior standard deviations, which we evaluate numerically by MCMC and generally find to be below 2% (**Fig. S8E**). To that end, we first obtain a guess for the shape of the posterior peak by determining the points where $\log \frac{p(\boldsymbol{\theta}_{\mathsf{MAP}} \mid \mathcal{Y})}{p(\boldsymbol{\theta} \mid \mathcal{Y})} = \frac{1}{2}\chi^2_{5,0.95} \approx 5.85$ along each of the 5 parameter axes. We then initialize an MCMC chain at the MAP estimate and set its step size as the width of the peak estimated in each direction. This guarantees good sampling of the posterior peak, skipping the burn-in period of the MCMC (**Fig. S8E**).

### 6 The Rouse model

Throughout this work we use the Rouse polymer model [35–37] in various different versions (continuous or discrete, finite or infinite chain). While this is a well-known model, some of the specific results (e.g. MSD for relative position of two loci) are not readily available in the literature; we therefore provide a complete derivation. We start out by deriving the MSD for two loci on an infinite continuous chain in section 6.1, which serves as the basis for our physical interpretation of the data (see also section 5). Our inference approach (BILD), however, can work only with a finite, discrete chain; we derive the associated likelihood function in section 6.2. We then show in section 6.3 that the discrete model indeed provides a good approximation to the continuous one and characterize the two finite-size effects controlling that approximation.

#### 6.1 MSD for the relative position of two loci on an infinite continuous Rouse polymer

We use $s \in \mathbb{R}$ as coordinate along the backbone of the polymer and write $x(s,t)$ for its time-dependent conformation. The polymer follows the overdamped Langevin equation

$$\gamma \dot{x}(s,t) = -\kappa \partial_s^2 x(s,t) + \xi(s,t)\,, \qquad \left\langle \xi(s,t)\xi(s',t')\right\rangle = 2\gamma k_{\mathsf{B}} T \delta(s-s')\delta(t-t')\,, \tag{17}$$

where $\xi(s,t)$ is a zero-mean Gaussian field representing the thermal noise whose amplitude is given by the Einstein relation (fluctuation dissipation theorem). For convenience we introduce $D \equiv \frac{k_{\mathsf{B}} T}{\gamma}$.

Equation (17) is a heat equation, whose solution is given by a Weierstrass transform of the noise:

$$x(s,t) = \int_{\mathbb{R}} \mathrm{d}\sigma \int_0^t \mathrm{d}\tau \, \frac{\mathrm{e}^{-\frac{\gamma(\sigma-s)^2}{4\kappa\Delta t}}}{\sqrt{4\pi\gamma\kappa(t-\tau)}} \xi(\sigma,\tau) = \frac{1}{\gamma}\int_{\mathbb{R}} \mathrm{d}\sigma \int_0^t \mathrm{d}\tau \, N\left(\sigma; s, \frac{2\kappa}{\gamma}(t-\tau)\right) \xi(\sigma,\tau)\,, \tag{18}$$

where $N\left(x; A, B\right) \equiv (2\pi B)^{-\frac{1}{2}} \exp\left(-\frac{(x-A)^2}{2B}\right)$ is a Gaussian and we assumed the collapsed initial condition $x(s,0) = 0\,\forall s$. This solution allows us to calculate the full covariance structure of $x(s,t)$. Without loss of generality we assume $t' \equiv t + \Delta t \geq t$, such that we can write

$$\begin{aligned}
\left\langle x(s',t')x(s,t)\right\rangle &= \frac{1}{\gamma^2}\int \mathrm{d}\sigma'\mathrm{d}\sigma \int_0^{t'} \mathrm{d}\tau \int_0^t \mathrm{d}\tau' \, N\left(\sigma'; s', \frac{2\kappa}{\gamma}(t'-\tau')\right) N\left(\sigma; s, \frac{2\kappa}{\gamma}(t-\tau)\right) \left\langle \xi(\sigma',\tau')\xi(\sigma,\tau)\right\rangle &(19)\\
&= 2D\int \mathrm{d}\sigma \int_0^t \mathrm{d}\tau \, N\left(-\sigma; s'-s, \frac{2\kappa}{\gamma}(t-\tau)\right) N\left(\sigma; 0, \frac{2\kappa}{\gamma}(t'-\tau)\right) &(20)\\
&= 2D\int_0^t \mathrm{d}\tau \, N\left(0; s'-s, \frac{2\kappa}{\gamma}(t'+t-2\tau)\right) &(21)\\
&= 2D\int_0^t \frac{\sqrt{\gamma}\,\mathrm{d}\tau}{\sqrt{8\pi\kappa t\left(\frac{\Delta t}{2t}+1-\frac{\tau}{t}\right)}} \exp\left(-\frac{\gamma\Delta s^2}{8\kappa t\left(\frac{\Delta t}{2t}+1-\frac{\tau}{t}\right)}\right)\,, &(22)
\end{aligned}$$

where we first use the noise correlations $\langle \xi(\sigma',\tau')\xi(\sigma,\tau)\rangle = 2\gamma k_{\mathsf{B}} T \delta(\sigma'-\sigma)\delta(\tau'-\tau)$ and introduce $D = \frac{k_{\mathsf{B}} T}{\gamma}$, then transform $\sigma \leftarrow s-\sigma$, and finally execute the integral over $\sigma$, which is a convolution of two Gaussians. In the last step, we expand the expression for the Gaussian and introduce $\Delta s \equiv s' - s$ and $\Delta t \equiv t' - t$. We now substitute $z \equiv 1 - \frac{\tau}{t}$ and employ the incomplete Gamma function $\Gamma(\nu, z) \equiv \int_z^\infty t^{\nu-1} \mathrm{e}^{-t} \mathrm{d}t$ to rewrite the resulting integral as

$$\begin{aligned} \left\langle x(s',t')x(s,t) \right\rangle &= 2D \int_0^1 \frac{\sqrt{\gamma t}\,\mathrm{d}z}{\sqrt{8\pi\kappa\left(z + \frac{\Delta t}{2t}\right)}} \exp\left(-\frac{\gamma \Delta s^2}{8\kappa t\left(z + \frac{\Delta t}{2t}\right)}\right) & (23) \\ &= 2D\,\frac{\gamma\,|\Delta s|}{8\kappa\sqrt{\pi}} \left[\Gamma\left(-\frac{1}{2}, \frac{\gamma\Delta s^2}{4\kappa(2t+\Delta t)}\right) - \Gamma\left(-\frac{1}{2}, \frac{\gamma\Delta s^2}{4\kappa\Delta t}\right)\right]. & (24) \end{aligned}$$

Ultimately we are interested in the equilibrium behavior of the chain. We therefore aim to expand eq. (24) for large $t$, while holding $\Delta t$ and $\Delta s$ constant. To that end we note the following representation of the incomplete Gamma function*: $\Gamma\left(-\frac{1}{2}, z\right) = \frac{2\mathrm{e}^{-z}}{\sqrt{z}} - 2\sqrt{\pi}\,\mathrm{erfc}\,\sqrt{z}$, which for small $z$ expands as $\Gamma\left(-\frac{1}{2}, z\right) = \frac{2}{\sqrt{z}} - 2\sqrt{\pi} + \mathcal{O}(\sqrt{z})$. Expanding the first term and substituting the exact expression for the second one we find

$$\begin{aligned} \left\langle x(s',t')x(s,t) \right\rangle &= 2D\left[\sqrt{\frac{\gamma t}{2\pi\kappa}} - \sqrt{\frac{\gamma\Delta t}{4\pi\kappa}} \exp\left(-\frac{\gamma\Delta s^2}{4\kappa\Delta t}\right) - \frac{\gamma\,|\Delta s|}{4\kappa}\,\mathrm{erf}\,\sqrt{\frac{\gamma\Delta s^2}{4\kappa\Delta t}}\right] + \mathcal{O}\left(\sqrt{\frac{\Delta t}{t}}, \sqrt{\frac{\gamma\Delta s^2}{\kappa t}}\right) & (25) \\ &\equiv A\sqrt{t} + C^0(\Delta s, \Delta t) + \mathcal{R}\,, & (26) \end{aligned}$$

Note that the first term $A\sqrt{t}$ describes the continuing expansion of the chain, and accordingly diverges as $t \to \infty$. This means that the system as a whole actually never equilibrates, agreeing with the intuition that an infinite polymer with an initially completely collapsed conformation would not reach a steady state, but just keep expanding (note that since the chain is infinitely long, there is no whole coil diffusion at long times). However, for quantities that do not depend on the absolute position of the chain (like MSDs) this term drops out, such that they do reach a steady state on time scales $t \gg \max\left(\Delta t, \frac{\gamma}{\kappa}\Delta s^2\right)$, where $\Delta t$ and $\Delta s$ are the time scale and chain length relevant to the subsystem under study. This is exemplified by the specific quantity we are interested in here, namely the MSD for the relative position of two loci on the chain, as shown in the following.

We study the dynamics of two loci on the chain, situated at $s = s_1$ and $s = s_2 \equiv s_1 + \Delta s$. Writing $y(t) \equiv x(s_1, t) - x(s_2, t)$ we are interested in the MSD $\mu(\Delta t) \equiv \left\langle (y(t+\Delta t) - y(t))^2 \right\rangle$. Using eq. (25) we find

$$\begin{aligned} \mu(\Delta t) = &\left\langle (x(s_1, t+\Delta t) - x(s_2, t+\Delta t) - x(s_1,t) + x(s_2,t))^2 \right\rangle & (27) \\ = &\Big\langle x^2(s_1,t+\Delta t) - 2x(s_1,t+\Delta t)x(s_2,t+\Delta t) - 2x(s_1,t+\Delta t)x(s_1,t) + 2x(s_1,t+\Delta t)x(s_2,t) \\ &+ x^2(s_2,t+\Delta t) + 2x(s_2,t+\Delta t)x(s_1,t) - 2x(s_2,t+\Delta t)x(s_2,t) \\ &+ x^2(s_1,t) - 2x(s_1,t)x(s_2,t) \\ &+ x^2(s_2,t)\Big\rangle & (28) \\ = &A\sqrt{t+\Delta t} - 2A\sqrt{t+\Delta t} - 2C^0(\Delta s,0) - 2A\sqrt{t} - 2C^0(0,\Delta t) + 2A\sqrt{t} + 2C^0(\Delta s,\Delta t) \\ &+ A\sqrt{t+\Delta t} + 2A\sqrt{t} + 2C^0(\Delta s,\Delta t) - 2A\sqrt{t} - 2C^0(0,\Delta t) \\ &+ A\sqrt{t} - 2A\sqrt{t} - 2C^0(\Delta s,0) \\ &+ A\sqrt{t} + \mathcal{R} & (29) \\ = &4\left[C^0(\Delta s,\Delta t) - C^0(\Delta s,0) - C^0(0,\Delta t)\right] + \mathcal{R}\,. & (30) \end{aligned}$$

Now that the explicit dependence on $t$ is only in the remainder term $\mathcal{R} \equiv \mathcal{O}\left(\sqrt{\frac{\Delta t}{t}}, \sqrt{\frac{\gamma\Delta s^2}{\kappa t}}\right)$, we can actually take the limit $t \to \infty$, i.e. wait until the subchain we study is equilibrated. We can then expand the non-vanishing terms of eq. (30) to obtain

$$\mu(\Delta t) = 2D\left[\sqrt{\frac{4\gamma\Delta t}{\pi\kappa}}\left(1 - \mathrm{e}^{-\frac{\gamma\Delta s^2}{4\kappa\Delta t}}\right) + \frac{\gamma\,|\Delta s|}{\kappa}\,\mathrm{erfc}\,\sqrt{\frac{\gamma\Delta s^2}{4\kappa\Delta t}}\right], \tag{31}$$

which we rewrite as

$$\mu(\Delta t) = 2\Gamma\sqrt{\Delta t}\left(1 - \mathrm{e}^{-\frac{\tau}{\Delta t}}\right) + 2J\,\mathrm{erfc}\,\sqrt{\frac{\tau}{\Delta t}}\,, \tag{32}$$

with the phenomenological constants $\Gamma \equiv 2D\sqrt{\frac{\gamma}{\pi\kappa}}$, which is also the scaling prefactor in the MSD $\Gamma\sqrt{\Delta t}$ of a *single* locus on the polymer (calculation not shown); $J \equiv \frac{D\gamma}{\kappa}\,|\Delta s| = \left\langle y^2 \right\rangle$, the steady state variance of the distance between the two loci; and the crossover time scale $\tau \equiv \frac{\gamma\Delta s^2}{4\kappa} = \frac{1}{\pi}\left(\frac{J}{\Gamma}\right)^2$.

*http://functions.wolfram.com/06.06.03.0006.01; also easily checked by differentiation

To give some intuition for eq. (32) we study the long and short lag time limits. At short lag times $\Delta t \ll \tau$ we have $\mu(\Delta t) \approx 2\Gamma\sqrt{\Delta t} = 2\mu_{\text{single locus}}(\Delta t)$. In this regime the two loci do not yet feel the finite tether between them and effectively behave as if they were on independent polymers, thus their MSDs simply add and we obtain the familiar $\sqrt{\Delta t}$ scaling, with a doubled prefactor. For long lag times, we find that $\mu(\Delta t) \approx 2J =$ const., consistent with the intuition that the system reaches steady state and we simply sample from the associated steady state distance distribution. These two asymptotes coincide for $\Delta t = \left(\frac{J}{\Gamma}\right)^2 \equiv \pi\tau$, though of course the crossover is somewhat washed out by the exponential weights on the individual terms (**Fig. S8A**).

Note that the derivation in this section assumed a single spatial dimension, $d = 1$. In higher spatial dimensions in principle all the microscopic constants ($\gamma$, $\kappa$, $D$) become second rank tensors. Assuming isotropicity these are all proportional to identity, such that the full Langevin equation (17) decouples into $d$ one-dimensional equations exactly like the one we studied here. Thus, ultimately the only modification is an additional factor $d$ whenever spatial indices are contracted (e.g. $\mu_d(\Delta t) \equiv \left\langle \sum_i \left(y_i(t+\Delta t) - y_i(t)\right)^2 \right\rangle = d\mu_1(\Delta t)$). This does of course not modify the phenomenological shape (32) of the MSD, but just the microscopic representation of the constants:

$$\Gamma = 2dD\sqrt{\frac{\gamma}{\pi\kappa}}\,, \qquad J = \frac{dD\gamma}{\kappa}\left|\Delta s\right|\,. \tag{33}$$

To summarize, in this section we derived the analytical expression (32) for the MSD of the relative position of two loci on a continuous, infinite Rouse polymer, which we then fit to the experimental data using the approach of section 5.

#### 6.2 Likelihood function of a multi-state discrete Rouse model

The model underlying our inference is a two-state discrete Rouse model. We define a *looping profile* $\theta(t)$ as the indicator function of the looped state, i.e. $\theta(t) = 1$ whenever the looped state is present, while $\theta(t) = 0$ for times where the looped state is absent. Using the exact solution to the Rouse model, we are then able to calculate the likelihood function $p\left(y \mid \theta\right)$ analytically. For computational efficiency (see below) we choose to perform the likelihood calculation via the recursive Kalman filter equations.

The discrete Rouse model is defined as a number $N$ of point particles (monomers), connected by springs with spring constant $k$, suspended in solvent with friction coefficient $\gamma$ (we assume overdamped motion), and driven by thermal noise such that a free monomer would move diffusively with diffusivity $D$, for which the Einstein relation (fluctuation-dissipation theorem) dictates $D = \frac{k_\text{B}T}{\gamma}$. Consequently, the equations of motion are

$$\gamma\dot{\boldsymbol{x}}(t) = -kB(\theta(t))\boldsymbol{x}(t) + \boldsymbol{\xi}(t)\,, \qquad \left\langle \boldsymbol{\xi}(t) \otimes \boldsymbol{\xi}(t')\right\rangle = 2D\gamma^2\mathbb{1}_N\delta(t-t')\,, \tag{34}$$

where $\boldsymbol{x}(t)$ is the $N$-dimensional vector of monomer positions, $\mathbb{1}_N$ denotes the associated $N$-dimensional identity matrix, $B(\theta(t))$ is the connectivity matrix pertaining to the state $\theta(t)$, and for simplicity we omit the additional spatial index $i = x, y, z$. The connectivity matrix $B(\theta)$ is constructed iteratively: starting from the zero matrix, we insert a bond between monomers $i$ and $j$ by updating $B_{ii} \leftarrow B_{ii} + 1$, $B_{jj} \leftarrow B_{jj} + 1$, $B_{ij} \leftarrow B_{ij} - 1$, and $B_{ji} \leftarrow B_{ji} - 1$. For a linear chain (our model for the unlooped state $\theta = 0$), this yields

$$B(0) = \begin{pmatrix} 1 & -1 & & & \\ -1 & 2 & -1 & & \\ & & \ddots & & \\ & & -1 & 2 & -1 \\ & & & -1 & 1 \end{pmatrix}\,, \tag{35}$$

while for the looped state we insert an additional bond between the two monomers corresponding to the CTCF sites (c.f. section 7.2), thus obtaining a "looped" connectivity matrix $B(1)$.

Since $\theta(t)$ is piecewise constant, we can immediately give the solution of eq. (34) as

$$\boldsymbol{x}(t+\Delta t) = \mathrm{e}^{-\frac{k}{\gamma}\Delta t B(\theta)}\boldsymbol{x}(t) + \frac{1}{\gamma}\int_0^{\Delta t} \mathrm{d}\tau\, e^{-\frac{k}{\gamma}\tau B(\theta)}\boldsymbol{\xi}(t+\Delta t-\tau) \tag{36}$$

$$\equiv A_{\theta,\Delta t}\boldsymbol{x}(t) + \boldsymbol{\eta}_{\theta,\Delta t}(t)\,, \tag{37}$$

where we assume $\theta(t) \equiv \theta =$ const. during the interval $[t, t+\Delta t)$. With the second line we transition to a discrete time picture, adding Gaussian noise to the incrementally relaxed conformation $A_{\theta,\Delta t}\boldsymbol{x}(t)$ at each time step. Thus, if we start from an initially Gaussian ensemble $p_t(\boldsymbol{x}) = N\left(\boldsymbol{x}; \boldsymbol{\mu}_t, \Sigma_t\right)$, then the propagated ensemble $p_{t+\Delta t}(\boldsymbol{x})$ will also be Gaussian with mean $\boldsymbol{\mu}_{t+\Delta t}$ and variance $\Sigma_{t+\Delta t}$, for which eq. (37) gives

$$\mu_{t+\Delta t} = A_{\theta,\Delta t}\mu_t \tag{38}$$

$$\Sigma_{t+\Delta t} = A_{\theta,\Delta t}\Sigma_t A^T_{\theta,\Delta t} + S_{\theta,\Delta t}\,, \tag{39}$$

where $S_{\theta,\Delta t} \equiv 2D \int_0^{\Delta t} \mathrm{d}\tau \, \mathrm{e}^{-2\frac{k}{\gamma}\tau B(\theta)}$ is the covariance matrix of $\boldsymbol{\eta}_{\theta,\Delta t}$.

If $B$ is positive definite, all the eigenvalues of $A_{\Delta t}$ will decay exponentially with $\Delta t$, such that we can take the limit of eqs. (38) and (39) as $\Delta t \to \infty$ to find the steady state

$$\mu_\infty = 0 \tag{40}$$

$$\Sigma_{\theta,\infty} = S_{\theta,\infty} = \frac{D\gamma}{k} B^{-1}(\theta) \,. \tag{41}$$

Note that $B > 0$ is *not* the generic case. Specifically, the connectivity matrix for the linear chain (35) has rank $N-1$ and is thus only positive semi-definite and not invertible. Physically this is due to the unconstrained center of mass diffusion of the chain, which means that the system does not have a steady state (the coil will just keep diffusing). Practically, however, we are not interested in the center of mass position of the coil, but only in projected quantities $\boldsymbol{w}^T \boldsymbol{x}$ with $\sum_i w_i = 0$, which are invariant under translation of the full conformation (specifically, for the relative position of two monomers $i$ and $j$, we have $w_k = \delta_{ik} - \delta_{jk}$). These degrees of freedom do reach steady state, and we can obtain a corresponding steady state ensemble by pinning the chain to a fixed point in space. We do so by introducing a modified connectivity matrix $\tilde{B}$ with

$$\tilde{B}_{00} := B_{00} + 1 \,, \qquad \tilde{B}_{ij} := B_{ij} \, \forall (i \neq 0) \vee (j \neq 0) \,, \tag{42}$$

effectively inserting an additional bond between the first monomer and the origin. We emphasize that we introduce this additional bond *only* to find the steady state ensemble, and remove it when we study dynamics.

Equation (37) describes a linear model driven by Gaussian noise. As such, we can calculate the trajectory likelihood $p\,(y \mid \theta)$ efficiently via the Kalman filter equations [34, 38]. We start out by defining a set of observation times $\{t_n\}$ and introduce the discrete time notation $\boldsymbol{x}_n \equiv \boldsymbol{x}(t_n)$. We then define transition matrices $F_n$ and process noise correlations $Q_n$ by assembling the contributions dictated by the looping profile. If, for example, $\theta(t)$ switches from 1 (looped) to 0 (unlooped) at $\tau \in [t_{n-1}, t_n]$, we set

$$F_n = A_{0,t_n-\tau} A_{1,\tau-t_{n-1}} = \mathrm{e}^{-\frac{k}{\gamma}(t_n-\tau)B(0)} \mathrm{e}^{-\frac{k}{\gamma}(\tau-t_{n-1})B(1)} \tag{43}$$

$$Q_n = S_{0,t_n-\tau} + A_{0,t_n-\tau} S_{1,\tau-t_{n-1}} \left(A_{0,t_n-\tau}\right)^T \tag{44}$$

$$= 2D \left[ \int_\tau^{t_n} \mathrm{d}\tau' \, \mathrm{e}^{-2\frac{k}{\gamma}(t_n-\tau')B(0)} + \int_{t_{n-1}}^{\tau} \mathrm{d}\tau' \, \mathrm{e}^{-\frac{k}{\gamma}(t_{n+1}-\tau)B(0)} \mathrm{e}^{-2\frac{k}{\gamma}(\tau-\tau')B(1)} \mathrm{e}^{-\frac{k}{\gamma}(t_{n+1}-\tau)B^T(0)} \right] . \tag{45}$$

At each time point we observe the relative position of the two loci $i$ and $j$ with a given localization error $\sigma$, which we capture by the observation model

$$y_n = \boldsymbol{w}^T \boldsymbol{x}_n + \epsilon_n \,, \tag{46}$$

where $w_k = \delta_{ik} - \delta_{jk}$ as mentioned above and $\epsilon_n$ are zero-mean Gaussian localization errors with covariance $\langle \epsilon_m \epsilon_n \rangle = \sigma^2 \delta_{mn}$. The corresponding measurement likelihood is given by

$$p(y_n \mid \boldsymbol{x}_n) = N\left(y_n; \boldsymbol{w}^T \boldsymbol{x}_n, \sigma^2\right) \,. \tag{47}$$

Our goal is now to calculate the trajectory likelihood

$$p(y \mid \theta) = \prod_n p\left(y_n \mid y_{n-1}, \ldots, y_1, \theta\right) \,, \tag{48}$$

where the individual factors can be calculated as

$$p\left(y_n \mid y_{n-1}, \ldots, y_1, \theta\right) = \int_{-\infty}^{\infty} \mathrm{d}\boldsymbol{x}_n p(y_n \mid \boldsymbol{x}_n) p(\boldsymbol{x}_n \mid y_{n-1}, \ldots, y_1, \theta) \,. \tag{49}$$

Now, given a predicted ensemble $p(\boldsymbol{x}_n \mid y_{n-1}, \ldots, y_1, \theta) = N\left(\boldsymbol{x}_n; \boldsymbol{\mu}_n, \Sigma_n\right)$ we obtain

$$p\left(y_n \mid y_{n-1}, \ldots, y_1, \theta\right) = N\left(y_n; \boldsymbol{w}^T \boldsymbol{\mu}_n, \boldsymbol{w}^T \Sigma_n \boldsymbol{w} + \sigma^2\right) \,, \tag{50}$$

where the conditioning on the previous observations and $\theta$ is subsumed in $\boldsymbol{\mu}_n$ and $\Sigma_n$. Moreover, we calculate the optimal Kalman gain

$$\boldsymbol{k} = \frac{\Sigma_n \boldsymbol{w}}{\boldsymbol{w}^T \Sigma_n \boldsymbol{w} + \sigma^2} \,, \tag{51}$$

allowing us to update our knowledge of the state ensemble expressed by the posterior distribution

$$p(\boldsymbol{x}_n \mid y_n, \ldots, y_1, \theta) = N\left(\boldsymbol{x}_n; \boldsymbol{\mu}_n^{\text{post}}, \Sigma_n^{\text{post}}\right) \tag{52}$$

with

$$\boldsymbol{\mu}_n^{\text{post}} = \boldsymbol{\mu}_n + \boldsymbol{k}\left(y_n - \boldsymbol{w}^T\boldsymbol{\mu}_n\right) \tag{53}$$

$$\Sigma_n^{\text{post}} = \left(\mathbb{1}_N - \boldsymbol{k}\boldsymbol{w}^T\right)\Sigma_n\,, \tag{54}$$

which we then propagate under eqs. (43) and (44) to find the new predicted ensemble statistics

$$\boldsymbol{\mu}_{n+1} = F_{n+1}\boldsymbol{\mu}_i^{\text{post}} \tag{55}$$

$$\Sigma_{n+1} = F_{n+1}\Sigma_n^{\text{post}}F_{n+1}^T + Q_{n+1} \tag{56}$$

and restart the scheme with the next observation $y_{n+1}$. We initialize this recursive scheme with the steady state associated with $\theta(0)$, given by eqs. (40) and (41) as

$$\boldsymbol{\mu}_1 = 0 \tag{57}$$

$$\Sigma_1 = \frac{D\gamma}{k}\tilde{B}^{-1}(\theta(0))\,, \tag{58}$$

where $\tilde{B}$ indicates that we pin the chain in space to guarantee the existence of a steady state, as described in eq. (42), and take the notation $B_{\theta(0)}$ to mean the free chain connectivity matrix $B$ of eq. (35) if $\theta(0) = 0$ and the looped connectivity matrix $B_L$ if $\theta(0) = 1$.

Finally, we add up all the contributions (50) to find the total likelihood

$$p(y \mid \theta) = \prod_n p\left(y_n \mid y_{n-1}, \dots, y_1, \theta\right) = \prod_n N\left(y_n; \boldsymbol{w}^T\boldsymbol{\mu}_n, \boldsymbol{w}^T\Sigma_n\boldsymbol{w} + \sigma^2\right)\,, \tag{59}$$

which is the center piece of our Bayesian inference framework, described in detail in the next section.

Note that instead of using the Kalman filter to calculate the Rouse likelihood in eq. (59), we could also have exploited the fact that the full ensemble of state trajectories $\boldsymbol{X} \equiv (\boldsymbol{x}_1, \dots, \boldsymbol{x}_T)$ for a given looping profile $\theta(t)$ is always Gaussian, and just evaluate the likelihood directly. This however would require assembling the full $TN \times TN$ covariance matrix of that distribution and therefore scales quadratically in the trajectory length $T$. The calculation via the Kalman filter on the other hand clearly scales only linearly in $T$ and is thus computationally more efficient.

While in principle switches in a looping profile can happen at any time $s \in (0, T)$, we constrain them to occur only at integer multiples of the movie frame time $\Delta t$. This allows us to precalculate the matrices

$$A_{\theta,\Delta t} \equiv \mathrm{e}^{-\frac{k}{\gamma}\Delta t B(\theta)} \tag{60}$$

$$S_{\theta,\Delta t} \equiv 2D\int_0^{\Delta t} \mathrm{d}\tau\, \mathrm{e}^{-2\frac{k}{\gamma}\tau B(\theta)} \tag{61}$$

for $\theta \in \{0, 1\}$ and thus also greatly contributes to computational feasibility.

##### 6.3 MSD for the relative position of two loci on an infinite discrete Rouse polymer

Following the argument presented in section 5.1, as a Gaussian process the trajectories generated by the Rouse model are fully determined by their MSD [34]. We therefore study how well the discrete Rouse model approximates the continuous one by comparing their MSDs, assuming an infinite chain in both cases. Having found a closed analytical expression for the continuous model in section 6.1, we here provide the asymptotic behavior of the discrete one. Unsurprisingly we find perfect agreement in the asymptotics for the Rouse scaling and equilibrium regimes. On length scales shorter than the monomer size, the discrete model deviates from the continuous one into an unphysical regime of free monomer diffusion. We avoid this finite size effect by always choosing $\frac{\gamma}{k} < \Delta t_{\text{frame}}$ (see below). Finally, a second finite size effect is introduced if the chain in which our two loci are embedded is finite. We study this numerically in **Fig. S8C** and minimize the introduced error by always having the embedding chain be at least three times as long as the polymer between the two loci.

We extend the model in eq. (34) to an infinite chain of discrete monomers, such that the connectivity matrix $B$ becomes a Toeplitz operator with $B_{nn} = 2$, $B_{n,n\pm1} = -1$, and all other $B_{mn} = 0$. As such it is diagonalized by the Fourier transform, with eigenvalues given by the discrete cosine transform (DCTIII) of the diagonal entries:

$$B_{nm} = \int_{-\pi}^{\pi} \mathrm{d}\omega\, e^*(n,\omega)e(m,\omega)\lambda(\omega)\,, \tag{62}$$

with $\lambda(\omega) = 2(1 - \cos\omega)$, the Fourier basis $e(n,\omega) = \frac{1}{\sqrt{2\pi}}\mathrm{e}^{\mathrm{i}n\omega}$, and $^*$ denoting complex conjugation. Note that while technically $B$ is still not invertible (since $\lambda(0) = 0$), the infinite chain takes infinite time to reach the coil diffusion regime. Mathematically this has the consequence that $\{\omega \in [-\pi, \pi] : \lambda(\omega) = 0\} = \{0\} \subset [-\pi, \pi]$ is a subset of measure zero, such that the integrals below are well-defined, allowing us to mostly ignore this problem.

From (in that order) eqs. (10), (37), (41) and (62) we find the MSD $\mu(\Delta t)$ of $y(t) \equiv \boldsymbol{w}^T \boldsymbol{x}(t)$ as

$$
\begin{aligned}
\mu(\Delta t) &= 2\left(\gamma(0) - \gamma(\Delta t)\right) && (63)\\
&= 2\boldsymbol{w}^T\left(\Sigma_\infty - A_{\Delta t}\Sigma_\infty\right)\boldsymbol{w} && (64)\\
&= \frac{2D\gamma}{k}\boldsymbol{w}^T\left(1 - \mathrm{e}^{-\frac{k}{\gamma}B\Delta t}\right)B^{-1}\boldsymbol{w} && (65)\\
&= \frac{D\gamma}{k}\int_{-\pi}^{\pi}\mathrm{d}\omega\left|\sum_n w(n)e(n,\omega)\right|^2 \frac{1 - \mathrm{e}^{-\frac{2k}{\gamma}\Delta t(1-\cos\omega)}}{1-\cos\omega}\,. && (66)
\end{aligned}
$$

For our case, $y(t)$ should be the distance between two monomers $a$ and $b$, which means $w(n) = \delta_{an} - \delta_{bn}$. We thus find

$$
\sum_n w(n)e(n,\omega) = \frac{1}{\sqrt{2\pi}}\left(\mathrm{e}^{\mathrm{i}\omega a} - \mathrm{e}^{\mathrm{i}\omega b}\right) = \sqrt{\frac{2}{\pi}}\mathrm{i}\mathrm{e}^{\mathrm{i}\omega\frac{a+b}{2}}\sin\omega\frac{a-b}{2}\,, \tag{67}
$$

such that the expression for the MSD becomes

$$
\begin{aligned}
\mu(\Delta t) &= \frac{2D\gamma}{\pi k}\int_{-\pi}^{\pi}\mathrm{d}\omega\,\frac{1-\mathrm{e}^{-\frac{2k}{\gamma}\Delta t(1-\cos\omega)}}{1-\cos\omega}\sin^2\frac{(a-b)\omega}{2} && (68)\\
&\equiv \frac{2D\gamma}{\pi k}\int_{-\frac{\pi}{2}}^{\frac{\pi}{2}}\mathrm{d}z\,\frac{\sin^2 Lz}{\sin^2 z}\left(1 - \mathrm{e}^{-\frac{4k}{\gamma}\Delta t\sin^2 z}\right)\,, && (69)
\end{aligned}
$$

where we utilized $1 - \cos\omega = 2\sin^2\omega$, substituted $z \equiv \frac{\omega}{2}$, and introduced the tether length $L \equiv a - b$.

We now study the three scaling regimes of eq. (69):

- at very short times we expand the exponential to first order and find

$$
\mu(\Delta t \ll \frac{\gamma}{4k}) \approx \frac{2D\gamma}{\pi k}\int_{-\frac{\pi}{2}}^{\frac{\pi}{2}}\mathrm{d}z\,\frac{4k}{\gamma}\Delta t\sin^2 Lz = 4D\Delta t = 2\mu_{\text{single free monomer}}(\Delta t)\,, \tag{70}
$$

  in accordance with the intuition that at short times the monomers do not feel their neighbors and thus diffuse freely.

- at long times, we have that $\mathrm{e}^{-\frac{4k}{\gamma}\Delta t\sin^2 z} \to 0\,\forall z \in \left[-\frac{\pi}{2}, \frac{\pi}{2}\right] \setminus \{0\}$. We thus obtain

$$
\mu(\Delta t \to \infty) = \frac{2D\gamma}{\pi k}\int_{-\frac{\pi}{2}}^{\frac{\pi}{2}}\mathrm{d}z\,\frac{\sin^2 Lz}{\sin^2 z} = 2\frac{D\gamma}{k}L\,, \tag{71}
$$

  where we prove the last equality by induction over $L \in \mathbb{N}_0$: the induction hypothesis $\int_{-\frac{\pi}{2}}^{\frac{\pi}{2}}\mathrm{d}z\,\frac{\sin^2 Lz}{\sin^2 z} = \pi L$ is trivially true for $L = 0$ and $L = 1$; we then use standard trigonometry to show that

$$
\begin{aligned}
\sin^2(L+1)z + \sin^2(L-1)z &= (\sin Lz\cos z + \cos Lz\sin z)^2 + (\sin Lz\cos z - \cos Lz\sin z)^2 && (72)\\
&= 2\sin^2 Lz\left(1 - \sin^2 z\right) + 2\cos^2 Lz\sin^2 z && (73)\\
&= 2\sin^2 Lz + 2\sin^2 z\cos 2Lz\,, && (74)
\end{aligned}
$$

  such that, using the induction hypothesis for $L$ and $L - 1$, we find

$$
\begin{aligned}
\int_{-\frac{\pi}{2}}^{\frac{\pi}{2}}\mathrm{d}z\,\frac{\sin^2(L+1)z}{\sin^2 z} &= 2\int_{-\frac{\pi}{2}}^{\frac{\pi}{2}}\mathrm{d}z\,\frac{\sin^2 Lz}{\sin^2 z} - \int_{-\frac{\pi}{2}}^{\frac{\pi}{2}}\mathrm{d}z\,\frac{\sin^2(L-1)z}{\sin^2 z} + 2\int_{-\frac{\pi}{2}}^{\frac{\pi}{2}}\mathrm{d}z\,\cos 2Lz && (75)\\
&= \pi(2L - (L-1)) && (76)\\
&= \pi(L+1)\,, && (77)
\end{aligned}
$$

  where the integral in the third term runs over full periods of the cosine and thus vanishes. Finally, we make the connection to eq. (33) by restating the result as

$$
\mu(\Delta t \to \infty) = 2\frac{D\gamma}{k}L \equiv 2J\,. \tag{78}
$$

- to find the scaling behavior at intermediate times, where the chain connectivity is relevant but the two tracked loci are still not constrained by their finite tether, we take that tether to be infinitely long, $L \to \infty$. In this limit,

$\sin^2 Lz$ oscillates arbitrarily fast, such that $\int \mathrm{d}z\, f(z) \sin^2 Lz = \frac{1}{2} \int \mathrm{d}z\, f(z)$ for continuous $f(z)$. Consequently, eq. (69) becomes

$$\mu(\Delta t) = \frac{D\gamma}{\pi k} \int_{-\frac{\pi}{2}}^{\frac{\pi}{2}} \mathrm{d}z\, \frac{1 - \mathrm{e}^{-\frac{4k}{\gamma}\Delta t \sin^2 z}}{\sin^2 z} \tag{79}$$

$$= \frac{D\gamma}{\pi k} \int_{-1}^{1} \mathrm{d}\zeta\, \frac{1 - \mathrm{e}^{-\frac{4k}{\gamma}\Delta t \zeta^2}}{\zeta^2 \sqrt{1-\zeta^2}} \tag{80}$$

$$= 4D\Delta t\, \mathrm{e}^{-\frac{2k}{\gamma}\Delta t} \left[ I_0\left(\frac{2k}{\gamma}\Delta t\right) + I_1\left(\frac{2k}{\gamma}\Delta t\right) \right], \tag{81}$$

where we substitute $\zeta := \sin z$ and $I_\alpha(z)$ are the modified Bessel functions of the first kind, which have the asymptotic expansion $I_\alpha(z) = \frac{\mathrm{e}^z}{\sqrt{2\pi z}} \left[1 + \mathcal{O}\left(\frac{1}{z}\right)\right]$. Thus, for $\Delta t \to \infty$ we find

$$\mu(\Delta t) \approx 4D\sqrt{\frac{\gamma \Delta t}{\pi k}} \equiv 2\Gamma\sqrt{\Delta t}\,, \tag{82}$$

with $\Gamma \equiv 2D\sqrt{\frac{\gamma}{\pi k}}$, c.f. eq. (33).

It is most practical to define the crossovers between these regimes by equating the asymptotes, yielding

$$\tau_{D\to\Gamma} = \frac{\gamma}{\pi k} \qquad \text{and} \qquad \tau_{\Gamma\to J} = \frac{\pi}{4k} L^2 \tag{83}$$

for the diffusive-to-Rouse and Rouse-to-equilibrium transitions respectively. Note that $\tau_{\Gamma\to J}$ is a factor $\frac{\pi^3}{4} \approx 7.75$ greater than the commonly quoted Rouse time of $\frac{L^2}{\pi^2 k}$. The latter should not be interpreted as the location of the crossover, but as the time where the MSD starts deviating markedly from the Rouse scaling. Similarly, while $\tau_{D\to\Gamma}$ marks the position of the crossover, numerical evaluation shows that the Rouse scaling remains a good approximation only for $\Delta t \gtrsim \frac{\gamma}{k}$ (**Fig. S8B**).

Finally, comparing eqs. (78) and (82) to eq. (33) we note that the discrete Rouse model provides an excellent approximation to the continuous one, up to two finite size effects: first, for $\Delta t \lesssim \frac{\gamma}{k}$ the MSD starts deviating from the familiar Rouse scaling due to the finite monomer size. We thus take care not to apply the model on time scales shorter than $\frac{\gamma}{k}$. Second, using a finite chain (as we are forced to do in the inference model) speeds up the crossover between the Rouse scaling regime and the steady state. To minimize this effect, we always work with chains at least three times as long as the tether between the tracked monomers (**Fig. S8C**).

### 7 Bayesian Inference of Looping Dynamics (BILD)

#### 7.1 Method

To set up a well-defined inference problem, we parametrize the space of looping profiles $\Theta \equiv \{\theta : [0,T] \mapsto \{0,1\}\}$ as follows:

- $\theta_0 \equiv \theta(0) \in \{0,1\}$ is the initial state,
- $k \in \mathbb{N}_0$ counts the number of switches (0 to 1 or *vice versa*),
- $0 < s_1 < s_2 < \ldots < s_k < T$ are the positions of the $k$ switches.

These parameters fully and uniquely describe any possible looping profile. Clearly the number of parameters needed to describe a given profile depends on the number of switches $k$. It therefore appears natural to regard $k$ as a hyperparameter and define the one-parameter model family

$$M_k \equiv (\Theta_k, \mathcal{L}_k, \pi_k)\,, \tag{84}$$

where $\Theta_k \subset \Theta$ is the subspace of profiles with $k$ switches, $\mathcal{L}_k(\theta, y) = \mathcal{L}(\theta, y) \equiv p(y \mid \theta)$ is the Rouse likelihood from section 6.2, and $\pi_k(\theta) = \text{Uniform}_{\Theta_k}(\theta) = \frac{k!}{2T^k}$ is a uniform prior over profiles with $k$ switches.

We have thereby set up the problem as a hierarchical Bayesian model, which can be inferred by the evidence approximation [34, 39, 40]: first we fix the hyperparameter (number of switches $k$) by maximizing the evidence, then we find the posterior distribution for the actual profile $\theta(t)$ with that fixed number of switches. Directly following this procedure, we infer looping with high recall, but low precision (section 7.3). In order to control this precision-recall trade-off, we introduce the *evidence tolerance* $\Delta E$ as described below. We find that a small $\Delta E > 0$ increases precision, while only marginally decreasing recall (**Fig. S10J**), and set $\Delta E = 2$ throughout the analysis presented in the main text (c.f. section 7.3). A complete overview over our inference scheme follows.

The inference task is to find the best profile $\theta$ for a given observed trajectory $y$. To estimate the hyperparameter $k$ we calculate the log-evidences

$$E_k \equiv \log p(y \mid k) = \log \int_{\Theta_k} \mathrm{d}\theta\, p(y \mid \theta) \pi_k(\theta) \tag{85}$$

and maximize, subject to the tolerance $\Delta E$. Specifically, we find the minimal $\hat{k}$ such that the log-evidence is within $\Delta E$ of the true maximum:

$$k^* := \underset{k}{\operatorname{argmax}}\, E_k = \underset{k}{\operatorname{argmax}}\, p(y \mid \theta) \tag{86}$$

$$\hat{k} := \underset{\{k\,:\, E_{k^*} - E_k \le \Delta E\}}{\operatorname{argmin}} k\,. \tag{87}$$

Having thus identified $M_{\hat{k}}$, we then calculate the posterior distribution over looping profiles under $M_{\hat{k}}$

$$p_{\hat{k}}\left(\theta \mid y\right) = \frac{p\left(y \mid \theta\right) \pi_{\hat{k}}(\theta)}{p(y \mid \hat{k})} \tag{88}$$

and pick the maximum a posterior (MAP) profile as a point estimate of the best profile for the given trajectory $y$:

$$\hat{\theta}(y) = \underset{\theta}{\operatorname{argmax}}\, p_{\hat{k}}\left(\theta \mid y\right)\,. \tag{89}$$

The remainder of this section describes our technical implementation of eqs. (85) and (89) (see also Fig. 3C).

We estimate the integral (85) by Adaptive Multiple Importance Sampling (AMIS; [41]). In the following, $k$ remains fixed and is often suppressed to declutter notation. AMIS utilizes a family of proposal distributions $q_\psi(\theta)$, parametrized by some set of parameters $\psi$, and then alternates between evaluating the target function $f(\theta) \equiv p(y \mid \theta)\pi_k(\theta)$ at points sampled from the proposal, and updating the proposal based on past samples. The key advantage of this approach is that samples from past steps are reweighed appropriately instead of discarded, such that we build up a properly weighted sample relatively quickly, without discarding any of the evaluations (which greatly contributes to computational feasibility). The $n$-th step in this sampling scheme is described in algorithm 1, while we refer to the original work [41] for more details.

**Input:** the sample size $R_n$, proposal parameters $\psi_n$

1 draw $R_n$ new samples $\{\theta_r^n\}_{r=1,\ldots,R_n}$ from the current proposal $q_{\psi_n}(\theta)$

2 let $N_{\text{sample}} \equiv \sum_{i=0}^n R_n$ (the total number of samples)

3 evaluate on *all* samples $\{\theta_r^i\}_{\substack{r=1,\ldots,R_i\\ i=1,\ldots,n}}$ the importance weights

$$\omega_r^i = \frac{N_{\text{sample}} f(\theta_r^i)}{\sum_{j=0}^n R_j q_{\psi_j}(\theta_r^i)} \tag{90}$$

4 estimate the evidence as

$$P(y \mid k) = \int_{\Theta_k} \mathrm{d}\theta\, f(\theta) \approx \frac{1}{N_{\text{sample}}} \sum_{i=0}^{n} \sum_{r=1}^{R_i} \omega_r^i \equiv \bar{\omega} \tag{91}$$

5 calculate the estimates for log-evidence $\hat{E}_k^n$ and its standard error $\Delta\hat{E}_k^n$ as

$$\hat{E}_k^n = \log \bar{\omega}\,, \qquad \Delta\hat{E}_k^n = \frac{\Delta\bar{\omega}}{\bar{\omega}} = \frac{1}{\bar{\omega} N_{\text{sample}}} \sqrt{\sum_{i=0}^{n} \sum_{r=1}^{R_i} \left(\omega_r^i - \bar{\omega}\right)^2} \tag{92}$$

6 find the new proposal parameters $\psi_{n+1}$ by fitting the proposal to the current sample $\left\{\left(\theta_r^i, \omega_r^i\right)\right\}$

**Algorithm 1:** The $n$-th step in AMIS. See [41] for more details. Note that at each iteration, on top of the actual evidence we also get an estimate of the standard error.

To implement this scheme for our problem, we recall that since the number of switches $k$ is fixed, a looping profile is parametrized by its initial value $\theta_0$ and the switch positions $s_1, \ldots, s_k$. We rewrite the latter in terms of the fractions $u_a \equiv \frac{s_{a+1} - s_a}{T} \in [0, 1]$ with $s_{k+1} \equiv T$ (the total trajectory length) and $s_0 \equiv 0$, and write them collectively as $\boldsymbol{u} \equiv (u_0, \ldots, u_k)$. This allows us to employ the proposal distribution

$$q_{m,\boldsymbol{\alpha}}\left(\theta_0, \boldsymbol{u}\right) := \text{Bernoulli}_m\left(\theta_0\right) \otimes \text{Dirichlet}_{\boldsymbol{\alpha}}\left(\boldsymbol{u}\right)\,, \tag{93}$$

where Bernoulli$_m\left(\theta_0\right) \equiv m^{\theta_0}\left(1-m\right)^{1-\theta_0}$ and Dirichlet$_{\boldsymbol{\alpha}}\left(\boldsymbol{u}\right) \equiv \frac{1}{B(\boldsymbol{\alpha})}\prod_{a=0}^{k} u_a^{\alpha_a - 1}$ are, respectively, a Bernoulli distribution with mean $m$ and a Dirichlet distribution with concentration parameters $\boldsymbol{\alpha}$. For initialization of the recursive AMIS scheme we choose the uniform distribution, given by the parameter values $m_0 = \frac{1}{2}$ and $\alpha_a = 1\,\forall a$. In the sampling step, we choose $R_n = 100\,\forall n$. For the updating step we estimate the proposal parameters by the method of moments, setting

$$m_{n+1} = \left\langle \theta_0 \right\rangle_n \,, \tag{94}$$

$$\boldsymbol{\alpha}_{n+1} = \frac{\left\langle \boldsymbol{u} \right\rangle_n}{k}\left(\sum_{a=0}^{k} \frac{\left\langle u_a \right\rangle_n \left(1 - \left\langle u_a \right\rangle_n\right)}{\mathrm{Var}_n\, u_a} - k\right) \,, \tag{95}$$

where $\left\langle \cdot \right\rangle_n$ are expectation values under the sample at step $n$, and similarly

$$\mathrm{Var}_n\, u_a \equiv \frac{1}{\sum_{i,r}\omega_r^i} \sum_{i=0}^{n}\sum_{r=1}^{R_i} \omega_r^i \left(\left(u_a\right)_r^i - \left\langle u_a \right\rangle_n\right)^2 \tag{96}$$

is the variance of $u_a$ at step $n$. To prevent overfitting of the proposal distribution to small samples at the beginning of the iterative scheme (the first proposal update happens after $R_0 = 100$ samples are drawn), we introduce two *braking parameters* $b_m$ and $b_\alpha$. Whenever parameters are updated, these limit the difference to the previous value:

$$\left|m_{n+1} - m_n\right| \le R_n b_m \,, \tag{97}$$

$$\left|\log \frac{\left|\boldsymbol{\alpha}_{n+1}\right|_1}{\left|\boldsymbol{\alpha}_n\right|_1}\right| \le R_n b_\alpha \,, \tag{98}$$

$$\tag{99}$$

where $\left|\alpha\right|_1 \equiv \sum_a \alpha_a$ is the total concentration. In practice we use $b_m = 10^{-3}$ and $b_\alpha = 10^{-2}$. With this, we now have all the ingredients to initialize AMIS, run individual sampling steps using algorithm 1, and perform well-regulated updates of the proposal parameters.

Finally, we have to provide a stopping criterion for the iterative AMIS scheme. To that end, we take a step back and consider the problem of $k$ selection as a whole. For $k = 0$ there are only two possible profiles (completely looped or completely unlooped), so we can calculate the evidence exactly with two likelihood evaluations. For $k = 1$, taking into account our constraining switch positions to integer frames (section 6.2), we can still completely enumerate all possible profiles with $2T \sim 10^2$ evaluations of the likelihood. For $k = 2$ we would have to enumerate $2T(T-1) \sim 10^4$ profiles, while the sampling approach usually converges with $\sim 10^3$ evaluations, so for $k \ge 2$ we resort to sampling. However, taking this iterative approach, when sampling at $k$ we can assume to already have an estimate or exact value for $E_{\tilde{k}}$ for all $\tilde{k} < k$. Importantly, the sampling also gives us estimates of the standard error $\Delta\hat{E}_k$, so after each AMIS iteration we can check which of the estimates still has a realistic chance of being the best one. We formalize this approach in algorithm 2, thus ultimately obtaining all relevant estimates $\hat{E}_k$.

**Input:** $k_{\text{max}} = 10$, $n_{\text{init}} = 20$, $C_{\text{close}} = 5$, $\Delta k_{\text{lookahead}} = 2$
**Output:** posterior samples and evidence estimates $\hat{E}_k$

calculate $E_0$ and $E_1$ by complete enumeration
**for** $k \le k_{\text{max}}$ **do**
  run $n_{\text{init}}$ AMIS samples to get initial estimate of $E_k$
  **repeat** // relevance resolution
    find $k^* \equiv \operatorname{argmax}_k \hat{E}_k$
    find all $k$ that are reasonably close to this maximum:

$$I \equiv \left\{ k : \frac{\hat{E}_{k^*} - \hat{E}_k}{\sqrt{\left(\Delta\hat{E}_k\right)^2 + \left(\Delta\hat{E}_{k^*}\right)^2}} \le C_{\text{close}} \right\} \tag{100}$$

    find the $k_\Delta \in I$ where the estimated error on the evidence is highest
    run one more AMIS iteration for $k = k_\Delta$
  **until** $|I| = 1$ **or** $k + 1 - \Delta k_{\text{lookahead}} \in I$
**end**
run through the relevance resolution once more

**Algorithm 2:** Scheme for successive AMIS sampling, focussed on sampling the relevant $k$ values.

Having estimated the evidences $E_k$ and found the optimal number of switches $k$ via eq. (87), we now turn to the question of finding the optimal profile $\theta$ with that given number of switches. To that end we note that in calculating the evidences via AMIS, we already generated extensive posterior samples for all relevant $k$. We can therefore simply pick the profile $\theta_r^i$ with the highest posterior weight $\omega_r^i$ as point estimate. We note that on top of this point estimate we actually obtain a full (weighted) posterior sample

$$S \equiv \left\{ \left( \theta_r^i, \omega_r^i \right) \right\} , \tag{101}$$

which will be used in some of the downstream analysis in section 7.4.

Summarizing, we have shown in this section how we infer the looping profile $\theta$ from an observed trajectory, using the Rouse likelihood (59), hierarchical Bayesian inference, and Adaptive Multiple Importance Sampling (AMIS). We refer to this approach as Bayesian Inference of Looping Dynamics (BILD).

#### 7.2 Calibration of the inference model

To run the inference scheme described in section 7.1, we have to make sure that the underlying model (section 6.2) accurately captures the behavior we expect for the looped and unlooped states, i.e. we have to find numerical values for the constants such that the model captures our data most accurately (**Fig. S9B**).

In a first step, we calibrate the model for the unlooped state. Following (Fig. 3A), the unlooped state should capture the behavior of the chain in the absence of sustained looping. It is thus a coarse-grained representation, capturing not only the fully unlooped conformations, but also partial extrusion and random transient contacts ("everything except the looped state"). Our experimental realization of this situation is the CTCF-AID cell line, in which the dynamics of the *Fbn2* loci should be mostly the same as in C36 (the "wild-type" cell line), except for the absence of sustained looping. We find that we can capture the dynamics of this condition well with an MSD of the form (32), our expectation for two loci on a linear polymer. We stress that this does not amount to the assumption that chromatin in the absence of CTCF is unlooped, but we are simply subsuming all the conformations associated with the unlooped state into an "effectively free" chain. One important consequence is that from this calibration of the unlooped state we cannot yet assemble the looped state (see below). To capture the physical parameters of this coarse-grained model, we use the method of section 5 to fit the MSD (32) to the CTCF-AID data, thus obtaining numerical values for the phenomenological parameters $\Gamma$ and $J$. We now utilize the correspondence between the continuous and discrete Rouse model (section 6.3) to assemble the discrete model needed for BILD from the continuous model we use to capture the dynamics of the calibration data.

The model for the unlooped state is specified by eqs. (34) and (46), from which we identify the following parameters: diffusivity $D$ and friction constant $\gamma$ of individual monomers; the spring constant $k$ determining the strength of the backbone bonds; the number $N$ of monomers, and positions $a$ and $b$ of the tracked loci on the chain (**Fig. S9A**). First of all, we note that eq. (34) can always be rescaled by $\frac{1}{\gamma}$, such that our effective degrees of freedom are $D$, $\frac{k}{\gamma} \in \mathbb{R}^+$, $N \in \mathbb{N}$, and $a, b \in \{1, \dots, N\}$ (wlog $a > b$). We also use the auxiliary variable $L \equiv a - b$. From the fits to the experimental data we know

$$\Gamma = 2D\sqrt{\frac{\gamma}{\pi k}} \qquad \text{and} \qquad J = \frac{D\gamma}{k} L \,. \tag{102}$$

Together with the constraint mentioned at the end of section 6.3 this reads

$$\frac{J}{\Gamma} = \sqrt{\frac{\pi\gamma}{4k}} L \,, \qquad \frac{\Gamma^2}{J} = \frac{4D}{\pi L} \,, \qquad \text{and} \qquad \Delta t_{\text{frame}} > \frac{\gamma}{k} \,. \tag{103}$$

Using only the first two equations to fix the microscopic parameters, we would have one degree of freedom left, since we can always add more monomers to the chain (increase $L$), if we rescale $D$ and $k$ appropriately. For computational efficiency we prefer $L$ to be as small as possible, which in this context means that we aim to satisfy the bound provided by the last inequality as tightly as possible (with integer $L$). We therefore find

$$L = \left\lceil \sqrt{\frac{4}{\pi}} \frac{J}{\Gamma\sqrt{\Delta t_{\text{frame}}}} \right\rceil \,, \qquad D = \frac{\pi L \Gamma^2}{4J} \,, \qquad \frac{k}{\gamma} = \frac{\pi}{4} \left( \frac{L\Gamma}{J} \right)^2 \,, \tag{104}$$

where $\lceil x \rceil$ indicates the smallest integer larger than $x$. The remaining parameters for the unlooped state are now the total length of the chain $N$ and the positions $a$, $b \equiv a - L$ of the tracked monomers. We choose

$$N = 3L + 1 \,, \qquad a = 2L + 1 \,, \qquad b = L + 1 \,, \tag{105}$$

such that on both ends of the "chain of interest" of length $L$ we have chains of equal length $L$, resulting in a total chain long enough to emulate an infinite polymer up to the relaxation time scale of the chain of interest. This concludes our calibration of the unlooped state, which finally is determined by eqs. (104) and (105), in terms of $\Gamma$ and $J$ of eq. (102), which we find by fitting the CTCF-AID data (**Fig. S9B**).

For the looped state of our inference model we now have only the parameters associated with the extra bond left to determine. These are its strength $r$ relative to a backbone bond (such that its spring constant is $rk$) and the two monomers $m$ and $n$ it connects to (**Fig. S9A**). Assuming the chain in the loop to be much longer than the single extra bond (thus not contributing to the steady state variance in the looped state), the effective tether length in the looped state is given by $L_\text{looped} = a - m + \frac{1}{r} + n - b$ (**Fig. S9A**), such that we find the steady state variance as

$$J_\text{looped} = \frac{D\gamma}{k} L_\text{looped} \,. \tag{106}$$

Experimentally, we can access $J_\text{looped}$ via our RAD21-AID cell line and genomic information. For RAD21-AID (and only there) we can assume that the chain length $L_{\triangle\text{RAD21}}$, which we can extract from fitting the steady state variance $J_{\triangle\text{RAD21}}$, matches the known genomic separation of our tracers of 515 kb. Under the Rouse model (c.f. e.g. eq. (33)) $J \propto L$, such that we get

$$J_\text{looped} = \frac{10\,\text{kb}}{515\,\text{kb}} J_{\triangle\text{RAD21}} \,, \tag{107}$$

where 10 kb is the combined distance of the tracked loci from the CTCF sites, i.e. the effective tether length in the looped state. This allows us to calculate $L_\text{looped}$ from eq. (106) and the previously determined constants $D$ and $\frac{k}{\gamma}$, as well as the ΔRAD21 steady state variance $J_{\triangle\text{RAD21}}$. We then convert the real-valued $L_\text{looped}$ into the parameters of the extra bond as outlined in algorithm 3, which is designed to achieve two goals: first, place the extra bond as symmetrically as possible between the tracked monomers. Second, keep $r$ and $\frac{1}{r}$ as close to $1$ as possible, such that the extra bond is as similar to a backbone bond as possible. We realize this latter goal by choosing $r \in \left[\frac{1}{\varphi}, \varphi\right]$, with the golden ratio $\varphi = 1 + \frac{1}{\varphi} = \frac{1+\sqrt{5}}{2}$.

**Input:** $L_\text{looped} \in \mathbb{R}^+$, positions $a, b$ of the tracked monomers
**Output:** parameters $m$, $n$, $r$ of the extra bond

**if** $L_\text{looped} \geq \frac{1}{\varphi}$ **then**
  find $\bar{L} \in \mathbb{N}_0$, $r \in \left[\frac{1}{\varphi}, \varphi\right]$ such that $L = \bar{L} + \frac{1}{r}$
**else**
  $\bar{L} \leftarrow 0$
  $r \leftarrow L_\text{looped}^{-1}$
**end**
$m \leftarrow a - \left\lceil \frac{\bar{L}}{2} \right\rceil$
$n \leftarrow b + \left\lfloor \frac{\bar{L}}{2} \right\rfloor$

**Algorithm 3:** Splitting the real-valued $L_\text{looped}$ into proper parameters for the extra bond.

In summary, to calibrate our inference model to experimental data we

- fit the unlooped state to our CTCF-AID data, then
- match strength and position of the extra bond to the RAD21-AID steady state, rescaled by the known genomic separations.

This allows us to fully calibrate the model (**Fig. S9B**), and thus run BILD (section 7.1).

##### 7.3 Benchmarking BILD on simulations

We benchmarked BILD by comparing its output on a data set of 53 simulations out of the sweep performed in section 4.4 to the corresponding ground truth, i.e. the intervals where the (simulated) CTCF sites were truly bridged by (simulated) cohesin (**Fig. S10A**). We begin by investigating the performance of the "raw inference", setting the evidence bias parameter $\Delta E$ (introduced in eq. (87)) to zero. To combat the high false positive rate uncovered through this analysis, we then proceed to study $\Delta E > 0$ (see also section 8). Where appropriate below, we use standard terminology for binary classification tasks, including *true positive* (TP), *false positive* (FP; type I error), *false negative* (FN; type II error), the performance scores

$$\text{Recall} \equiv \frac{\text{TP}}{\text{TP} + \text{FN}} \qquad \text{and} \qquad \text{Precision} \equiv \frac{\text{TP}}{\text{TP} + \text{FP}} \,, \tag{108}$$

and the *prevalence*, which is defined as the fraction of ground truth positive data in the whole data set. In the context of this study, we also refer to the prevalence as the *looped fraction*, the fraction of time the locus spends in the looped state. We now study the inference performance from two perspectives

- First, the point-wise view: at each time point in each trajectory, the inference is a binary classification task, calling "looped" or "unlooped" states. Overall we find a recall of 96%, at 54% precision (c.f. **Fig. S9J**), meaning the inference reliably identifies looping, but roughly half of the data that is inferred as looped are false positives. We thus expect to overestimate the prevalence (looped fraction) by a factor Recall/Precision $\approx 1.8$. In fact, however, we find (**Fig. S9G**)

$$f_{\text{looped}}^{\text{inferred}} \approx 1.34 f_{\text{looped}}^{\text{true}} + 2.8\%\,, \tag{109}$$

  leading us to assume two distinct error mechanisms: any true looping in the trajectory will be overestimated by a factor $\sim 1.34$, and on top of that $\sim 2.8\%$ of the trajectory will be called "looped" just due to random fluctuations. Importantly, while the former is stable under our evidence bias parameter $\Delta E$, the second effect shows some sensitivity to $\Delta E$ (**Fig. S9G,J**), which we can therefore use to control this error contribution. In summary the point-wise perspective shows that most looping is indeed detected, but there is a significant number of false positives. We can quantify the overall effect on the inferred looped fraction relatively well and thus correct for this overestimate.

- Second, the interval-wise view (based on [42]): since ultimately we are most interested in the lifetime of the looped state, we have to ensure that whole looping intervals are inferred correctly. The biggest concern here is that of cardinality: a single true interval might be split in the inference, leading to (say) three different inferred intervals that, even though they might collectively cover the true interval completely and accurately, by themselves would have a lifetime approximately three times shorter than the true interval. Conversely, a single inferred interval might cover multiple true intervals and thus have a much longer inferred lifetime than the individual true intervals. We term the number of true intervals overlapping with each inferred interval the *type I cardinality*; similarly, the *type II cardinality* labels the number of inferred intervals overlapping with each true interval. We find both cardinalities to be 0 or 1 for almost all intervals, with cardinality $> 1$ occuring in only 0.79% and 1.6% of the cases for type I and type II respectively (**Fig. S9B,C**). We therefore conclude that cardinality is not a significant problem in this study.

  Having established that most inferred intervals correspond to at most one true interval, we can immediately study how well we capture the lifetime of these true intervals; we generally find that if the inferred interval corresponds to a true one, it also captures the lifetime well (**Fig. S9D**). We therefore arrive again at the picture of a binary classifier, but now at the interval level: generally, an inferred interval will either accurately capture an interval in the true profile, or it will be a false positive altogether. This binary picture is reinforced by **Fig. S9E** showing that each true interval is either inferred correctly, or missed completely.

  On the interval level, we find a high false positive rate of 61%. These false positive inferences are composed of two populations (**Fig. S9F**): first, we find intervals that correspond to "almost looped" events, where the effective tether length between the CTCF sites becomes very short, but does not quite cross the threshold of $1.1$ monomers that we use to define the looped state. This population is robust under variation of $\Delta E$. Second, we find a population of short ($\lesssim 10$ frames) false positive detections that are independent of the extrusion state of the locus. These are random fluctuations of the polymer/noise that happen to resemble the looped state and show a strong dependence on $\Delta E$, which we can thus use to control these erroneous inferences.

Having seen that we can use our evidence bias $\Delta E$ to control the false positive inferences, we now study how it affects the lifetime distribution we infer from our data (**Fig. S9H**). As expected, high $\Delta E$ mostly removes short intervals from the inference, thus reducing the false positive fraction (increasing precision; **Fig. S9J**). At the same time though, recall is reduced as some truly short intervals are not found anymore. It is *a priori* not clear how best to balance these effects, so ultimately $\Delta E$ remains a free parameter. From the precision-recall curves in **Fig. S9J** we conclude that $0 < \Delta E < 5$ improves overall inference quality (higher increase in precision than decrease in recall). A reasonable value within that range seems to be $\Delta E \approx 2$, for which the Kolmogorov-Smirnov distance between the true and inferred lifetime distributions on our validation data set becomes minimal (**Fig. S9K**). We emphasize, however, that this is essentially a rule of thumb, and any final results of the inference should be checked for their variation with $\Delta E$. Corresponding analysis of our results from the main text (Fig. 3F,G) can be found in **Fig. S11.**

##### 7.4 Downstream processing: estimation of looped fraction and loop lifetime

In this section we describe how we use the results of the inference (section 7.1) to measure looped fraction and loop lifetimes, the latter either non-parametrically with the Kaplan-Meier estimator, or with an exponential fit.

We define the looped fraction $f_{\text{looped}}$ as

$$\text{looped fraction} = \frac{\text{time spent in looped state}}{\text{total trajectory length}}\,, \tag{110}$$

which is straight-forward to calculate from the inferred looping profiles $\theta(t)$. We bootstrap an ensemble of mean looped fractions as described in algorithm 4, allowing us to report the mean over that ensemble as a point estimate, and its $2.5$th and $97.5$th percentile as $95\%$ confidence interval.

**Input:** for each trajectory $y_n(t)$ the posterior sample $S_n$ from eq. (101), $N_{\text{bootstrap}} = 1000$
**Output:** a list $F$ of $N_{\text{bootstrap}}$ evaluations of the mean looped fraction $\langle f_{\text{looped}} \rangle$

initialize an empty list $F$
**for** $i \in \{1, \ldots, N_{\text{bootstrap}}\}$ **do**
    initialize an empty list $A$
    **for** *each trajectory* $n$ **do**
        sample a profile $\theta_r^i$ from $S_n$ (i.e. according to the weights $\omega_r^i$)
        calculate the fraction looped $f_{\text{looped}}(\theta_r^i)$ for that profile
        append $f_{\text{looped}}(\theta_r^i)$ to $A$
    **end**
    append the mean over $A$ to $F$
**end**
**return** F

**Algorithm 4:** Bootstrapping the mean looped fraction

In calculating a characteristic lifetime of the looped state, we encounter the problem of censoring (c.f. **Fig. S10A**). This means that many of the inferred looping intervals start or end with the trajectory, such that we do not observe them fully. Simply calculating mean lifetimes from the inferred intervals in this scenario would provide a heavy underestimate of the true lifetime. This censoring problem is well-known in the medical literature and customarily addressed by using the Kaplan-Meier estimator for the survival function [43]. We follow that standard procedure: from the MAP looping profiles provided by BILD (section 7.1) we compute the set $\{(t_i, c_i)\}$ of loop lifetimes $t_i$ and corresponding boolean variables $c_i$ indicating whether the observation $t_i$ was censored or not. We then use the Kaplan-Meier estimator

$$\hat{S}(\tau) = \prod_{t_i < \tau} \left(1 - \frac{d_i}{N_i}\right) \tag{111}$$

for the survival function $S(\tau) := P(t \geq \tau)$. Here, $d_i$ is the number of uncensored events of length $t_i$, while $N_i$ is the total number of events of length greater than $t_i$, censored or uncensored. Confidence intervals (at confidence level $1 - \alpha$) for this estimator are given by the exponential Greenwood formula

$$\mathrm{e}^{z_{\alpha/2}\sqrt{V(t)}} \log \hat{S}(t) \quad < \quad \log S(t) \quad < \quad \mathrm{e}^{-z_{\alpha/2}\sqrt{V(t)}} \log \hat{S}(t)\,, \tag{112}$$

$$V(t) \equiv \left(\log \hat{S}(t)\right)^{-2} \sum_{t_i < t} \frac{d_i}{N_i\,(N_i - d_i)}\,, \tag{113}$$

with $z_{\alpha/2}$ the $\frac{\alpha}{2}$th quantile of the normal distribution [44, 45]. Note that generally $z_{\alpha/2} < 0$ and specifically for 95% confidence intervals we have $z_{0.025} = -1.96$.

Finally, we also provide an estimate of the median lifetime from an exponential fit to the lifetime distribution. Starting from the same set $\{(t_i, c_i)\}$ as for the Kaplan-Meier estimate, we aim to fit the distribution

$$p(t) = \frac{1}{\tau}\mathrm{e}^{-\frac{t}{\tau}}\,. \tag{114}$$

The likelihood contributions for uncensored ($c_i = 0$) and censored ($c_i = 1$) intervals are respectively

$$p(t_i \mid \tau, c_i = 0) = \frac{1}{\tau}\mathrm{e}^{-\frac{t_i}{\tau}}\,, \qquad p(t_i \mid \tau, c_i = 1) = \int_{t_i}^{\infty} \mathrm{d}t\, p(t) = \mathrm{e}^{-\frac{t_i}{\tau}}\,. \tag{115}$$

The total log-likelihood of observing the given data from a model with mean lifetime $\tau$ is then the sum of these contributions:

$$\log p(\boldsymbol{t} \mid \tau, \boldsymbol{c}) = -N_{\text{uncensored}} \log \tau - \frac{1}{\tau}\sum_i t_i\,, \tag{116}$$

where we introduce the number of *uncensored* observations $N_{\text{uncensored}}$. Taking the derivative and finding its root gives the MLE point estimate

$$\hat{\tau} = \frac{1}{N_{\text{uncensored}}} \sum_i t_i\,. \tag{117}$$

To find confidence intervals (at confidence level $1 - \alpha$) on this estimate, we numerically find the roots $\tau_\pm$ of

$$\log p(\boldsymbol{t} \mid \tau_\pm, \boldsymbol{c}) \overset{!}{=} \log p(\boldsymbol{t} \mid \hat{\tau}, \boldsymbol{c}) - \frac{1}{2}\chi^2_{1,1-\alpha}\,, \tag{118}$$

where $\chi^2_{n,p}$ is the $p$th percentile of the $\chi^2$ distribution with $n$ degrees of freedom. Specifically, $\chi^2_{1,0.95} \approx 3.84$.

Note that $\hat{\tau}$ is an estimate for the mean of the distribution, but from our non-parametric approach via the Kaplan-Meier curves we can estimate only medians. We thus finally also report not the mean $\hat{\tau}$ of the fitted exponential, but its median $\hat{\tau} \log 2 \approx 0.7\hat{\tau}$.

#### 8 Variation in inference results with evidence bias

As shown in section 7.3, the final results of our looping inference depend on the free parameter $\Delta E$. We found from simulations that $\Delta E > 0$ helps to combat false positives, but what exactly this parameter should be remained unclear. Based on those simulations, a reasonable range is $0 < \Delta E < 5$, with $\Delta E \approx 2$ performing well overall. We proceed in a manner similar to section 7.3, first taking a point-wise perspective and studying the looped fraction, then moving on to an interval-based point of view and loop lifetimes.

In **Fig. S10G** we saw that the relationship between true looped fraction $f_{\text{looped}}^{\text{true}}$ and inferred looped fraction $f_{\text{looped}}^{\text{inferred}}$ is captured well by a linear relationship, whose prefactor (relative overestimation of looped fraction) is nearly independent of $\Delta E$, while the offset (random inference of looping without correspondence to ground truth) decreases with $\Delta E$. A fuller picture of this relationship is shown in **Fig. S11A**, left panel, where we perform that linear regression for $\Delta E \in [0, 10]$. This relationship being comparatively robust allows us to use it for correction of looped fractions at all $\Delta E$. On our simulated data set, this leads to a corrected inferred looped fraction $f_{\text{looped}}^{\text{corrected}}$ that is now nearly independent of $\Delta E$ and deviates from the ground truth looped fraction by a few percentage points at most (**Fig. S11A**, middle). This corrected looped fraction now shows a strong correlation with the ground truth value (**Fig. S11A**, right). We then applied this same correction to our real data. We find that we do not abolish all variation with $\Delta E$, hinting that the real data contains errors not accounted for in the simulations. Nevertheless, we find a corrected looped fraction for our C36 cell line of $2 - 4\%$, with the WAPL degron generally exhibiting an increased looped fraction of three percentage points above C36. The correction also places the looped fractions for the CTCF and RAD21 degrons close to zero, which shows that indeed in these degron conditions there is very little looping, if any at all (**Fig. S11B**).

We then investigated the lifetime inference outlined in section 7.4. Specifically, we asked how $\Delta E$ affects the median lifetime as estimated from the Kaplan-Meier survival curves, which is our main estimate of loop life time. For each simulation in our validation data set we find the value of $\Delta E$ that makes the inferred median lifetime match the ground truth median, measured from the Kaplan-Meier survival curve of the true intervals in the data set. In **Fig. S11C** we then show how the inferred lifetime deviates from the true one if $\Delta E$ deviates from its optimal value. We find that while too small $\Delta E$ can lead to heavy underestimation of the lifetime, with a $\Delta E$ somewhat higher than optimal we estimate a lifetime that is at most a factor 2 higher than the true one. We therefore aim to err on the side of high $\Delta E$.

Plotting the inferred lifetime of the real data against $\Delta E$ reveals the change to be modest over the region $1 \leq \Delta E \leq 3$ (**Fig. S11D**), matching our prior expectation of $\Delta E \approx 2$ (**Fig. S10K**). Within that range we find a median loop lifetime in C36 of 10-20 min. To include our exponential estimate (**Fig. 3G**), we ultimately report a range of 10-30 min in the main text.

#### 9 Software, code, and data availability

The code and software used in this paper can be found in **Table S5**. The Micro-C and ChIP-Seq data associated with this paper are available at https://www.ncbi.nlm.nih.gov/geo/ under accession number GSE187487. For a list of datasets used in this paper, please see **Table S6**. The raw trajectory data is available at: https://doi.org/10.5281/zenodo.5770531

### 10 Supplementary Figures

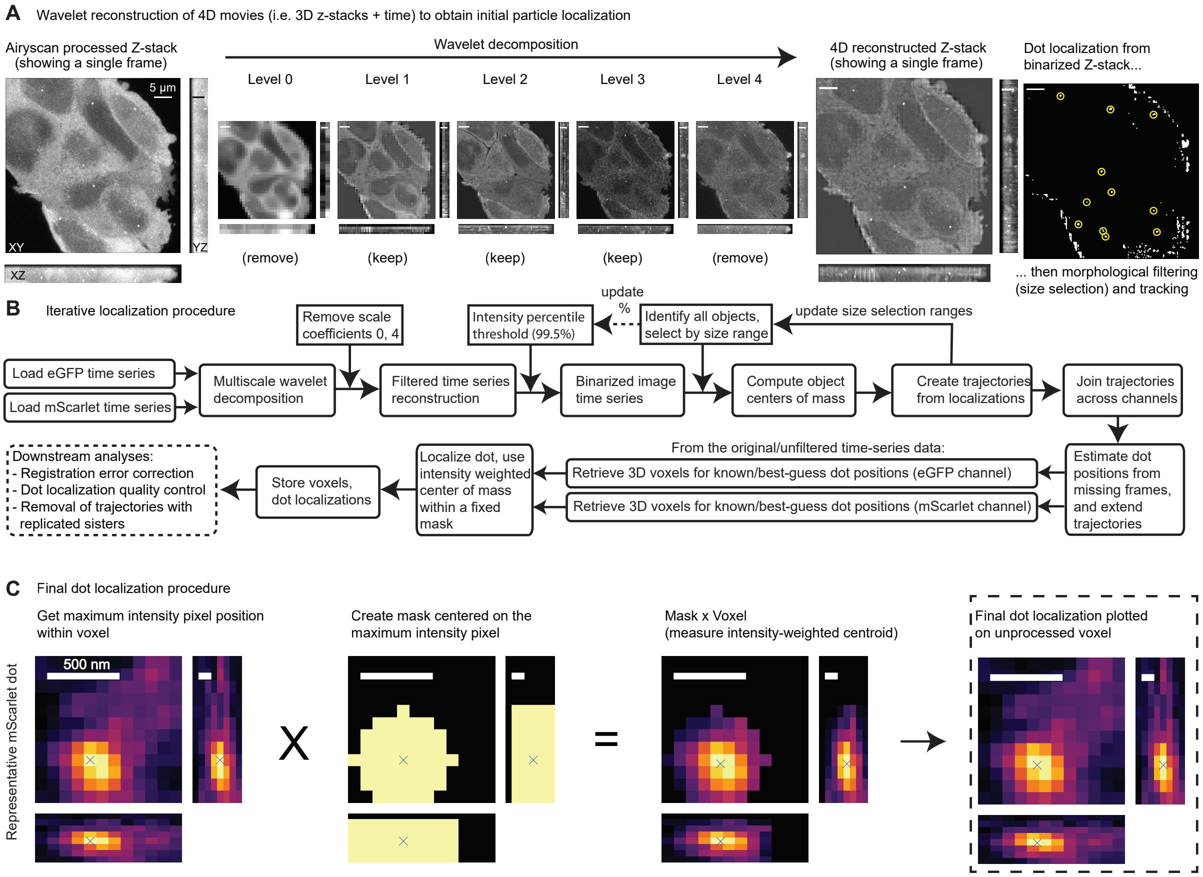

**Fig. S1: Image processing pipeline for dot localization.** **(A)** Overview of the wavelet decomposition method used to filter out background cellular fluorescence and high frequency noise from the 3D image time series. Reconstructed z-stacks are subsequently used with the iterative thresholding and morphological filtering method to identify dots. The image shown is representative of the signal in the EGFP channel after 200 timepoints of imaging and photobleaching (≈67 min corresponding to 200 times 30 z-stacks of images)). **(B)** Workflow diagram showing the iterative thresholding procedure used to identify and track dots in the 3D image time series. **(C)** Illustration of the centroid fitting routing used to localize dots. Localization is performed on AiryScan processed (but otherwise unfiltered) data. Pixel intensities within a fixed spheroidal mask centered on the maximum intensity pixel are used to compute the intensity-weighted center of mass of the dot, achieving average localization precision per dot as tabulated in **Table S3**.

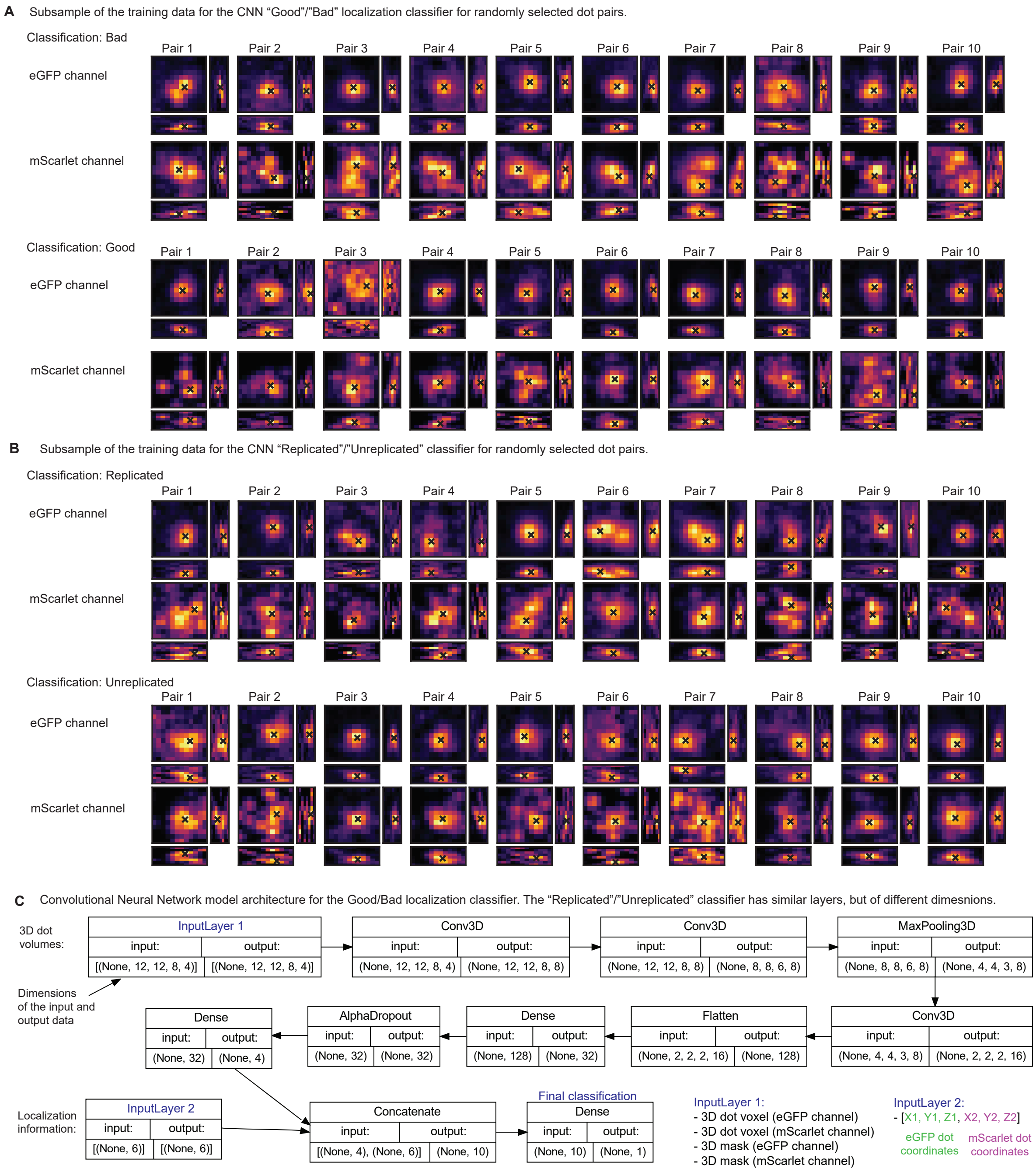

**Fig. S2: Convolutional Neural Network training data and model architecture for classifying dots. (A)** Example maximum intensity projections of voxels used for training the “Good/Bad” classifier, or **(B)** the “Replicated/Unreplicated” classifier. **(C)** The CNN model architecture showing the sequence of layers, inputs, outputs and the sizes of the data for each dimension. Input Layer 1 contains the 3D voxels of each dot as seen by the MIPs in as well as the masked dot localizations (see **Fig. S1B**). Input Layer 2 contains the dot coordinates relative to the voxel position. The final classification is a floating point number ranging from 0 to 1 which classifies the dot pair (e.g. “Good” or “Bad”) based on its value above or below 0.5.

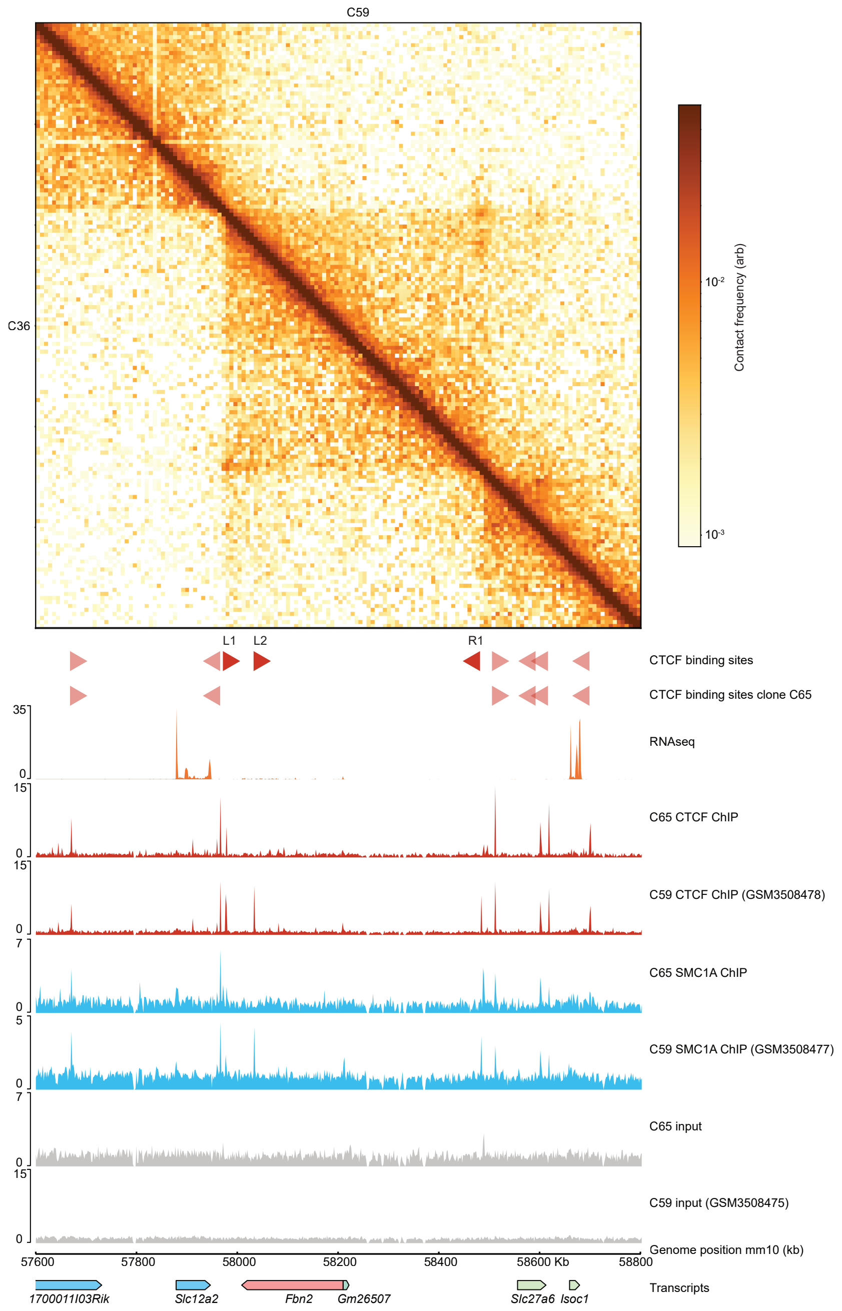

**Fig. S3: Deletion of L1, L2, R1 CTCF binding sites largely abolish CTCF and SMC1A recruitment in clone C65.** Contact map showing Micro-C data from C36 (bottom left) and C59 (top right), at the *Fbn2* locus, bin size = 4kb. Colorbar represent log normalized counts. Genome coordinates mm10. Red triangles show all CTCF binding sites together with the orientation or polarity of the CTCF motif. At the bottom: ChIP-seq tracks of C59 (wild type condition) and C65 ($\Delta$L1, $\Delta$L2, $\Delta$R1 CTCF binding sites), transcripts (GRCm38).

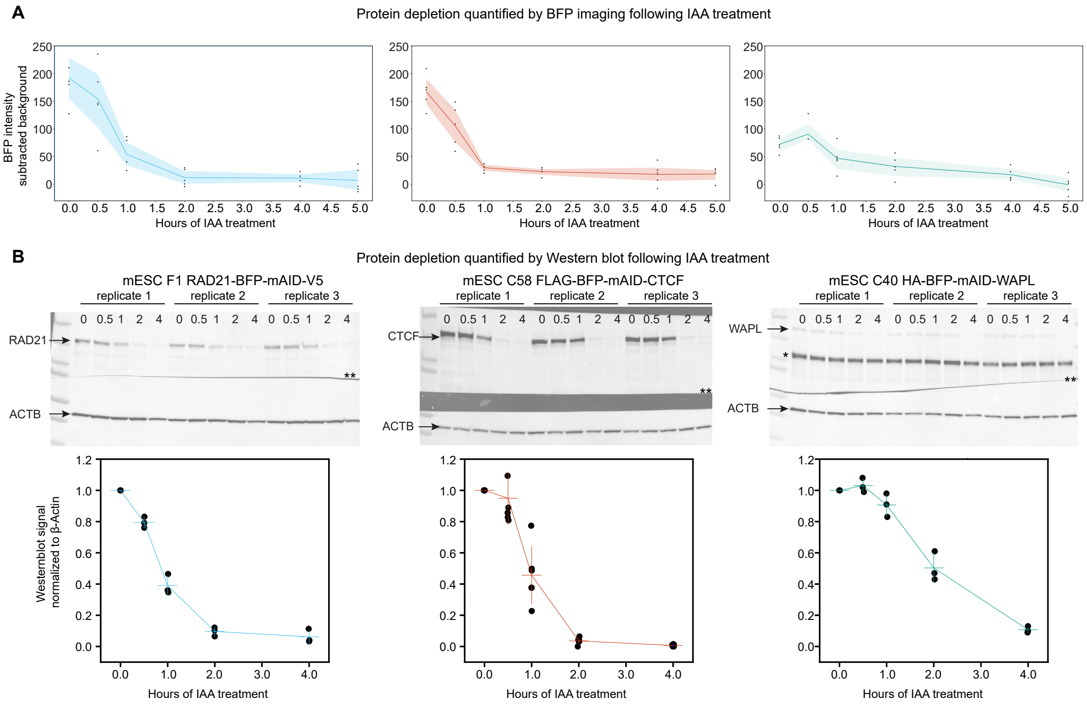

**Fig. S4: Time-course of auxin-induced protein depletion of RAD21, CTCF, and WAPL. (A)** Quantification of tagged protein depletion after IAA administration (500 µM) by BFP imaging. Each timepoint (n = 5) represent the average signal across all cells in one field of view. Fluorescence background was substracted to estimate the nuclear BFP signal. **(B)** Quantification of tagged protein depletion after IAA administration by western blot analysis. Each depletion experiment was performed in three replicates for RAD21 and WAPL, and six replicated for for CTCF. *: non-specific band. **: where the membrane was cut horizontally. Below, imageJ quantification of protein depletion as measured by Western blotting by normalizing the protein of interest to ACTB.

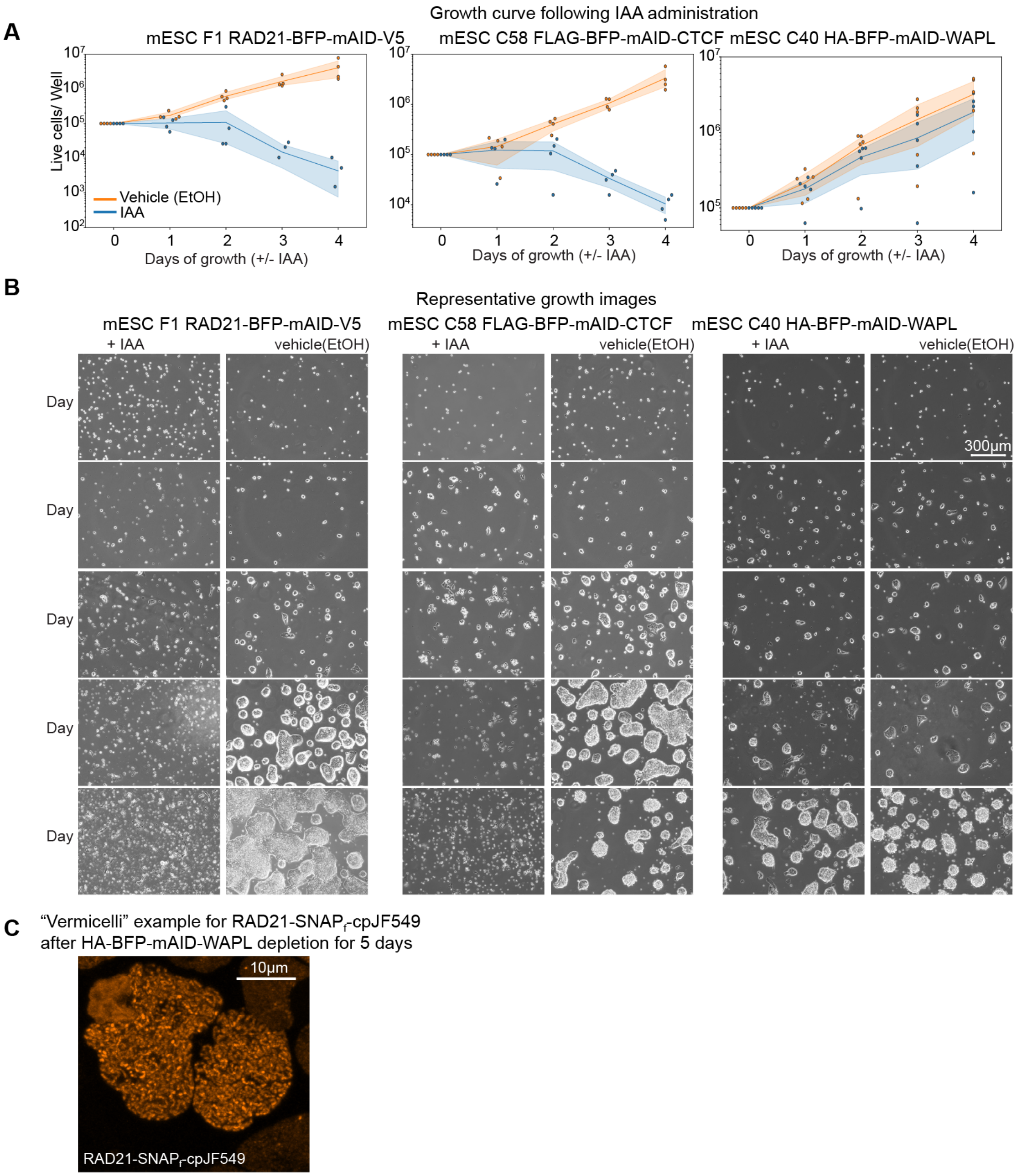

**Fig. S5: Long term effect on cell growth of protein depletion of RAD21, CTCF and WAPL. (A)** Growth curve following IAA administration (500 μM). Each timepoint shows the counted number of live cells. Each experiment was repeated 4 times (RAD21, CTCF) or 5 times (WAPL). **(B)** Representative images in each time point with and without IAA treatment. Phase contrast images were acquired with an EVOS5000 microscope using a 10X objective. **(C)** Example of "Vermicelli chromosomes" resulting from long-term of WAPL depletion (5 days of treatment). Confocal live-cell imaging of RAD21-SNAP$_f$-V5 using cp-SNAP-JF549.

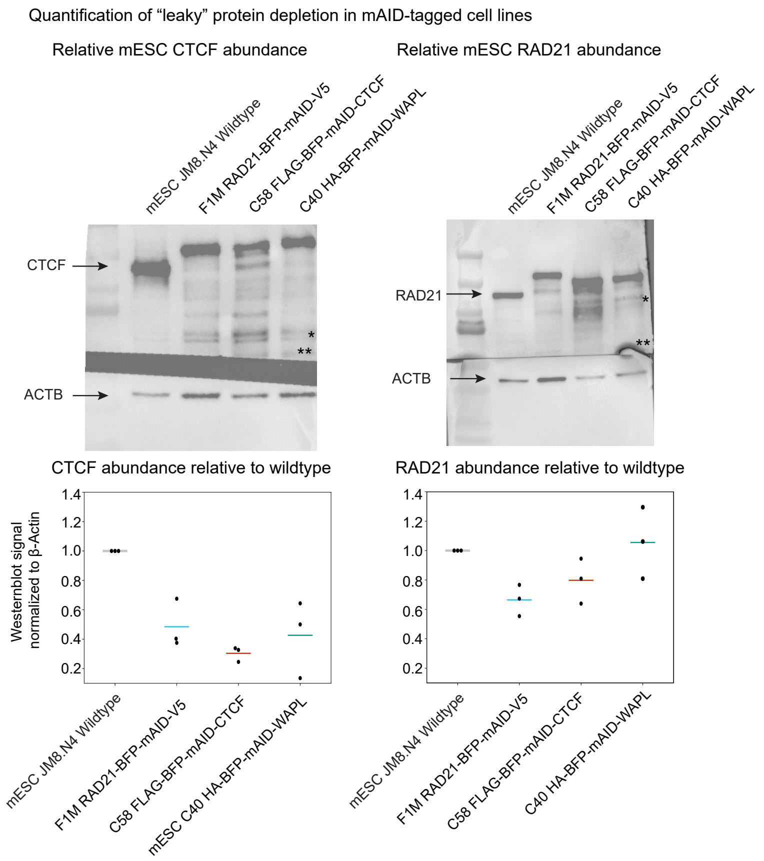

**Fig. S6: Quantification of ’leaky’ protein depletion in AID-tagged RAD21, CTCF, and WAPL cell lines** Quantification of relative mESC expression of CTCF and RAD21 relative to JM8.N4 wildtype mESCs by western blot analysis. Each Western blot experiment was performed in three replicates for JM8.N4 wildtype mESCs, clone F1M RAD21-AID, clone C40 AID-WAPL, and clone C58 AID-CTCF. *: non-specific band. **: membrane cut horizontally. Below, imageJ quantification of protein abundance as measured by Western blot after normalizing to ACTB.

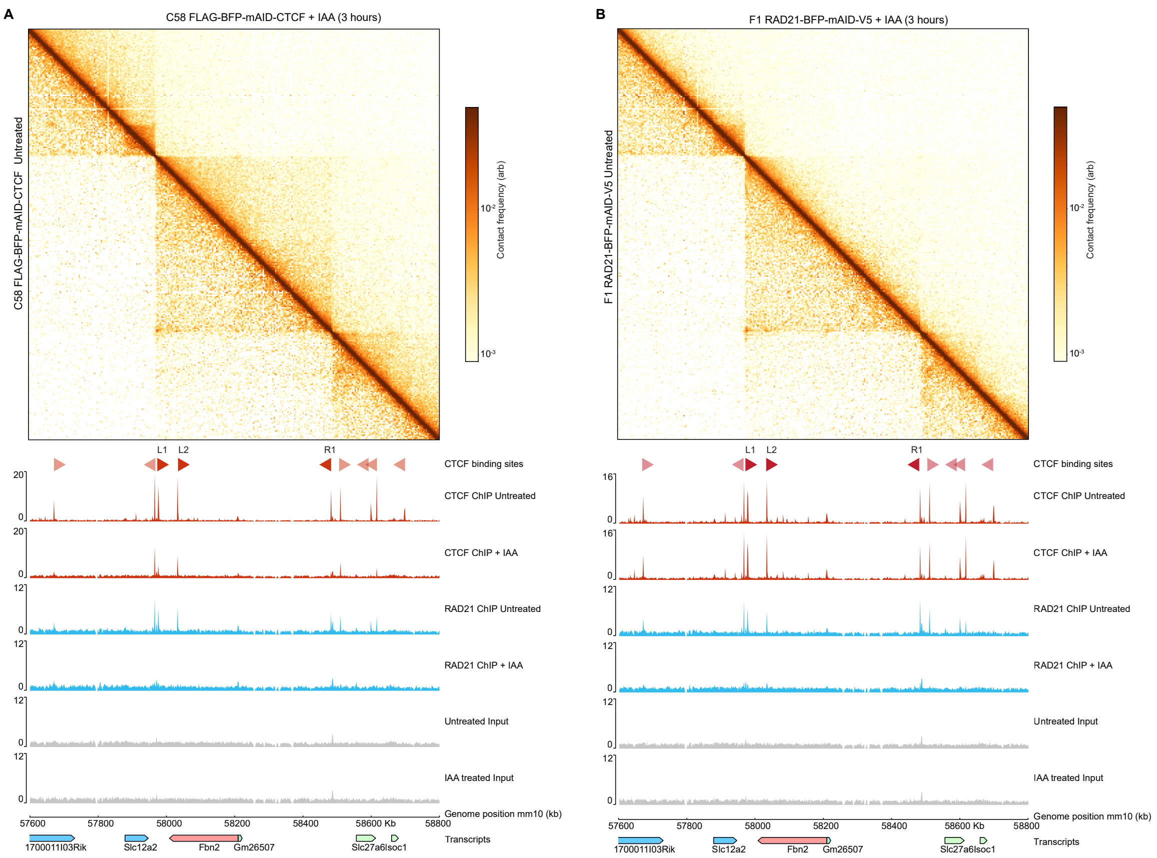

**Fig. S7: Acute depletion of CTCF and RAD21 leads to loss of CTCF and cohesin binding by ChIP-Seq and loss of *Fbn2* looping by Micro-C (A)** Contact map showing Micro-C data from C58 FLAG-BFP-mAID-CTCF untreated (bottom left, GSE178982) and C58 FLAG-BFP-mAID-CTCF following 3 hours of IAA treatment (top right, GSE178982), in the *Fbn2* locus, bin size = 4kb. Colorbar represent log normalized counts. Genome coordinates: mm10.Red triangles show all CTCF binding sites together with the orientation or polarity of the CTCF motif. At the bottom: ChIP-seq tracks of C58 FLAG-BFP-mAID-CTCF in treated and IAA treated (3 hours) conditions. CTCF IP untreated (GSM5402610), Rad21 IP untreated (GSM5402611), Input untreated (GSM5402609); CTCF IP+ IAA (GSM5402617), Rad21 IP + IAA (GSM5402618), Input + IAA (GSM5402616), transcripts (GRCm38). **(B)** Contact map showing Micro-C data from F1 RAD21-BFP-mAID-V5 untreated (bottom left, GSE178982) and F1 RAD21-BFP-mAID-V5 following 3 hours of IAA treatment (top right, GSE178982), in the *Fbn2* locus, bin size = 4kb. Colorbar represent log normalized counts. Genome coordinates: mm10. Red triangles show all CTCF binding sites together with the orientation or polarity of the CTCF motif. At the bottom: ChIP-seq performed in the line F1 RAD21-BFP-mAID-V5 with and without IAA treatment (3 hours). CTCF IP untreated (GSM5402622), Rad21 IP untreated (GSM5402623), Input untreated (GSM5402621); CTCF IP+ IAA (GSM5402629), Rad21 IP + IAA (GSM5402630), Input + IAA (GSM5402628), transcripts (GRCm38), transcripts (GRCm38).

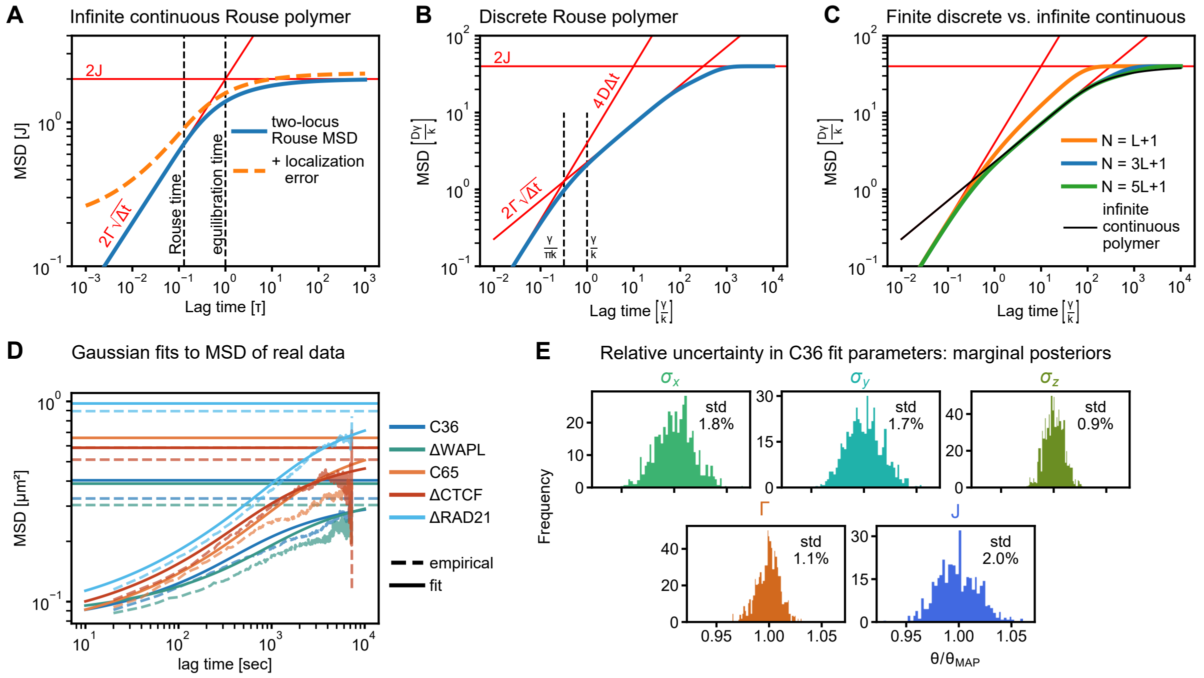

**Fig. S8: Visualization of analytical expressions for MSDs and fits to experimental data (A)** Numerical evaluation of eq. (32) showcasing asymptotic behavior and crossover. Note that the crossover region spans almost two orders of magnitude in time. **(B)** Visualization of the scaling results of section 6.3 (red), with exact (numerical) evaluation of eq. (64) for a chain of 61 monomers, using monomers 21 and 41 as tracer particles (blue). **(C)** Comparison of two-locus MSDs obtained from an infinite continuous chain, and discrete models of different (finite) length. All the discrete models have $L = 20$. **(D)** MAP MSD fits to our real data set, using the method of section 5. Horizontal lines are (twice) the steady state variance: $\mu(\infty) = 2\gamma(0) = 2\left\langle y^2 \right\rangle_c$. **(E)** Marginal posterior distributions over the individual parameters of the fit to C36 shown in **(D)**, evaluated by MCMC (n = 10,000 samples). Parameters are normalized to point estimate (MAP) value. ”std” gives standard deviation of the shown distribution.

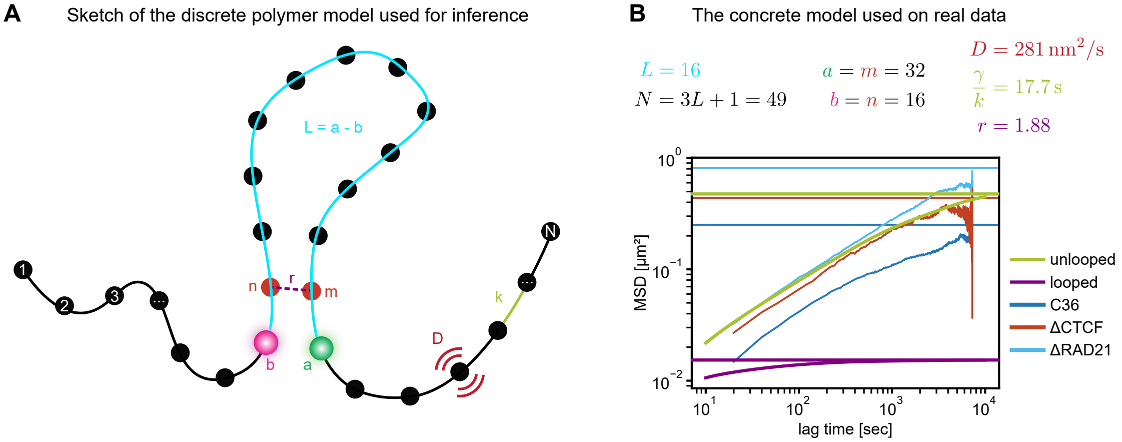

**Fig. S9: Overview over the model used for Bayesian Inference of Looping Dynamics (BILD) (A)** Cartoon of the two-state Rouse model used by BILD. Black circles represent monomers with individual diffusivity $D$, connected by springs with constant $k$. Tracer particles are at positions $a$ and $b$, bounding the "chain of interest" of length $L = a - b$ (wlog $a > b$). In the looped state an additional bond of relative strength $r$ is introduced between monomers $m$ and $n$. **(B)** Parameter values obtained by following the calibration scheme of section 7.2 (top). Bottom: MSDs of the looped and unlooped model states overlaid on top of experimental data. MSDs for the model states are obtained from eq. (64).

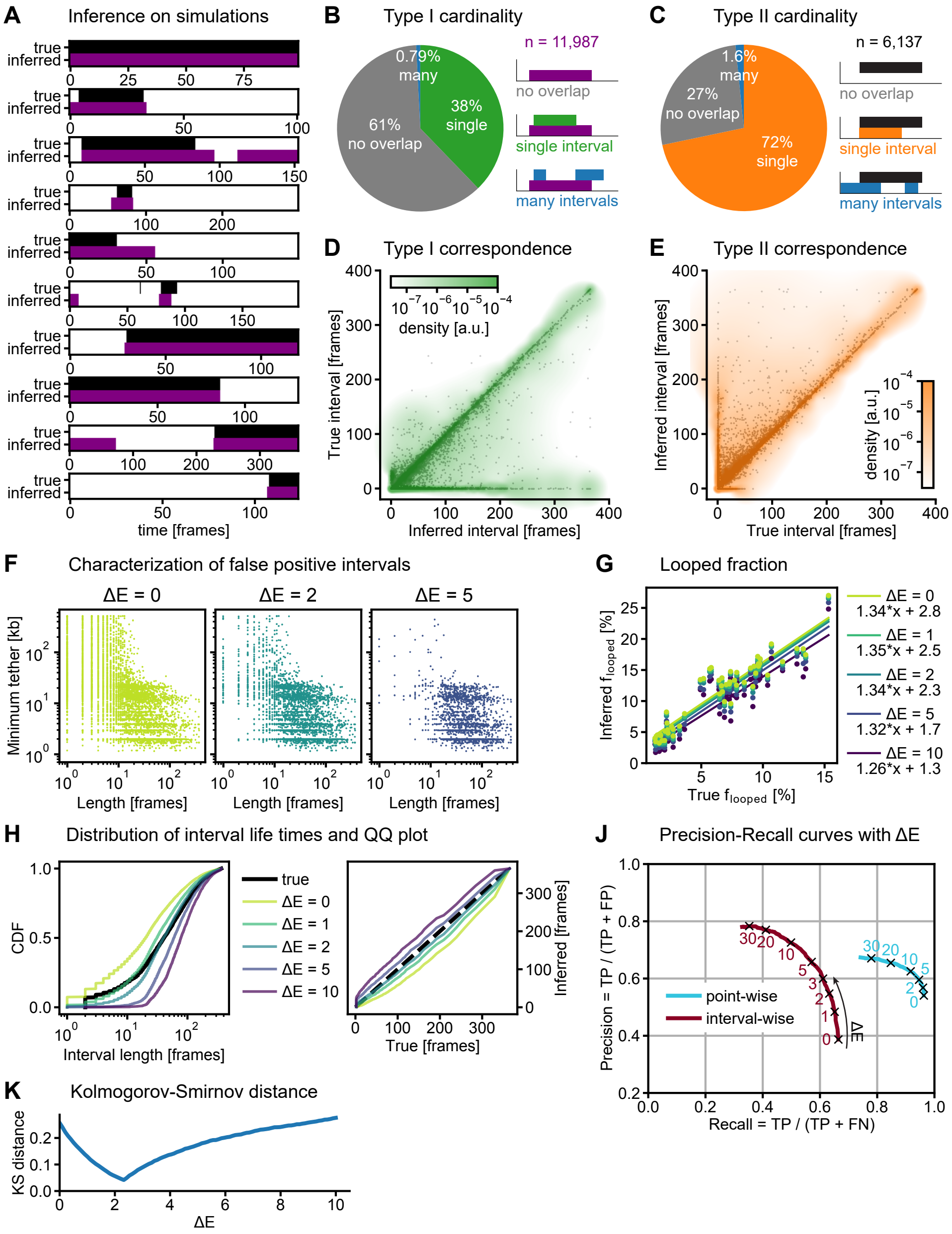

**Fig. S10: Validation of BILD on simulated trajectories (A)** Example inferences on randomly selected trajectories. Trajectories were randomly selected from all that had either at least 10 frames truly looped, or at least 10 frames inferred looped. **(B)** Counting true intervals overlapping each inferred interval. Counts higher than one are collectively labelled “many”. **(C)** Counting inferred intervals overlapping each true interval. Counts higher than one are collectively labelled “many”. **(D)** For each inferred interval, the length of the overlapping true interval, or zero if no true interval overlaps. In the rare cases where multiple intervals overlap, their length is summed. Green overlay is an adaptive Gaussian kernel density estimate, where each scatter point supports a Gaussian density with standard deviation equal to the distance to the tenth nearest neighbor or 2, whichever is greater. **(E)** Same as **(D)**, but applied to each true interval, showing the length of the overlapping inferred interval. **(F)** Scatter plots of false positive intervals (i.e. inferred intervals not overlapping any true interval) at different settings of $\Delta E$. The axes are length of the interval (horizontal) and minimum value of the ground truth effective tether length attained during the interval. **(G)** True vs. inferred looped fraction. Linear regression curves were fit by least squares. **(H)** Distribution of lengths of true and inferred intervals (left) and quantile-quantile plot (right). **(J)** Precision-Recall curves for the inference from point-wise and interval-wise points of view. See section 7.3 for details. **(K)** Kolmogorov-Smirnov distance (maximum difference between the cumulative distributions) of true and inferred lifetime distributions, at different $\Delta E$.

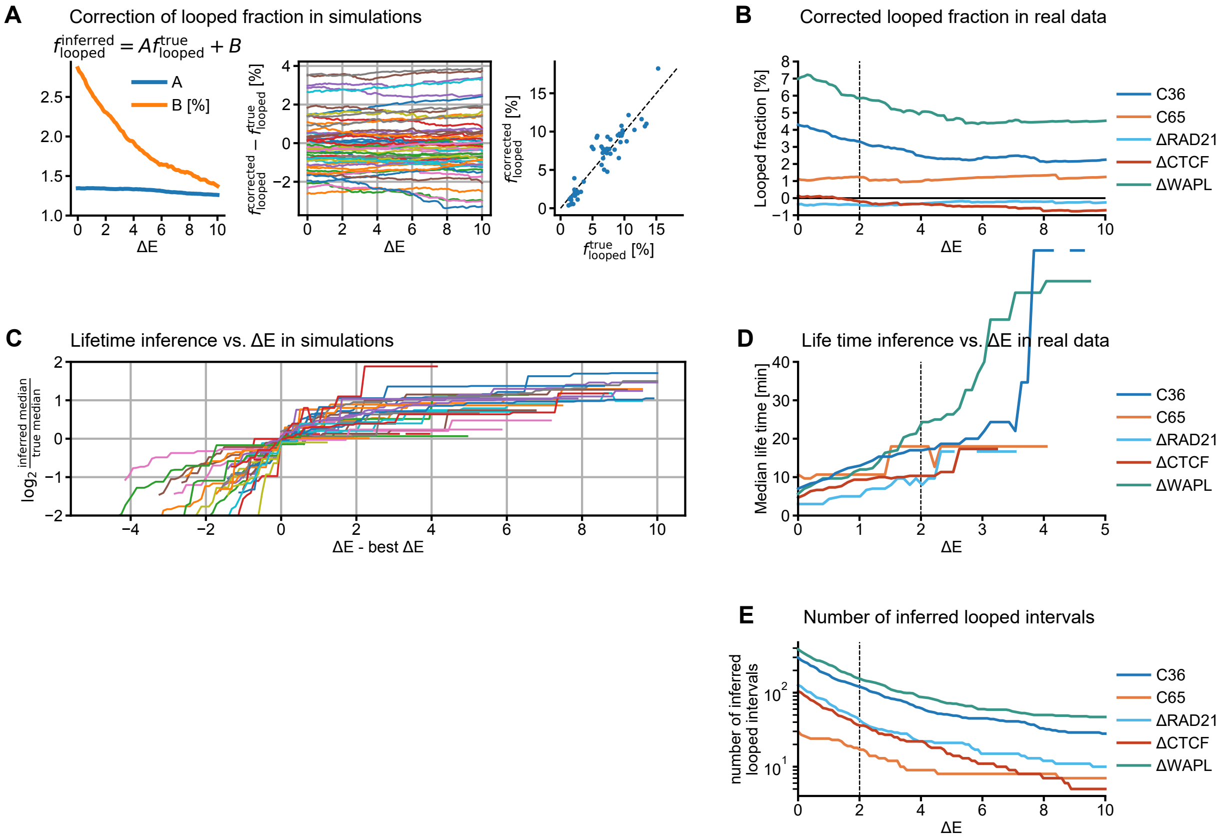

**Fig. S11: Variation of inference results with $\Delta E$ (A)** Left: quantifying the relationship between true and inferred looped fraction at different $\Delta E$. Compare **Fig. S10G**. Middle & Right: using the results displayed on the left to correct the inferred looped fractions gives stable values over a wide range of $\Delta E$, reproducing the ground truth to within two percentage points. "Corrected" values on the right are means over the curves shown in the middle, dashed line indicates identity. **(B)** Applying the same correction to our experimental data. **(C)** Inferred median lifetime vs. $\Delta E$ on all simulations in the validation data set. We normalize to the corresponding ground truth median lifetime, defined as the median of the Kaplan-Meier survival curve of the true intervals. Similarly, we match the horizontal coordinates to the $\Delta E$ value reproducing that true median lifetime for each simulation. **(D)** Variation of inferred median lifetimes with $\Delta E$ on the experimental data shown in Fig. 3. Note that for C65, ΔRAD21, and ΔCTCF there are very few intervals inferred as looped in the first place, such that the inferred lifetimes are not necessarily particularly meaningful (c.f. **(B)**, **(E)**). **(E)** Number of inferred looped intervals for each condition at different $\Delta E$.

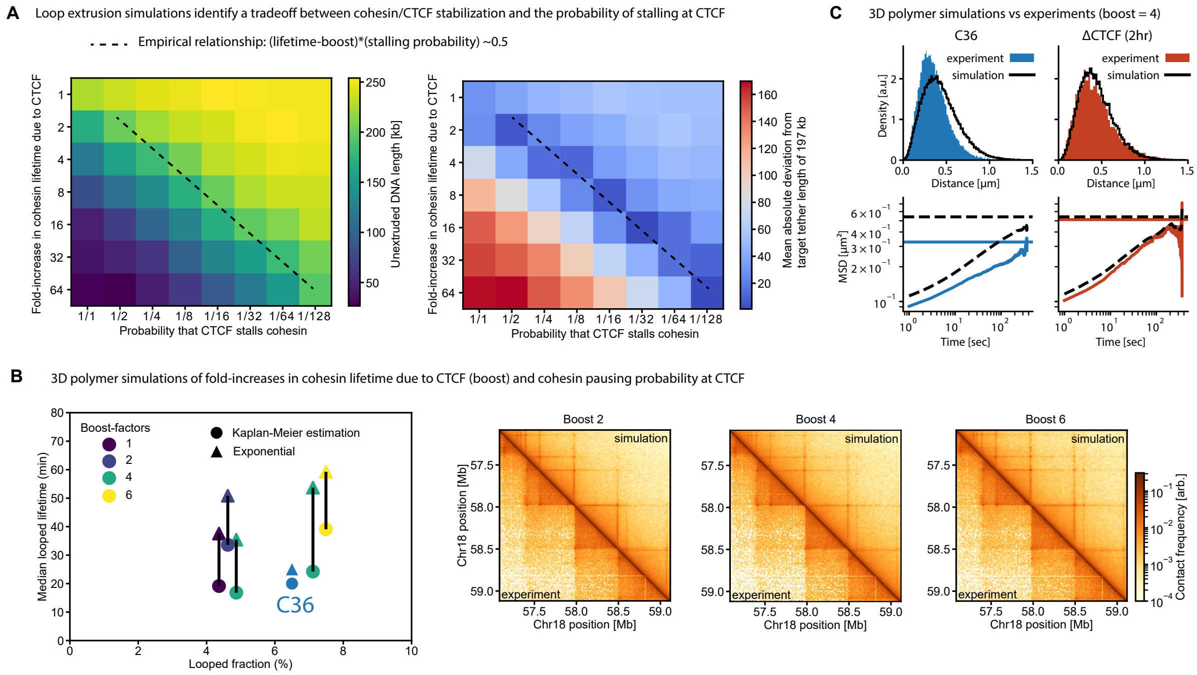

**Fig. S12: Additional loop extrusion and 3D polymer simulation results (A)** Relationship between the mean tether length (unextruded DNA length) to the probability of a CTCF stalling cohesin and the fold-increase in lifetime of cohesin due to stalling at CTCF. The mean tether length in C36 cells, identified from **Fig. 2D** and section 2.11, is $\approx$197 kb. This was used to find the empirical relationship between the fold-increase in cohesin lifetime and the stalling probability (black dashed line) **(B)** Simulation results for the loop lifetimes and looped fractions for simulations near the best-matching parameter values. For the boost factor 4 simulations, we show two independent repeats and the corresponding results for reference (left). **(C)** 3D probability density distributions and MSD curves comparing 3D polymer simulations to experimental data for the simulation with boost-factor=4 (and pausing probability = 1/8).

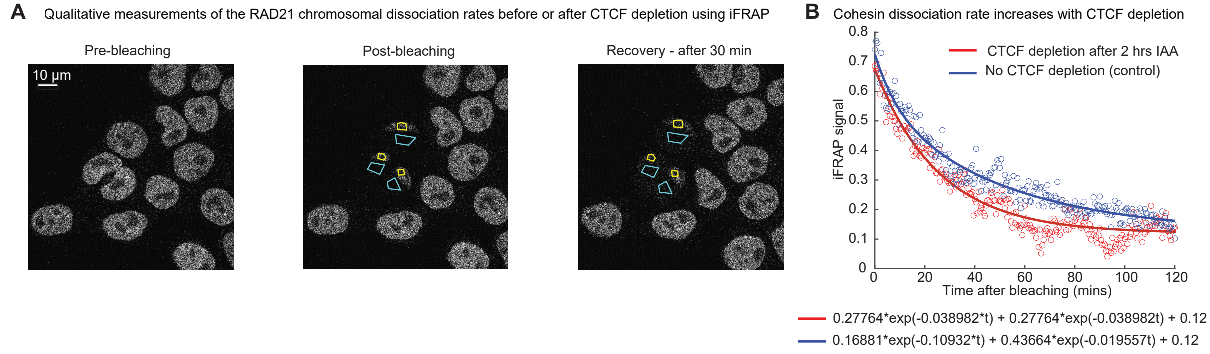

**Fig. S13: iFRAP image analysis of cohesin dissociation from the chromosome. (A)** Representative confocal images showing cells before (left) and after (right) bleaching the nucleus visualized by labeling R RAD21-$\text{SNAP}_{\mathrm{f}}$-V5 with 500 nM cp-SNAP-JF549 in the AID-CTCF cell line (clone C58). **(B)** Mean iFRAP signal decay and double exponential fits, showing that in the absence of CTCF depletion, the iFRAP curve of cohesin dissociation is best represented by a double exponential, whereas with 2 hours of CTCF depletion (500 μM IAA) the curve is best fit by a single exponential. The slow component of cohesin dissociation in the non-IAA treated cells is $\approx$5 times slower than the fast component in the non-IAA treated cells, and $\approx 2$ times slower than the dissociation rate for the IAA treated cells. The mean came from a total of n = 24 cells and n = 19 cells for the IAA and control conditions, respectively, where data was used from at least 8 different movies collected on at least three different days in each condition.

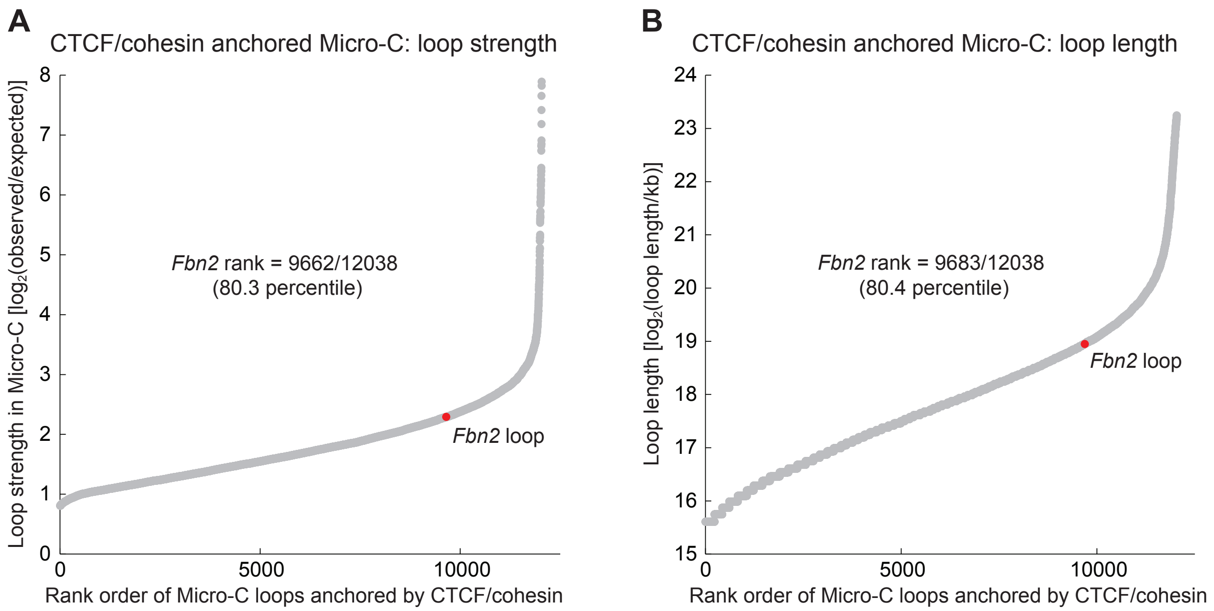

**Fig. S14: The *Fbn2* loop is somewhat stronger and longer than the average loop**. Using Micro-C data from Hsieh *et al.* [26], we used HiCCUPs [46] to call loops and focused our analysis on loops where both anchors are bound by both CTCF and cohesin yielding a total of 12,038 loops with sizes between 50 kb and 10 Mb. We then rank ordered these loops according to strength **(A)** and length **(B)** and plotted the position. Both when it comes to strength and length, the *Fbn2* loop is approximately at the 80th percentile, suggesting that it is in the strongest quartile of CTCF- and cohesin-mediated loops in mouse embryonic stem cells.

#### 11 Supplementary Tables

**Table S1:** List of plasmids used in this study

| Name | Purpose |
|---|---|
| pASH83 | Repair plasmid used to insert TetO array. 224x TetO array flanked by 131 homology against Fbn2 region. Also encodes PGK-driven HygroR |
| pASH84 | Cas9 Venus vector encoding sgRNA#1 5'-gttaactgagatctattcgc-3' targeting Fbn2 131 region for knocking in the TetO array. |
| pASH85 | Cas9 Venus vector encoding sgRNA#2 5'-ttaactgagatctattcgca-3' targeting Fbn2 131 region for knocking in the TetO array. |
| pASH86 | Cas9 Venus vector encoding sgRNA#3 5'-gatctattcgcagggaacta-3' targeting Fbn2 131 region for knocking in the TetO array. |
| pASH89 | Plasmid used to make Anchor3 knock-in. Anchor3 array flanked by 646 homology against Fbn2 region. Also encodes puromycin resistance. |
| pASH101 | Cas9 Venus vector encoding sgRNA#2 5'-TACCTTAGCACTGCCTCGTA-3' targeting Fbn2 chr18:58.646 region for knocking in the Anchor3 array. |
| pASH102 | Cas9 Venus vector encoding sgRNA#4 5'-AAGTTTTGTGAGTCCTTACG-3' targeting Fbn2 chr18:58.646 region for knocking in the Anchor3 array. |
| pASH103 | Cas9 Venus vector encoding sgRNA#8 5'-GTCCTTACGAGGCAGTGCTA-3' targeting Fbn2 chr18:58.646 region for knocking in the Anchor3 array. |
| pASH135 | PiggyBac L30 promoter driven TetR-NLS-3x-mScarlet-NLS-DEx4 (4x RNA destabilization element to lower the expression level). |
| pASH136 | PiggyBac L30 promoter driven EGFP-OR3-DEx4 (4x RNA destabilization element to lower the expression level). |
| pASH133 | AID vector expressing osTir1: PiggyBac pEF1a NES-osTir1-3xMyc-P2A-BSD (Blasticidin) |
| pASH107 | HR vector for N-terminal CTCF taggingwith FLAG-BFP-mAID-TEV-CTCF; Pair with sgRNAs pASH108-111. Targets FLAG-Halo-CTCF tagged endogenously. |
| pASH108 | Cas9 Venus vector encoding sgRNA #1N which targets Halo-mCTCF on the N-terminal side of Halo. Guide sequence: TCCACATAATGGGGGTCGAA |
| pASH109 | Cas9 Venus vector encoding sgRNA #3N which targets Halo-mCTCF on the N-terminal side of Halo. Guide sequence: TCCATGGCAGAAATCGGTAC |
| pASH110 | Cas9 Venus vector encoding sgRNA #4C which targets Halo-mCTCF on the C-terminal side of Halo. Guide sequence: TCGGCAGCGAGATCGCGCGC |
| pASH111 | Cas9 Venus vector encoding sgRNA #5C which targets Halo-mCTCF on the C-terminal side of Halo. Guide sequence: TTCCGGCGAGCCAACCACTG |
| pASH112 | HR vector for C-terminal RAD21 tagging with RAD21-GDGAGLIN-BFP-mAID-V5; Pair with sgRNAs pASH113-116. Targets mRad21-SNAP$_f$-V5 tagged endogenously. |
| pASH113 | Cas9 Venus vector encoding sgRNA #1N which targets mRad21-SNAP$_f$-V5 on the N-terminal side of SNAPf. Guide sequence: CACATCATTGGCGACGGCGC |
| pASH114 | Cas9 Venus vector encoding sgRNA #2N which targets mRad21-SNAP$_f$-V5 on the N-terminal side of SNAPf. Guide sequence: CGAAATGAAGCGCACCACCC |
| pASH115 | Cas9 Venus vector encoding sgRNA #1C which targets mRad21-SNAP$_f$-V5 on the C-terminal side of SNAPf. Guide sequence: GCGGGCTCGCCGTGAAAGAG |
| pASH116 | Cas9 Venus vector encoding sgRNA #2C which targets mRad21-SNAP$_f$-V5 on the C-terminal side of SNAPf. Guide sequence: CTGGGGCTCGACTCTACTTG |
| pASH117 | Homology repair vector for N-terminal tagging of mouse WAPL with HA-BFP-mAID-GDGAGLIN-mWAPL; Comes with the sgRNAs pASH118-120. Used to target wt-WAPL endogenously. |
| pASH118 | Cas9 Venus vector encoding sgRNA #1 which targets mouse WAPL for AID or other tagging. Guide sequence: ACTAAGAGTAGTCCGTTTGT |
| pASH119 | Cas9 Venus vector encoding sgRNA #2 which targets mouse WAPL for AID or other tagging. Guide sequence: GTCAAAATGACATCCAGATT |
| pASH120 | Cas9 Venus vector encoding sgRNA #3 which targets mouse WAPL for AID or other tagging. Guide sequence: GATTTGGAAAAACTTACAGT |
| pASH26 | Cas9 vector expressing sgRNA #71fw. For knocking out the L1 CTCF site in FBN2 loop, left side. TCCTGGGGTACAAATAGAAT TGG |
| pASH27 | Cas9 vector expressing sgRNA #43fw. For knocking out the L1 CTCF site in FBN2 loop, left side. ACACTGAGATTCTGTACAGC AGG |
| pASH28 | Cas9 vector expressing sgRNA #56fw. For knocking out the L1 CTCF site in FBN2 loop, left side. GTACAGCAGGACTTATCCTG GGG |
| pASH29 | Cas9 vector expressing sgRNA #325fw. For knocking out the L1 CTCF site in FBN2 loop, right side. GGCAGTATTGCTCCTGTCTT TGG |
| pASH30 | Cas9 vector expressing sgRNA #356rev. For knocking out the L1 CTCF site in FBN2 loop, right side. AGTGGACATTGCGCCAGACT CGG |
| pASH31 | Cas9 vector expressing sgRNA #385fw. For knocking out the L1 CTCF site in FBN2 loop, right side. GCGCAATGTCCACTCTACTG TGG |
| pASH18 | Cas9 vector expressing sgRNA #4. For knocking out the L2 CTCF site in FBN2 loop, left side. guide sequence: ATTGAACGTATCCTAAAGGC (PAM: TGG) on-target locus: chr18:+58193099 |
| pASH19 | Cas9 vector expressing sgRNA #12. For knocking out the L2 CTCF site in FBN2 loop, left side. guide sequence: CCTAAAGGCTGGTATGCTGA (PAM: AGG) on-target locus: chr18:+58193110 |
| pASH20 | Cas9 vector expressing sgRNA #9. For knocking out the L2 CTCF site in FBN2 loop, right side. guide sequence: GTGGCGAAACCTCAAGATG (PAM: GGG) on-target locus: chr18:+58193159 |
| pASH21 | Cas9 vector expressing sgRNA #5. For knocking out the L2 CTCF site in FBN2 loop, right side. guide sequence: GCATAGTAGGAGGGTGTC (PAM: AGG) on-target locus: chr18:+58193179 |
| pASH22 | Cas9 vector expressing sgRNA #10. For knocking out the R1 CTCF site in FBN2 loop, left side. guide sequence: GGCTACTCAAAAAAAAAAGG (PAM: GGG) on-target locus: chr18:+58641469 |
| pASH23 | Cas9 vector expressing sgRNA #8. For knocking out the R1 CTCF site in FBN2 loop, left side. guide sequence: CAGATACTGGTCATTTGAAA (PAM: TGG) on-target locus: chr18:+58641448 |
| pASH24 | Cas9 vector expressing sgRNA #1. For knocking out the R1 CTCF site in FBN2 loop, right side. guide sequence: AGTATGTTCAGCTTAGGGGT (PAM: AGG) on-target locus: chr18:+58641580 |
| pASH25 | Cas9 vector expressing sgRNA #4. For knocking out the R1 CTCF site in FBN2 loop, right side. guide sequence: TGCCCTCTAGTGGAAAAGTA (PAM: AGG) on-target locus: chr18:+58641522 |

**Table S2:** List of primers used in this study

| Name | Sequence | Purpose |
|---|---|---|
| ASH332_LYJ25_5'_131_OuterPrimer | AGGAAGAAGGGTTATATATGGCTAGTGTCTC | Primer for genotyping the 131 TetO Fbn2 array |
| ASH333_Gina2102_131_gPCR_KIN_REV | ATCGAAGCTTGGTCGACGATAGT | Primer for genotyping the 131 TetO Fbn2 array |
| ASH334_Gina2107_131_gPCR_WT_FWD1 | gggcattcggaaaggcactc | Primer for genotyping 131 TetO array, negative control for WT |
| ASH335_LYJ204_Tet_HygR_Check_O_Rev | aagagcgacaaacctgcctt | Primer for genotyping 131 TetO array, negative control for WT |
| ASH183_TetO_qPCR_F1 | GAGCGACCTTTCTCTATCAC | qPCR primer for testing TetO array integrity |
| ASH184_TetO_qPCR_R1 | GAACCAGAGATTCCCTATCA | qPCR primer for testing TetO array integrity |
| ASH185_TetO_qPCR_F8 | CACTGATAGGGACAGCAAG | qPCR primer for testing TetO array integrity |
| ASH186_TetO_qPCR_R8 | GAGAGTAGTAATGCCGAACC | qPCR primer for testing TetO array integrity |
| ASH187_131_gDNA_con_F1 | tgacaaggttttgtcatgtt | Control qPCR primer for testing TetO array integrity |
| ASH188_131_gDNA_con_R1 | gttttcctcttttgatctgc | Control qPCR primer for testing TetO array integrity |
| ASH189_131_gDNA_con_F9 | agcaagagaaacagaaagga | Control qPCR primer for testing TetO array integrity |
| ASH190_131_gDNA_con_R9 | ttctcctggggattataaga | Control qPCR primer for testing TetO array integrity |
| ASH172_Ginas2021 | GGATACGCTGCTTTAATGCCTTTG | Primers for genotyping the chr18:58.646 Anchor3 insertion |
| ASH173_Ginas2023 | CTCCAATTTTTCAAGGCCACATGG | Primers for genotyping the chr18:58.646 Anchor3 insertion |
| ASH174_Ginas2025 | CATGGGGACCCTGTGTAAGTG | Primers for genotyping the chr18:58.646 Anchor3 insertion |
| ASH175_Ginas2026 | CAGGTGGTGAACCCGGAATG | Primers for genotyping the chr18:58.646 Anchor3 insertion |
| ASH243_5'_646_Outer_F1 | cacagttgcagagtccaatctgttgt | Primers for genotyping the chr18:58.646 Anchor3 insertion |
| ASH244_646_gPCR_KIN_Anch3_R1 | GAAGATCCTGCCAAAGTGACGTC | Primers for genotyping the chr18:58.646 Anchor3 insertion |
| ASH66_mCTCF_genome_F1 | AGCACTGGTAGTTCTTTGTGGT | Primers for genotyping AID-tagging of CTCF |
| ASH67_mCTCF_genome_R1 | GTTGGCTTCGGAGGCTTCATA | Primers for genotyping AID-tagging of CTCF |
| 3DN2 | GGTGCCCTCGTAGGGTTG | Primers for genotyping AID-tagging of CTCF |
| 1332_Halo_REV | AACAGCAGCTTCGGGACAGG | Primers for genotyping AID-tagging of CTCF |
| 1331_Halo_FWD | ACCACGTCCGCTTCATGGAT | Primers for genotyping AID-tagging of CTCF |
| ASH97_GMD/gPCR_OUT_mRad21_fwd | ctggagcacccgtgacagttc | Primers for genotyping AID-tagging of RAD21 |
| ASH98_GMD/gPCR_OUT_mRad21_rev | CTGAGGAGTCACGCCACTGT | Primers for genotyping AID-tagging of RAD21 |
| ASH99 GMD/SNAP_fwd_END | ACATGAAGGACACCGCCTTGG | Primers for genotyping AID-tagging of RAD21 |
| ASH100 GMD/SNAP_rev_END | TCTCGTGCAGCCCTTGTTCG | Primers for genotyping AID-tagging of RAD21 |
| ASH101_mRad21_F1_genome | GAAGTGTCAAGCCAGGGGTA | Primers for genotyping AID-tagging of RAD21 |
| ASH102_mRad21_R1_genome | TGCGGTGCATAGCATCAGAG | Primers for genotyping AID-tagging of RAD21 |
| 2441_mWAPL_gPCR_OUTfwd | GGAGTTCTCCAACTTCAGAAAGATGT | Primers for genotyping AID-tagging of WAPL |
| 2442_mWAPL_gPCR_OUTrev | ccaaacactccatgcacacaag | Primers for genotyping AID-tagging of WAPL |
| ASH223_136_L1_gDNA_F1 | GTACAGCAGGACTTATCCTGGG | Primers for genotyping L1, L2, or R1 deletions |
| ASH224_136_L1_gDNA_R1 | AGTAGAGTGGACATTGCGCC | Primers for genotyping L1, L2, or R1 deletions |
| ASH225_136_L1_gDNA_F2 | AGCAGGACTTATCCTGGGGT | Primers for genotyping L1, L2, or R1 deletions |
| ASH226_136_L1_gDNA_R2 | ATTCCTCTGCTTTCCGTGGG | Primers for genotyping L1, L2, or R1 deletions |
| ASH227_136_L1_gDNA_F3 | AAAGTAGTTTCTGCTCGGCG | Primers for genotyping L1, L2, or R1 deletions |
| ASH228_136_L1_gDNA_R3 | TCCGTGGGACCCAGTAAGAA | Primers for genotyping L1, L2, or R1 deletions |
| ASH229_136_L1_deWit_F4 | AGCAGGACTTATCCTGGGGTAC | Primers for genotyping L1, L2, or R1 deletions |
| ASH230_136_L1_deWit_R4 | AAGGGGGCTCACGAGAAGTA | Primers for genotyping L1, L2, or R1 deletions |
| ASH231_192_L2_gDNA_F1 | GATTTGAGGAACGGGAGCGT | Primers for genotyping L1, L2, or R1 deletions |
| ASH232_192_L2_gDNA_R1 | CCATCTTGAGGTTTCGCCAC | Primers for genotyping L1, L2, or R1 deletions |
| ASH233_192_L2_gDNA_F2 | GGAACGGGAGCGTATGCTTT | Primers for genotyping L1, L2, or R1 deletions |
| ASH234_192_L2_gDNA_R2 | CCCCATCTTGAGGTTTCGCC | Primers for genotyping L1, L2, or R1 deletions |
| ASH235_192_L2_gDNA_F3 | ACACGGAGAGTGGTTTTGAGT | Primers for genotyping L1, L2, or R1 deletions |
| ASH236_192_L2_gDNA_R3 | GGGCCCATGCTTCGAAATTA | Primers for genotyping L1, L2, or R1 deletions |
| ASH237_641_R1_gDNA_F1 | TTCCTTCCCCAACATATGCCC | Primers for genotyping L1, L2, or R1 deletions |
| ASH238_641_R1_gDNA_R1 | GGCTGATCTGAGTAGACGACC | Primers for genotyping L1, L2, or R1 deletions |
| ASH239_641_R1_gDNA_F2 | TGTGATACCCACCCAAGTGC | Primers for genotyping L1, L2, or R1 deletions |
| ASH240_641_R1_gDNA_R2 | CCTACCCCTAAGCTGAACATACT | Primers for genotyping L1, L2, or R1 deletions |
| ASH241_641_R1_gDNA_F3 | CCCAACCATCATTGCTTGTCAG | Primers for genotyping L1, L2, or R1 deletions |
| ASH242_641_R1_gDNA_R3 | GGCTTCGAAGGAAGTCACAT | Primers for genotyping L1, L2, or R1 deletions |
| ASH281_CTCF_L1_long_F1 | AGAGCCTGCCAGTTGTTAGTG | Primers for genotyping L1, L2, or R1 deletions |
| ASH282_CTCF_L1_long_R1 | ATTGCACAGTGTATGCACCATT | Primers for genotyping L1, L2, or R1 deletions |
| ASH283_CTCF_L1_long_F2 | GACCCCATAGGGCTGCATCA | Primers for genotyping L1, L2, or R1 deletions |
| ASH284_CTCF_L1_long_R2 | CCCTTGTTTGTGAAATCTGGGTG | Primers for genotyping L1, L2, or R1 deletions |
| ASH285_CTCF_L2_long_F1 | CTGCATGCGCATCGTTCATA | Primers for genotyping L1, L2, or R1 deletions |
| ASH286_CTCF_L2_long_R1 | GGAAATAGCCGTGAACGTGG | Primers for genotyping L1, L2, or R1 deletions |
| ASH287_CTCF_L2_long_F2 | GCATGCGCATCGTTCATATTG | Primers for genotyping L1, L2, or R1 deletions |
| ASH288_CTCF_L2_long_R2 | CCGTGAACGTGGAGCAAAG | Primers for genotyping L1, L2, or R1 deletions |
| ASH289_CTCF_L2_long_F3 | TTGATGGGGGCTCTTCTATGTC | Primers for genotyping L1, L2, or R1 deletions |
| ASH290_CTCF_L2_long_R3 | AAGAGATACAGGGCATACAGCG | Primers for genotyping L1, L2, or R1 deletions |
| ASH291_CTCF_R1_long_F1 | CTCCCTGCTTTGAAAACTCGT | Primers for genotyping L1, L2, or R1 deletions |
| ASH292_CTCF_R1_long_R1 | TGGTCGACCTTGCCATTCAC | Primers for genotyping L1, L2, or R1 deletions |
| ASH293_CTCF_R1_long_F2 | TATTCTCTGCTCCCTGCTTTGAA | Primers for genotyping L1, L2, or R1 deletions |
| ASH294_CTCF_R1_long_R2 | GTAGGAACGCTCTACCGCC | Primers for genotyping L1, L2, or R1 deletions |
| ASH295_CTCF_R1_long_F3 | AAATGTTTCCTCCAGACCTTTGC | Primers for genotyping L1, L2, or R1 deletions |
| ASH296_CTCF_R1_long_R3 | AAAGAAGGCACGGGCAACTC | Primers for genotyping L1, L2, or R1 deletions |
| ASH310_L1_mid_F1 | CTTCTCCGTATCTCTCACGCC | Primers for genotyping L1, L2, or R1 deletions |
| ASH311_L1_mid_R1 | TAAGGGGGCTCACGAGAAGT | Primers for genotyping L1, L2, or R1 deletions |
| ASH312_L1_mid_F2 | CCGTATCTCTCACGCCACCT | Primers for genotyping L1, L2, or R1 deletions |
| ASH313_L1_mid_R2 | GCTTAGCACGCGATTCCTCA | Primers for genotyping L1, L2, or R1 deletions |

**Table S3:** Localization error for each imaging condition

| Condition | Cell line | Error on distance (nm) | Error per dot X (nm) | Error per dot Y (nm) | Error per dot Z (nm) |
|---|---|---|---|---|---|
| WT | C36 | 188 | 47 | 47 | 115 |
| Looping control | C27 | 196 | 52 | 50 | 118 |
| No CTCF control | C65 | 182 | 43 | 43 | 113 |
| AID-CTCF (control) | C58 | 192 | 47 | 45 | 119 |
| AID-CTCF (2 hr) | C58 | 182 | 40 | 41 | 115 |
| AID-CTCF (4 hr) | C58 | 187 | 41 | 43 | 119 |
| RAD21-AID (control) | F1 | 230 | 44 | 45 | 150 |
| RAD21-AID (2 hr) | F1 | 192 | 46 | 46 | 119 |
| RAD21-AID (4 hr) | F1 | 283 | 43 | 42 | 191 |
| AID-WAPL (control) | C40 | 181 | 45 | 42 | 113 |
| AID-WAPL (4 hr) | C40 | 192 | 43 | 41 | 122 |
| AID-WAPL (6 hr) | C40 | 193 | 46 | 43 | 121 |

**Table S4:** Imaging summary statistics*

| Condition | Cell line | Number of movies | Number of trajectories | Trajectories over 2 hr | Trajectories over 1.5 hr | Avg number frames | Avg traj. length (min) |
|---|---|---|---|---|---|---|---|
| WT | C36 | 78 | 491 | 32 | 105 | 94.0 | 56.4 |
| Looping control | C27 | 37 | 358 | 29 | 79 | 89.9 | 53.7 |
| No CTCF control | C65 | 21 | 147 | 7 | 28 | 92.3 | 52.2 |
| AID-CTCF (control) | C58 | 20 | 151 | 3 | 27 | 77.0 | 46.3 |
| AID-CTCF (2 hr) | C58 | 31 | 137 | 5 | 21 | 76.4 | 49.6 |
| AID-CTCF (4 hr) | C58 | 32 | 165 | 11 | 30 | 77.2 | 50.9 |
| RAD21-AID (control) | F1 | 34 | 183 | 14 | 52 | 96.2 | 62.4 |
| RAD21-AID (2 hr) | F1 | 92 | 537 | 36 | 151 | 84.5 | 59.0 |
| RAD21-AID (4 hr) | F1 | 21 | 48 | 1 | 6 | 45.5 | 42.4 |
| AID-WAPL (control) | C40 | 35 | 257 | 11 | 55 | 81.7 | 51.7 |
| AID-WAPL (4 hr) | C40 | 38 | 215 | 12 | 42 | 84.4 | 50.0 |
| AID-WAPL (6 hr) | C40 | 33 | 205 | 17 | 43 | 79.8 | 49.5 |

*Statistics after applying the CNN to identify and remove frames with likely poor localization

**Table S5:** List of software

| Name | Link | Source |
|---|---|---|
| connect_the_dots | https://github.com/ahansenlab/connect_the_dots | This paper |
| tracklib | https://github.com/SGrosse-Holz/tracklib | This paper |
| polychrom 0.1.0 | https://github.com/open2c/polychrom | [30] |
| nglview 3.0.3 | https://github.com/nglviewer/nglview | [47] |
| BioFormats | https://www.openmicroscopy.org/bio-formats/ | [48] |
| python-bioformats 4.0.5 | https://github.com/CellProfiler/python-bioformats | |
| pywavelet 1.1.1 | https://pywavelets.readthedocs.io/en/latest/ | [49] |
| tensorflow 2.5.0 | https://www.tensorflow.org/ | |
| keras 2.5.0 | https://keras.io/ | |
| numpy 1.19.5 | https://numpy.org/ | [16] |
| scipy 1.4.1 | https://www.scipy.org/ | [50] |
| pandas 1.3.4 | https://pandas.pydata.org/ | [51] |
| matplotlib 3.1.3 | https://matplotlib.org/ | [17] |
| trackpy 0.4 | http://soft-matter.github.io/trackpy/v0.5.0/introduction.html | [52] |
| javabridge 0.0.0 | https://pythonhosted.org/javabridge/ | |
| skimage 0.16.2 | https://github.com/scikit-image/scikit-image | [53] |
| snakemake 6.0.0 | https://snakemake.readthedocs.io/en/stable/ | [54] |

**Table S6:** List of datasets

| Cell line | Dataset | Series | Series with GSM | Source |
|---|---|---|---|---|
| C59 | RNA-seq | GSE123636 | GSE123636 | [55] |
| C59 | CTCF ChIP | GSE123636 | GSM3508478 | [55] |
| C59 | SMC1A ChIP | GSE123636 | GSM3508477 | [55] |
| C59 | Input ChIP | GSE123636 | GSM3508475 | [55] |
| C36 | Micro-C | GSE187487 | GSE187487 | this study |
| C59 | Micro-C | GSE123636 | GSM3508483 | [55] |
| C65 | CTCF ChIP | GSE187487 | GSM5668637 | this study |
| C65 | SMC1A ChIP | GSE187487 | GSM5668638 | this study |
| C65 | Input ChIP | GSE187487 | GSM5668635 | this study |
| C65 | IgG ChIP | GSE187487 | GSM5668636 | this study |
| C58 FLAG-BFP-mAID-CTCF | Micro-C - IAA | GSE178982 | GSE178982 | [25] |
| C58 FLAG-BFP-mAID-CTCF | Micro-C + IAA | GSE178982 | GSE178982 | [25] |
| C58 FLAG-BFP-mAID-CTCF | CTCF ChIP - IAA | GSE178982 | GSM5402610 | [25] |
| C58 FLAG-BFP-mAID-CTCF | RAD21 ChIP - IAA | GSE178982 | GSM5402611 | [25] |
| C58 FLAG-BFP-mAID-CTCF | Input - IAA | GSE178982 | GSM5402609 | [25] |
| C58 FLAG-BFP-mAID-CTCF | CTCF ChIP + IAA | GSE178982 | GSM5402617 | [25] |
| C58 FLAG-BFP-mAID-CTCF | RAD21 ChIP + IAA | GSE178982 | GSM5402618 | [25] |
| C58 FLAG-BFP-mAID-CTCF | Input + IAA | GSE178982 | GSM5402616 | [25] |
| F1 RAD21-BFP-mAID-V5 | Micro-C - IAA | GSE178982 | GSE178982 | [25] |
| F1 RAD21-BFP-mAID-V5 | Micro-C + IAA | GSE178982 | GSE178982 | [25] |
| F1 RAD21-BFP-mAID-V5 | CTCF ChIP - IAA | GSE178982 | GSM5402622 | [25] |
| F1 RAD21-BFP-mAID-V5 | RAD21 ChIP - IAA | GSE178982 | GSM5402623 | [25] |
| F1 RAD21-BFP-mAID-V5 | Input - IAA | GSE178982 | GSM5402621 | [25] |
| F1 RAD21-BFP-mAID-V5 | CTCF ChIP + IAA | GSE178982 | GSM5402629 | [25] |
| F1 RAD21-BFP-mAID-V5 | RAD21 ChIP + IAA | GSE178982 | GSM5402630 | [25] |
| F1 RAD21-BFP-mAID-V5 | Input + IAA | GSE178982 | GSM5402628 | [25] |
| C40 HA-BFP-mAID-WAPL | Micro-C - IAA | GSE178982 | GSE178982 | [25] |
| C40 HA-BFP-mAID-WAPL | Micro-C + IAA | GSE178982 | GSE178982 | [25] |
